# Biological recognition of mirror-image glycans

**DOI:** 10.64898/2026.07.18.739369

**Authors:** Chuanhao Peng, Simatsidk Haregu, Claire Yi-ling Yang, Shreyas Gupta, Michael Evgen, Hani Choksi, Vanessa Affe, Eric J. Carpenter, Nicholas Twells, Mei-Ting Lin, Ayodeji Kulepa, Carolina Ortiz-Cordero, Alexxandra Sosa-Guir, Jonathan Lefèbre, Szu Yun Wang, Sunhee Bae, Laura L. Kiessling, Todd L. Lowary, Sheng-Kai Wang, Ching-Ching Yu, Landon J. Edgar, Christoph Rademacher, Lori J. West, Lara K. Mahal, Matthew S. Macauley, Ratmir Derda

**Affiliations:** Department of Chemistry, University of Alberta, Edmonton, AB, T6G 2G2, Canada; Department of Pharmacology and Toxicology, University of Toronto, Toronto, ON, M5S 1A8, Canada; Department of Chemistry, National Tsing Hua University, Hsinchu 300044, Taiwan; Department of Chemistry, Massachusetts Institute of Technology, Cambridge, MA, USA; Department of Pharmaceutical Sciences, University of Vienna, Vienna 1090, Austria; Department of Microbiology, Immunology and Genetics, University of Vienna, Max F. Perutz Laboratories, Vienna 1030, Austria; Institute of Biological Chemistry, Academia Sinica, Nangang, Taipei 11529, Taiwan; Institute of Biochemical Sciences, National Taiwan University, Taipei 106, Taiwan; 9Department of Chemistry, The University of Toronto, Toronto, ON, M5S 1A8, Canada; Department of Immunology, The University of Toronto, Toronto, ON, M5S 1A8, Canada; Department of Medical Microbiology and Immunology, University of Alberta, Edmonton, AB, Canada; Department of Pediatrics, University of Alberta, Edmonton, AB, Canada; Department of Surgery, University of Alberta, Edmonton, AB, Canada

## Abstract

Recent synthesis of essential enzymes, such as DNA and RNA polymerases with opposite chirality, has boosted the feasibility of creating mirror-image life. Such life, if ever produced, will undoubtedly be coated by a dense display of glycans (glycoproteins, glycolipids, polysaccharides) built from enantiomers of common monosaccharides. Recognition of mirror-image glycans by extant glycan-binding proteins (GBPs) may be critical for colonization by or immune response to mirror life organisms. We evaluated recognition of enantiomers of common glycans by a diverse set of purified GBPs (plant and human derived), antibodies (including IgM from human plasma), mammalian cells (including immune cells), and organs in live animals. We found that GBP binding to enantiomers of naturally prevalent glycans is widespread. Notably, L-glucose and L-galactose interact with fucose-binding lectins, including DC-SIGN, a C-type lectin expressed on immune cells. These interactions can be inhibited by soluble “natural” glycan ligands and enantiomeric ones confirming specificity. Binding of L-glycans to diverse immune cell repertoires revealed preferences for specific glycan enantiomers. IgM antibodies from human serum showed donor-specific recognition of L-glycans. We propose that the recognition of L-glycans by extant GBPs arises from their co-evolution over millennia with the L-glycans that are present in the glycocalyx of many microorganisms.

## Main

All known life is homochiral. The ‘molecular mirror world’ refers to a hypothetical biosphere composed entirely of enantiomeric (mirror-image) versions of the biomolecules that underpin life. An analysis of the threats posed by mirror-image life forms^1,2^ argued that, if such organisms were created, many aspects of the immune response to mirror bacteria could be deficient, presenting a cataclysmic hazard for life on earth. A follow-up commentary^3^ expanded on the crucial roles of cell surface glycans in assessing these risks, noting that glycan-dependent interactions play key roles in the innate and adaptive immune responses of humans and lower organisms^4–9^. Glycans are present on the surfaces of every cell on earth across all kingdoms of life and are crucial for functions at both the cellular and organismal levels^10^. Their prominent display on cell surfaces underscores their essential roles in intercellular communication. For instance, glycan recognition by GBPs is linked to self*versus* non-self-discrimination by immune systems^10^. An accurate assessment of our preparedness for the arrival of mirror-image organisms, therefore, requires consideration not only of proteins and nucleic acids but also glycans.

Substantial evidence indicates the cell surface glycans of different organisms can contain D-, or L-carbohydrate building blocks^11^. The molecular recognition of these different enantiomeric epitopes has never been systematically evaluated, but prediction of such interactions combined with preliminary experimental assessment has been recently disclosed^12^. Here, we apply DNA-encoded glycan technologies to evaluate such interactions across a range of enantiomeric glycan pairs and diverse GBPs *in vitro* and *in vivo*.

Microorganisms across the three kingdoms of life employ both Land D-glycans to build glycans, including glycocalyces and glycosylated natural products. Of the four core pentoses, three – arabinose, xylose and lyxose – occur in both L and D forms as components of natural glycans. Enantiomeric pairs of deoxyhexoses are common constituents of microbial surface polysaccharides, including lipopolysaccharide (LPS). For example, the LPS of *Pseudomonas fluorescens* contains both Dand L-quinovose, and the LPS of *Pseudomonas aeruginosa* contains Dand L-fucose derivatives. Both enantiomers of rhamnose are present in the LPS of some *Pseudomonas* spp.^13^, *Helicobacter pylori*^14^ and *Thiobacillus ferrooxidans*^15^, and in the N-glycans of giant viruses^16^. For both 3,6and 2,6-deoxy-hexoses, 15 of the 16 plausible stereoisomers have been observed as components of glycans, glycolipids or natural products^11^. Among *bona fide* hexopyranoses, natural oligosaccharides and natural product have been reported to contain L-galactose^17,18^, L-mannose^19,20^, L-altrose^21^ and D-altrose^22–24^, as well as D-gulose^25^ and L-gulose^11,26–28^ ^29^. The natural prevalence of Land D-pairs of glycan building blocks suggests the existence of GBPs that can recognize them. For example, due to the prominent presence of L-fucose (6-de-oxy-L-galactose) in mammalian Nand O-glycoproteins and glycolipids, L-fucose-binding proteins are some of the most studied GBPs. Recognition of L-galactose by L-fucose-binding GBPs has been described, but the extent to which this plasticity applies to GBP-glycan recognition is not known.

Structural data describing proteins that bind enantiomers of common glycans are scarce: from 2723 structures in the PDB of proteins bound to fucose, only eight contain D-fucose (1ABF, 1APB, 2AAC, 3WH7, 4WUT, 7ABP, 3CAH, 3N31), and of 2050 PDB structures of proteins bound to galactose only seven contain L-galactose (4D52, 9W6Y, 6ZN1, 4UFC, 2BP6, 4D52, 7LK7). The only two PDB entries that contain proteins in complex with L-glucose (7XSH, 9W7A) correspond to unusual carbohydrate processing enzymes. There are no entries in PDB that contain L-mannose. More broadly, until recently^12^ no GBPs were reported to recognize L-mannose or L-glucose.

Glycans can vary in sequence, length, and branching, and examples of synthesis from enantiomeric building blocks are rare^30^. GBPs recognize epitopes that are present in extended oligosaccharide structures as well as at non-reducing termini. The latter can be investigated using multivalent presentations of monosaccharides^31^. Here, we employed a DNA-encoded multivalent display of enantiomeric pairs of Dand L-mannose, Dand L-glucose, Dand L-galactose, and D-and L-fucose to perform the first comprehensive analysis of the recognition of enantiomers pairs by extant GBPs. Encoding both the structure and density of the glycan and measuring Land D-pairs in the same solution made it possible to do controlled measurements of all pairs of enantiomers with purified GBPs as well as GBPs displayed on cells *ex vivo* and live animals *in vivo*.

## Results

### DNA-barcoded multivalent display of Land D-glycan monomers

To create a DNA-encoded multivalent display of glycans, we coupled monosaccharides modified by an anomeric aglycone linker bearing a terminal activated carboxylic acid to pVIII, the major coat protein of M13 bacteriophage (**Fig. 1a,b**). Controlling the reaction conditions enabled the introduction of 20–4,000 copies of each glycan onto a single M13 virion of 700 nm length, as confirmed by MALDI-TOF mass spectrometry (**Fig. 1c**). When only 20 out of 2700 pVIII proteins were modified, the median spacing between glycans was greater than 30 nm (**Fig. 1d**).

**Figure 1.**
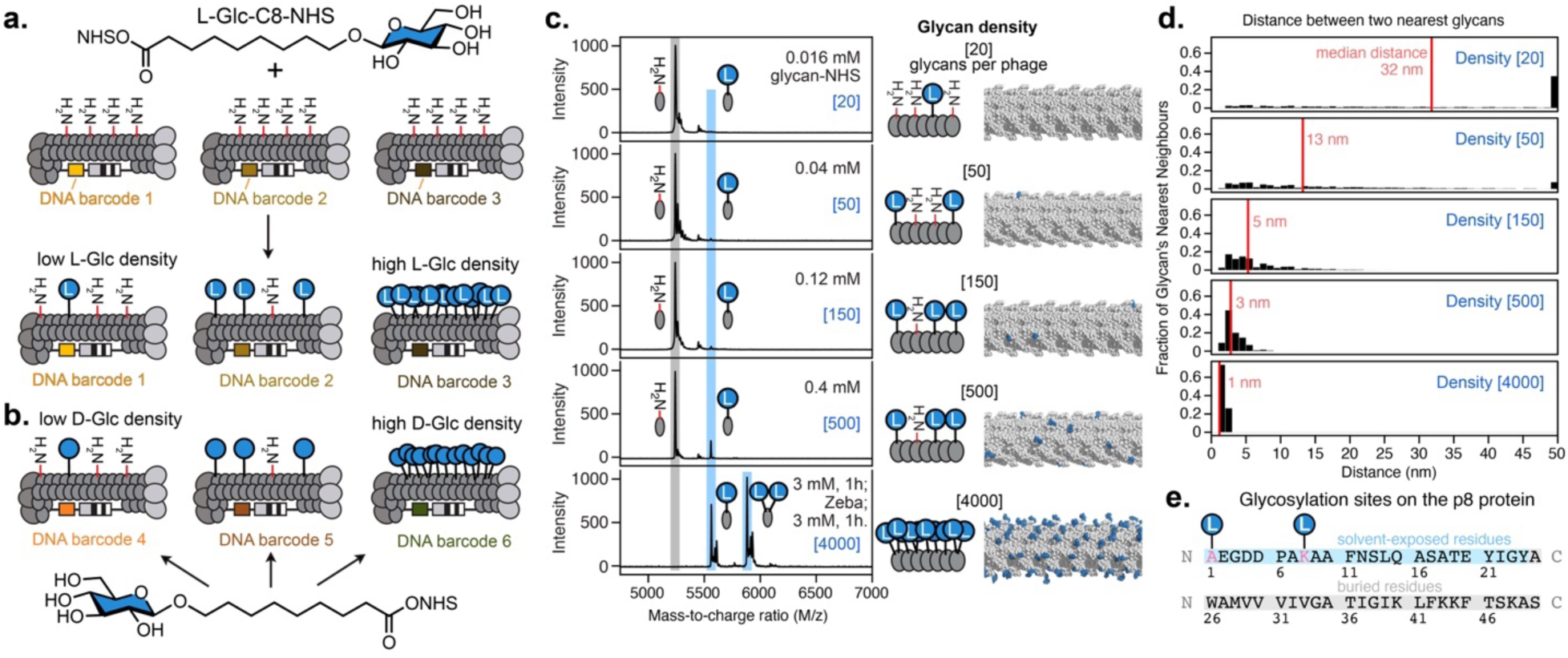
Synthesis of mirror-LiGA: a DNA-encoded display of a glycan and its mirror image at controlled display densities on M13 bacteriophage. **(a)** Amide-bond ligation of carboxy-glycans or **(b)** their enantiomers. **(c)** MALDI MS analysis of glycan conjugated to phage at a defined density. Rendering of phage displaying 20–4,000 copies of _L_-Glc monomer: a model built from a cryo-EM structure of M13. **(d)** Distribution of the proximity of glycan epitopes on phage displaying specific copy numbers of glycans. **(e)** N-terminus of Ala1 and side chains of Lys8 on pVIII protein modified by glycans. Density, or copies of glycans per virion, is represented as “[x]”.

When the phage was modified with 4,000 glycans, nearly every pVIII protein displayed one or two copies of the glycan (**Fig. 1b**), with both the N-terminus and Lys8 side-chain acylated (**Fig. 1b**), which corresponds to a median spacing of 1 nm (**Fig. 1d**).

To perform systematic analysis of structure and density, we prepared six discreet display densities of 20, 50, 150, 500, and 4,000 copies per phage virion for Dand L-mannose, Dand L-glucose, Dand L-galactose, and Dand L-fucose. Each unique glycan-density combination was linked to five distinct DNA-barcodes. The resulting mixture of glycophage constructs, termed “mirror-LiGA”, were tested alone or in combination with a previously reported Liquid Glycan Array (LiGA) containing ∼80 structures commonly observed in Nand O-linked glycans.

### Natural lectin specificity extends to rare mirror-image glycans

We calibrated the tested LiGA using 37 purified glycan-binding proteins with well-understood glycan recognition profiles. Binding of LiGA to plant and human GBPs followed canonical recognition of these lectins (**Fig. 2a,b** and **Extended Data Fig. 1**). In the same solution, the L-and Dmonomers exhibited canonical binding, such as D-Man binding to mannose-binding lectins (ConA, Griffithsin, BanLec H84T, SPD, and MBL), L-Fuc binding to fucose binding lectins (AAL, AOL, UEA-I, and RSL) and D-Gal binding to galactose-binding lectins (SBA, ECL, and PNA). Mirror LiGA confirmed previously reported interactions between L-Glc and UEA-I^12^ and L-Gal to purified DC-SIGN^32^. Novel observations were the recognition of L-Gal and L-Glc by the human lectin DC-SIGN (canonical D-Man / L-Fuc specificity) and plant AAL, AOL, and UEA-I, which are all lectins that recognize L-Fuc (**Fig. 2b**) plus the recognition of L-Glc by RSL, another L-Fuc binder. Recognition of L-Gal and L-Glc by these proteins was dependent on the anomeric configuration and absolute configuration. The presence of multiple density-matched stereoisomeric controls in mirror-LiGA offered rigorous tests for the validity of the observed interactions. For example, at the highest display density, neither D-Gal nor L-Man exhibited any binding to fucose-binding or mannose-binding lectins, ruling out non-specific interactions at high densities. Conversely, galactose-binding lectins engaged D-Gal, but not enantiomeric L-Gal, nor diastereomers D-Man, L-Man, D-Glc, or L-Glc. The same D-Gal-binding lectins bound to medium display densities of α-D-Fuc or high display density of β-D-Fuc but exhibited no detectable binding to any display density of any anomers of L-Fuc.

**Figure 2.**
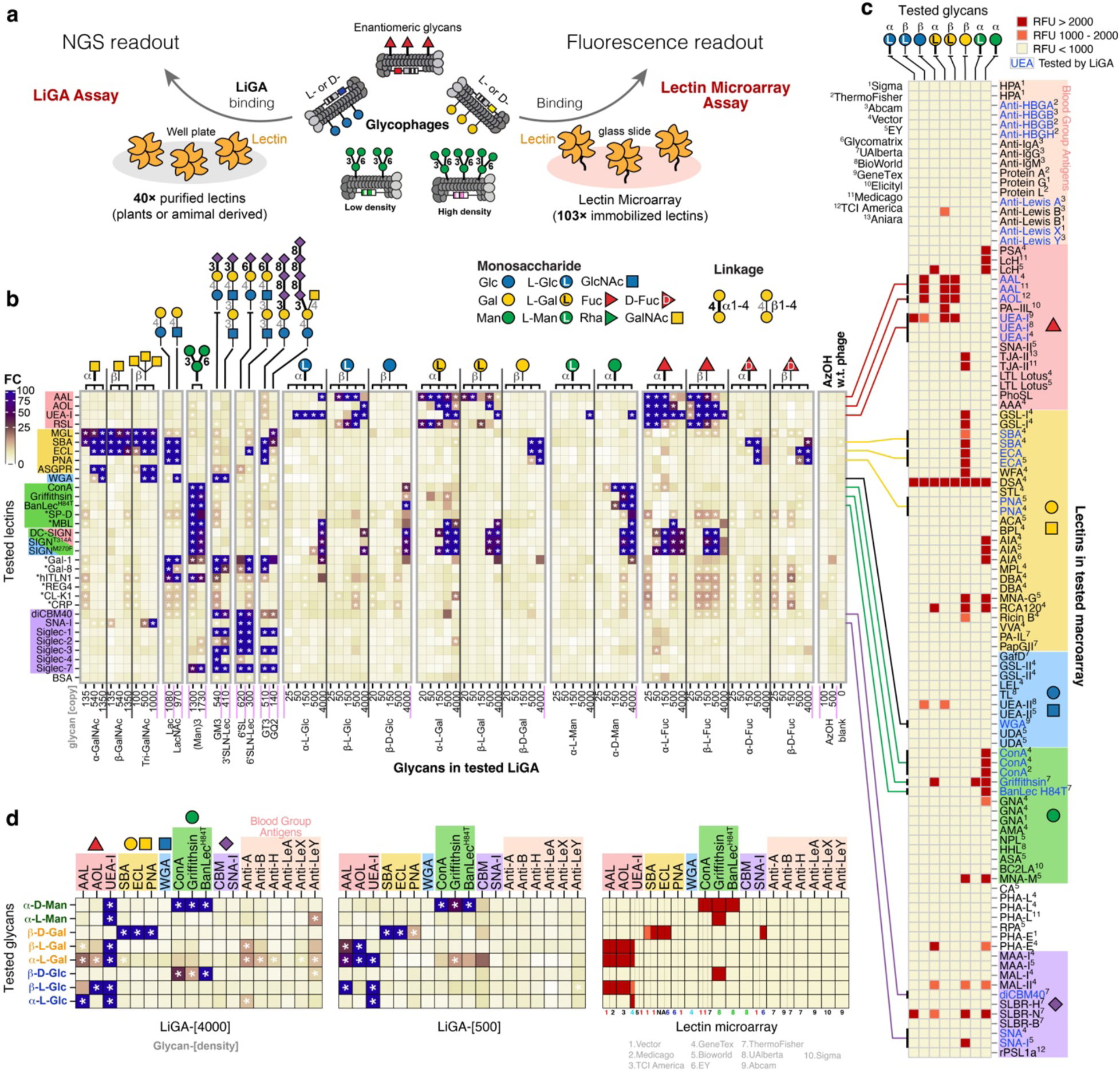
Interactions between enantiomeric glycans and purified GBPs. **(a)** Scheme of measuring binding of enantiomeric glycans to diverse plantor animalderived lectins using LiGA and lectin microarray assays. Glycosylated phages (glycophages) were labelled with a fluorophore before hybridization to lectin microarrays. **(b)** Heat map summary of binding between 37 GBPs, 15 control glycans, eight hexose stereoisomers (α_-L_-Glc, β_-L_-Glc, β_-D_-Glc, α_-L_-Gal, β_-L_-Gal, β_-D_-Gal, α_-L_-Man, and α_-D_-Man) and four fucose stereoisomers (α_-L_-Fuc, β_-L_-Fuc, α_-D_-Fuc, and β_-D_-Fuc). The data shows expected mimicry of galactose and fucose and unexpected similarity of glycan–GBP recognition between _L_-fucose and _L_-glucose. Among the 31 proteins shown and 37 proteins tested (see SI), only UEA-I exhibited binding to a dense display of _L_-Man. * indicates false-discovery rate (FDR) < 0.05. **(c)** Heat map summary of lectin-binding profile of eight hexose stereoisomers (α_-L_-Glc, β_-L_-Glc, β_-D_-Glc, α_-L_-Gal, β_-L_-Gal, β_-D_-Gal, α_-L_-Man, and α_-D_-Man) measured using lectin microarray. **(d)** Glycan binding comparison between LiGA and lectin microarray assay.

We found that both α and β-L-Gal bind to DC-SIGN, with the α-anomer binding at lower displayed density, whereas the enantiomer β-D-Gal or diastereomers α-L-Man had no measurable interactions with DC-SIGN (**Fig. 2b**). Recognition of L-Gal-containing glycans by DC-SIGN has been confirmed by Jimenez-Barbero and coworkers in their recent NMR spectroscopic study^32^. In the same solution, L-Fuc but not D-Fuc bind to DC-SIGN and, again, the α-anomer exhibited binding at as few as 50 copies per virion (13 nm median spacing). We found that even single mutations in these immune lectins broadened the recognition profile and enhanced recognition of mirror glycans. Based on prior work, the M270F mutant is expected to have higher affinity, and the T314A lower affinity than wild-type DC-SIGN. Both M270F and T314A variants bias the conformational equilibrium of an allosteric cryptic pocket in the carbohydrate-recognition domain of DC-SIGN^33^; these conformational changes transmit from a cryptic binding site to a canonical carbohydrate recognition domain. In accordance with the expectation, the M270F mutant of DC-SIGN engaged with α-L-Glc when displayed on M13 phage at 500 copies per virion whereas wild type DC-SIGN showed no binding to α-L-Glc until the modification reached 4,000 copies. In contrast, T314A mutant of DC-SIGN diminished its binding to beta-L-Glc, to 150 copies of α-L-Gal and α-L-Fuc, in accordance with the predicted decrease in affinity. Interestingly, tested in a set of 100 Nand O-linked glycan oligomers, none of these mutants changed the canonical recognition profile of DC-SIGN (**Extended Data Fig. 1**).

None of the monosaccharides, regardless of absolute stereochemistry were recognized by sialic-acid binding proteins (SNA and Siglecs) or proteins that recognize C2 *N*-acetyl-hexoses (MGL, ASGPR). No binding was observed for Intelectin-1, PTX3, REG4, CL-K1, and CRP1, which recognize microbial glycans^34,35^. All tested galectin proteins (Gal-1, Gal-2, Gal-3, Gal-7, Gal-8, Gal-10, and Gal-14) exhibited no to weak binding to glycan monomers (**Extended Data Fig. 1**).

Hybridization of glycosylated M13 phages modified by Land D-monosaccharides to slide-printed lectin microarrays provided complimentary information for 100 diverse glycan binding proteins (**Fig. 2c**). Every Land D-pair has a distinct lectin recognition profile but there were similarities in recognition by L-Glc to recognition by L-Fuc and L-Gal. We did note minor discrepancies between the binding of glycosylated clonal phages to the lectin microarray and the binding of the LiGA to individual lectins (**Fig. 2d**). For example, from 45 proteins tested by LiGA, only UEA-I exhibited modest recognition of the high-density L-Man, but this binding was not detected when a high-density L-Man display on phage was hybridized to the lectin array. Conversely, the Griffithsin lectin exhibited binding to L-Man in the microarray (**Fig. 2c, d**) but we could not detect any L-Man–Griffithsin interaction at any density in the LiGA (**Fig. 2b**).

Combined data from the LiGA and the lectin arrays suggested that L-Man was recognized the least by the tested lectins. From 100 lectins on the lectin array, DSA recognized L-Man but also 10 out of 13 tested monosaccharides (**Fig. 2c**) suggesting a poly-specificity rather than unique recognition of L-Man. Griffithsin and UEA-I emerged as plausible candidates, but their binding could not be reliably confirmed. L-Man was not recognized by classical mannose-binding proteins ConA, H84T BanLec, SPD, and MBL. These proteins exhibited binding to a high-valency D-Glc but not L-Glc. Both Land Dconfigurations of Man and Glc share a 3,4-bis-equatorial arrangement of hydroxyl groups forming the Ca^2+^-chelator, which exists in all D-Man binding proteins; however, the stereochemistry or C2 and C5 significantly impacts the recognition. Dedicated recognition of mannose represents a unique task for a protein interface and the search for extant GBP that binds to L-Man is an open challenge.

We previously demonstrated that the LiGA can measure the binding of glycans to GBPs on live cells^36,37^. Here, we employed this approach to confirm that recognition of L-glycans persists when GBPs are expressed on the cell surface. We confirmed that the L-Fuc/D-Man-binding lectins DC-SIGN (**Fig. 3** and **Extended Data Fig. 2a**) and Langerin (**Extended Data Fig. 2b**) recognize αand β-L-Gal but not β-D-Gal, as well as α-L-Glc and β-D-Glc but not β-L-Glc. We hypothesized that the non-canonical recognition of L-glycans by extant GBPs represents plasticity of the carbohydrate-recognition pocket. Under this model, interactions between GBPs and L-glycans should be competitively displaced by canonical D-glycan ligands. To test this hypothesis, we measured the binding of mirror-LiGA to live GBP-expressing cells in the presence of increasing concentrations of soluble D-Man (**Fig. 3**). Addition of soluble D-Man to DC-SIGN expressing Raji cells produced dose-dependent inhibition of glycophage binding, including both canonical DC-SIGN ligands and mirror-image glycans. Lewis A, Lewis X, (Man)_3_, α-D-Man, α-L-Gal, and α-L-Glc data fitted *IC_50_* values of 36, 25, 20, 21, 44, and <1 mM D-Man, respectively (**Fig. 3b**). Consistent with the observed *IC_50_*, van Liempt and coworkers have reported multivalent neoglycoconjugates displaying Lewis A showed greater affinity to DC-SIGN than similar neoglycoconjugates displaying Lewis X or (Man)_3_^38^. In the LiGA inhibition assay, the steep inhibition profiles observed for several glycophages are consistent with avidity effects arising from multivalent glycan display on the M13 virion and the multivalent presentation of DC-SIGN at the cell surface (**Fig. 3b**). As expected from the avidity-dependent interaction, glycophages displaying higher glycan densities required higher concentrations of soluble inhibitor to suppress binding (**Fig. 3** and **Extended Data Fig. 3**). Competition by soluble D-Man further supports the conclusion that recognition of L-Glc, L-Gal, and L-Fuc by DC-SIGN involves the canonical D-Man-binding site. Notably, α-L-Gal-conjugated glycophages were more resistant to D-Man competition than the canonical ligands shown, whereas α-L-Glc-conjugated glycophages were inhibited at sub-millimolar D-Man concentrations. In a converse competition experiment, soluble L-Glc also dose-dependently inhibited the binding of purified DC-SIGN to LiGA (**Extended Data Fig. 3b,d**), further supporting direct competition between canonical and mirror-image glycans for DC-SIGN recognition.

**Figure 3.**
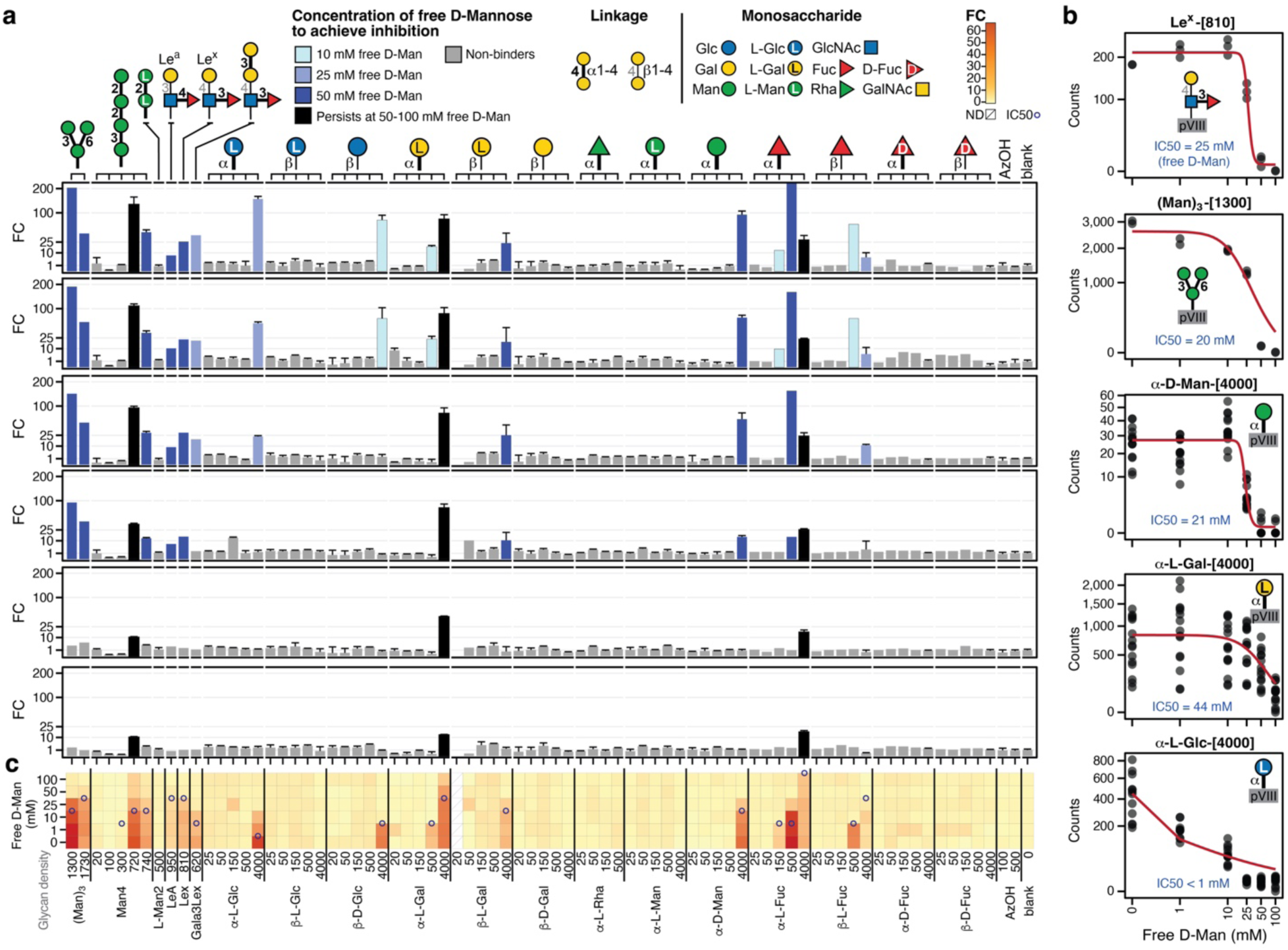
Soluble _D_-Man inhibits binding of mirror-image glycans to DC-SIGN on cells. **(a)** Binding of DC-SIGN^+^ Raji cells to mirror-LiGA in the presence of variable concentrations of soluble _D_-Man. Bars represent the median fold enrichment, and upper error bars indicate the 75th percentile across replicates. **(b)** Dose-dependent inhibition by soluble _D_-Man of Lewis X-, (Man)_3_-, α-_D_-Man-, α-L-Gal-, and α-L-Glc-conjugated glycophage binding to DC-SIGN^+^ Raji cells. **(c)** Heatmap summary for panel **(a)**. The concentration where inhibition reaches 50% (*IC_50_*) for each glycan-phage conjugates is indicated as ○ on the heatmap. In the bar charts, color is employed to highlight examples of strong, medium, and weak binders. The relative binding strengths of all glycan-phage conjugates to DC-SIGN can be estimated by comparing the concentrations of soluble _D_-Man required to achieve 50% inhibition. For glycans with the same stereochemistry at the secondary stereocenters, like _L_-Gal and _L_-Fuc, the higher display density per phage is a stronger binder whereas lower density of the same glycan produces a weak binder.

Just as in the inhibition experiment on DC-SIGN expressing Raji cells, soluble D-Man effectively inhibited binding of LiGA to purified DC-SIGN protein (**Extended Data Fig. 3a, b**). As little as 1 mM of D-Man was required to disrupt binding of purified DC-SIGN to β-D-Glc and α-L-Glc both at 500 glycans per virion. Inhibition at 500 copies per phage of α-D-Man, β-L-Gal, and α-L-Glc as well as β-L-Glc at 4,000 copies per phage occurred at 10–25 mM whereas interaction between DC-SIGN with α-L-Gal at 4,000 copies per phage persisted even at 100 mM and required 1,000 mM for inhibition. The extent of inhibition varied across the library, with several high-density L-monosaccharides displays retaining binding at intermediate D-Man concentrations (α-L-Glc, α-L-Gal, β-L-Gal, α-D-Man, and β-L-Fuc (all at 4,000 copies per phage) as well as α-L-Fuc at both 500 and 4,000 copies per phage).

As soluble D-Man disrupted binding of L-Glc to DC-SIGN, likewise soluble L-Glc inhibited binding of purified DC-SIGN to all canonical epitopes. Higher concentrations of L-Glc were generally required to achieve inhibition comparable to D-Man (**Extended Data Fig. 3b**). Binding was largely preserved at 1–2 mM L-Glc, reduced at 5–10 mM and strongly suppressed at 25–100 mM, with little residual binding at 500 mM. Notably, L-Glc inhibited DC-SIGN binding to both Dand L-configured glycans, including α-D-Man, β-L-Glc, α-L-Gal and β-L-Fuc conjugates. Dose-response analysis of four representative ligands yielded L-Glc *IC*_50_ values of 6 mM for α-L-Gal at 150 glycans per virion, 9 mM for α-D-Man at 500 glycans, 10 mM for β-L-Fuc at 150 glycans and 11 mM for β-L-Glc at 4,000 glycans (**Extended Data Fig. 3d**). Thus, L-Glc in solution inhibited DC-SIGN recognition of structurally and stereochemically distinct glycans with potency in a range similar to the physiological concentrations of blood glucose.

### Human peripheral immune cells recognize mirror-image glycans

The recognition of diverse mirror glycans by DC-SIGN prompted us to examine whether Land D-configured monosaccharides are also recognized by human peripheral blood mononuclear cells (PBMCs). Mirror-LiGA was incubated with PBMCs from four donors, after which monocytes, B cells, natural killer (NK) cells, and T cell subsets were isolated and subjected to sequencing-based glycan-binding profiling. CD4^+^ and CD8^+^ T cells were additionally analysed as pre-isolated populations without fluorescence-activated cell sorting (FACS).

In agreement with previous observations^39^, sialidase treatment of B cells promoted their binding to α-(2➞6)-linked sialylated glycans admixed to mirror-LiGA as positive controls (**Fig. 4**). This response is consistent with enzymatic removal of endogenous α-(2➞6)-linked sialylated cis ligands that mask Siglec-2 (CD22) on the B cell surface, thereby increasing the accessibility of Sig-lec-2 to exogenous α2,6-sialylated ligands. In contrast, binding to most monosaccharide conjugates was modest relative to the sialidase-enhanced recognition of the sialylated controls. Nevertheless, mirror-LiGA revealed reproducible and cell-type-dependent recognition of L-glycans. L- /D-Glc-modified phages produced the broadest overall binding profiles, particularly among the T cell subsets, whereas NK cells and monocytes showed comparatively sparse and heterogeneous recognition. B cells displayed binding to β-D-Glc (4,000 copies per virion) and α-L-Fuc in addition to a strong sialidase-dependent response to the α-(2➞6)-linked sialylated controls. Across the library, recognition varied with monosaccharide identity, stereochemistry, anomeric configuration and glycan display density, indicating that the observed interactions were not explained by nonspecific binding to the phage scaffold.

**Figure 4.**
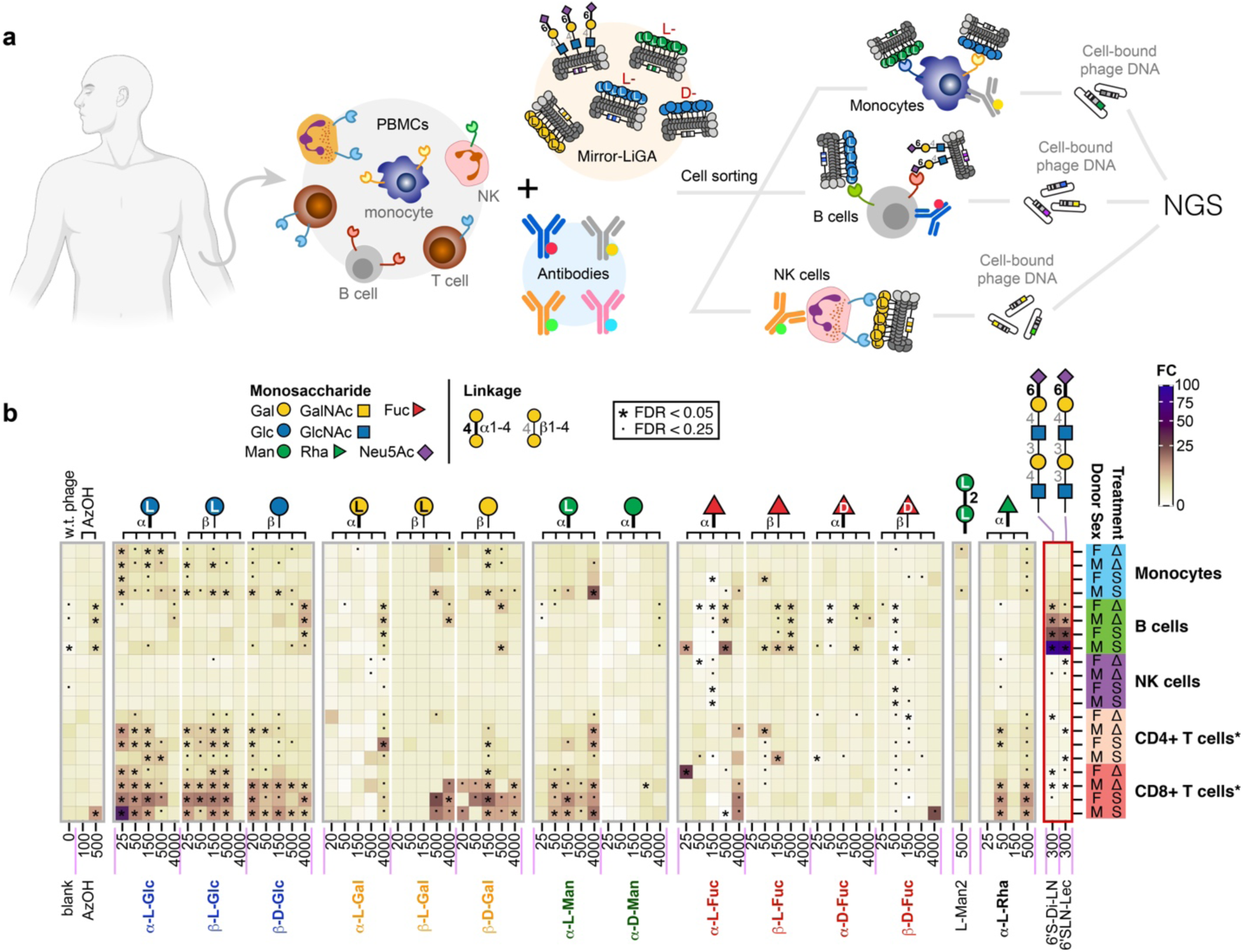
Binding of mirror-LiGA to human PBMCs. (a) Heat-map summary of enantiomeric glycan binding to cell subsets sorted from bulk PBMCs. Cells were treated by sialidase (S) or heat-inactivated form of the same enzyme (Δ) prior to binding with LiGA. n = 4 (two male and two female donors). Two sialylated glycans were used to benchmark glycan bindings. All cell subsets were sorted from PBMCs by FACS after LiGA binding, except for CD4⁺ and CD8⁺ T cells, for which pre-isolated cells were directly subjected to glycan-binding profiling without FACS sorting.

Pre-isolated CD8^+^ T cells exhibited the broadest mirror-glycan recognition profile. These cells bound L-Man but showed little corresponding recognition of D-Man, and recognized β-linked D-and L-Gal conjugates while showing weaker or inconsistent binding to linked α-L-Gal. They also bound Dand L-Glc conjugates. CD4^+^ T cells similarly recognized several glucose-containing glycans, although their overall binding profile was less extensive than that of CD8^+^ T cells. These results suggest the presence of previously uncharacterized glycan-binding activities on human T cells that can discriminate among closely related glycan stereoisomers and anomers. Together, these data show that human peripheral immune cells are not uniformly insensitive to mir-ror-image glycans. Instead, selected L-configured monosaccharides are recognized in a cell-type, configurationand displayed density-dependent manner, with the most extensive binding observed for pre-isolated CD8^+^ T cells.

### Natural human IgM exhibits broad recognition of mirror-image glycans

To test whether Land D-glycan monomers are recognized by human antibodies, we measured the interaction between mirror-LiGA and the IgM repertoire from 13 human donors (**Fig. 5**). The assay was conducted as described previously^40^, using IgM directly enriched from human serum sample in well plates coated by anti-human-IgM antibodies (**Fig. 5b**). We not only observed the recognition of L-glycans by IgM, but this recognition varies between individuals. For example, 6 of 13 donors contained IgM that bound to L-Man and L-Man-α-(1→2)-L-Man. The latter L-man-nobiose exhibited no measurable binding to any purified proteins or cells measured to date. The IgM from two other donors (# 688 and # 1109) recognized L-Man but exhibited no binding to L-Man-α-(1→2)-L-Man. In contrast, every tested donor contained antibodies against the classical α-D-Gal-α-(1→3)-D-Gal (α-Gal) xenoantigen and 12 out of 13 contained IgM that bound α-L-rhamnose. The monomers that showed similar across-the donor recognition were β-D-Fuc and α-D-Fuc, both are common components of OPS of diverse microorganisms.

**Figure 5.**
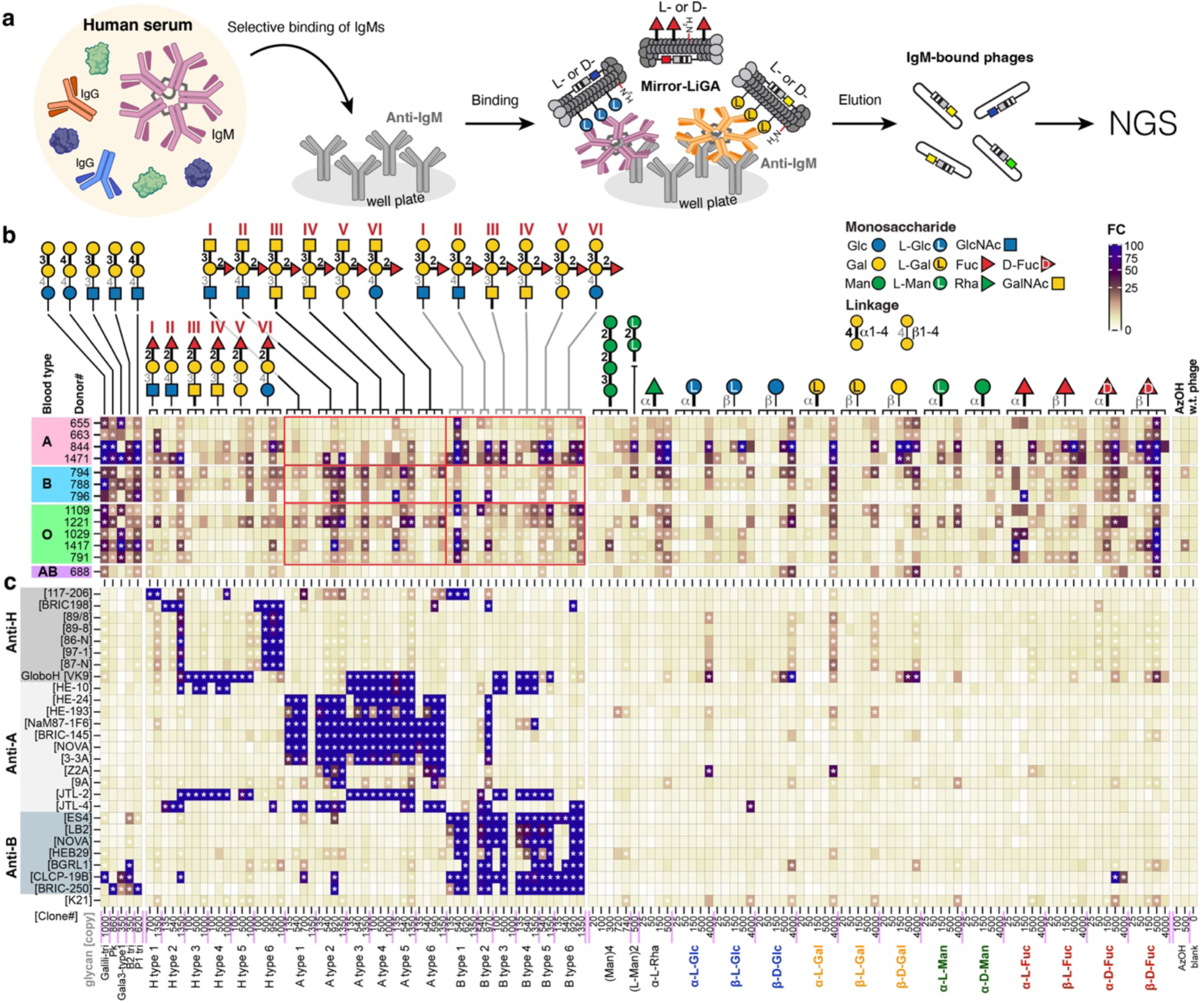
Human IgM repertoire recognizes _L_-glycans. **(a)** Scheme of profiling glycan binding of human IgM repertoire using LiGA. **(b)** Human IgM repertoire recognizes _L_-glycans including _L_-Man which has been showing little to no binding with the tested lectins. **(c)** Purified anti-human blood group antigen antibodies exhibit only weak recognition to _L_-Glc, _L_-Man and _L_-Gal.

Difficulty in isolation of monoclonal anti-glycan IgM clones make investigation of specific IgM components non-trivial. As alternative to *de novo* isolation of clonal IgM, we investigated recognition of a panel of existing anti-glycan antibodies with focus on 30 clonal IgM and IgG that recognize blood group and Lewis glycans (**Fig. 5c** and **Extended Data Fig. 4**). Expanded blood group glycan binding profiles of the tested antibodies correlated with reported specificities (**Extended Data Fig 4**). We further observed that half of the antibodies exhibit cross reactivity with L-Glc and L-Gal at high display densities, for specific antibody clones (**Fig. 5c**). The trait was clone-specific not class-specific: for example, only 2 out of 7 anti-B-group antibodies bound to D-Fuc; the same clones did not bind L-Fuc. The strong binding of clone CLCP-198 to α-D-Fuc was unique, with no detectable binding to β-D-Fuc and α-L-Fuc, nor any other glycan monomer. While binding of purified anti-glycan antibodies to L-Man was detectable and significant (FDR < 0.05), the magnitude of binding was too weak to explain the strong binding of the IgM repertoire to L-Man or L-mannobiose. The recognition of L-glycans is likely a combination of cross-reactivity of dedicated IgM with known recognition profiles and the presence of L-glycan IgM clones of hitherto unknown origins.

### *In vivo* biodistribution of glycosylated phages modified by L-glycans

Encouraged by the discovery of how prevalent the recognition of mirror-image glycans by GBPs from plants, human serums, and human peripheral immune cells, we sought to investigate if L-glycans can direct the preferential distribution / enrichment of phages to different organs and cell types in live animals. To this end, we injected mirror-LiGA into mice (strain C57BL/6J) and allowed it to circulate for 20 to 40 minutes before surgically removing livers and spleens. Splenic T and B cell subpopulations were then isolated by FACS to investigate organ and cell-type specific enrichment of L-glycans (**Fig 6**).

**Figure 6.**
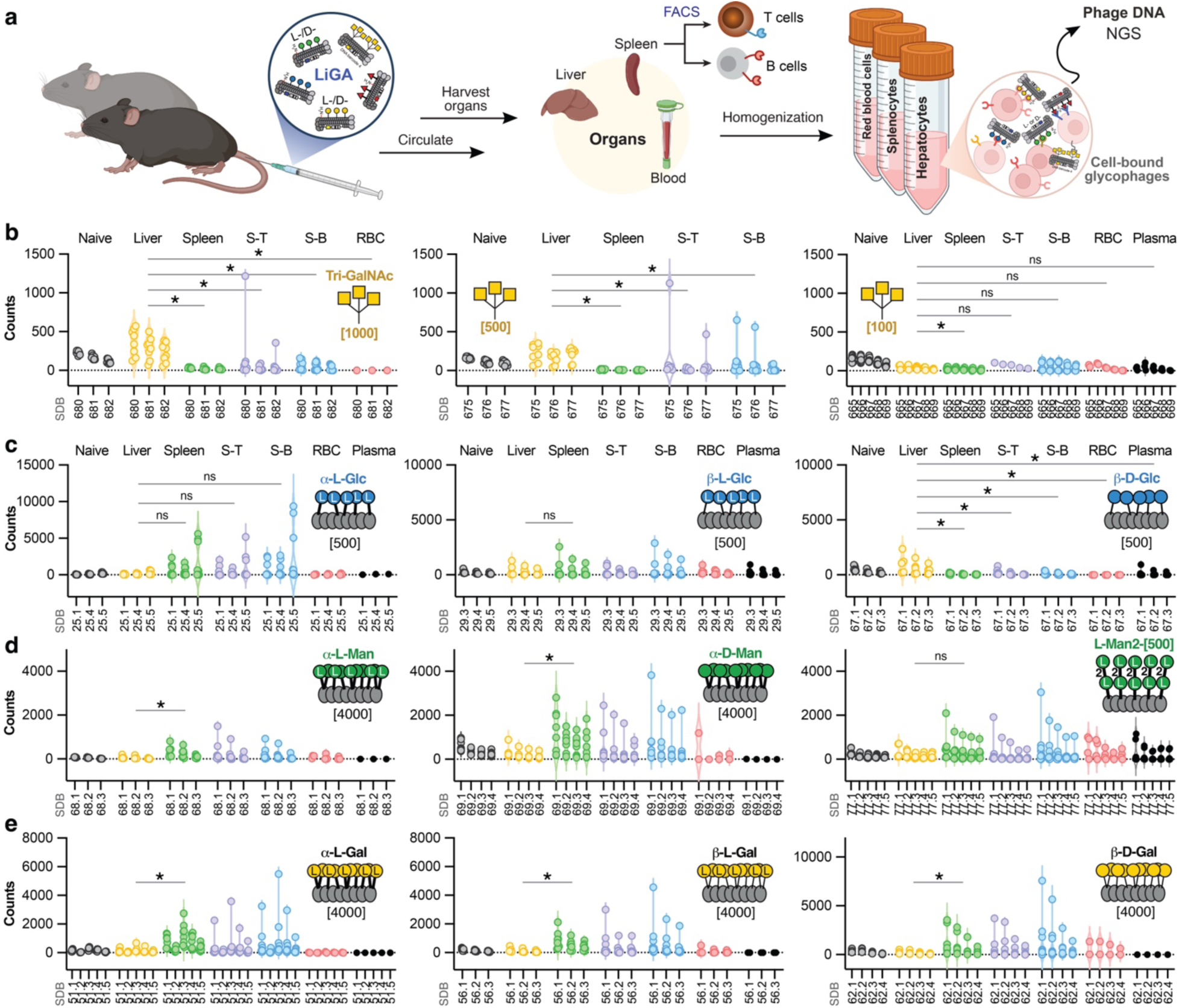
In vivo biodistribution of enantiomeric glycans in live mice. **(a)** Scheme of profiling the organ distribution of enantiomeric glycans in mice using LiGA. Splenocytes were further sorted by FACS into splenic T (S-T) and B lymphocytes (S-B) to examine binding of enantiomeric glycans at the cellular level. **(b)**-**(e)** Distribution of glycans among tested mouse organs. Independent experiments (n = 2 to 4 for different organs) were performed on different dates using 2-4 mice per experiment. Naïve, injected LiGA; S-T, FACS-sorted splenic T lymphocytes; S-B, FACS-sorted splenic B cells; RBC, red blood cells; SDB, “silent” double-barcoded phage clones. ns, statistical non-significant; * p < 0.05.

We spiked mirror LiGA with phage modified with tri-GalNAc, a well-established hepatocyte-targeting ligand used in multiple FDA-approved oligonucleotide therapeutics, to benchmark the biodistribution of glycan-decorated phage in mice^41–43^. Indeed, we observed specific enrichment of tri-GalNAc in the liver in independent experiments. In the LiGA mixture, three monoclonal phages were used to barcode tri-GalNAc, and they all showed specific accumulation in the liver. The hepato-tropism of tri-GalNAc-conjugated phages decreases at reduced tri-GalNAc display densities on the phage (**Fig. 6b**). In comparison, bacteriophages with unmodified coats showed no detectable organ tropism, with no preferential enrichment in any analyzed organ (**Extended Data Fig. 5i**). Interestingly, D-Glc showed enrichment in the liver but not in the spleen, while the L-Glc showed no significant enrichment in either spleen or liver (**Fig. 6c** and **Extended Data Fig. 5a,b)**.

L-Glc generally showed preferential binding towards L-Fuc binding lectins. Thus, we expected L-Gal would have similar organ enrichment as L-Glc. Indeed, both αand β-L-Gal showed similar enrichment to spleen but not to liver as L-Glc (**Fig. 6e**). Surprisingly, β-D-Gal also showed preferential binding towards spleen and splenic T and B cells but not to liver in contrast to β-D-Glc (**Fig. 6c,e**). In comparison to the distinct biodistribution of L-/D-Glc, both Land D-Man displayed at 4,000 glycans per phage showed similarly significant enrichment towards spleen, splenic B and T cell populations, but not to liver tissue. However, the D-Man modified phages showed much stronger enrichment in spleen than L-Man. Here, we observed that the L-disaccharide (L-Man-α-(1→2)-L-Man) showed similar biodistribution as L-Man when displayed on phages at 500 glycans per viron (**Fig. 6d**). The observed biodistribution and avoidance of hepato-tropism suggests that scaffolds like L-Man and L-Glc could be used to build targeting moieties for extrahepatic delivery.

## Discussion

Glycan-protein interactions exhibit a balance between specificity and plasticity. The mimicry of glycan reports reinforce more than fifty years of observations of promiscuity in carbohydrate processing enzymes; for example D-Ara*p* can be processed by enzymes that recognize L-fucose including fucosidases^44^, fucose dehydrogenases^45,46^, and fucosyltransferases^47^. While GBPs canonically recognize only one type of glycan, we show that it is possible for many proteins to recognize L-glycans. Inhibition of the binding of DC-SIGN to its native mannose and fucose-rich substrates by L-Glc further emphasises such plasticity. The high cost of L-glycan monomers precluded us from extensive inhibition studies that use soluble L-glycans. The mimicry is symmetric because soluble D-mannose inhibits interactions between multivalent displays of L-glycans and cell-surface displays of DC-SIGN. Future exploration can expand on these inhibition studies and even employ elution of proteins by diverse soluble L-glycans for omics-level discovery of protein-carbohydrate interactions that can be mimicked by L-glycans. Such recognition presents an unexplored avenue.

The ability of L-Glc to inhibit DC-SIGN binding has implications for how mirror organisms might interact with a homochiral host. Mirror organisms are generally predicted to evade many conventional immune-recognition pathways because their proteins, glycans and other biomolecules would have the opposite stereochemistry from those encountered by natural immune receptors. However, our results suggest that such stereochemical orthogonality may not be absolute: the natural immune lectin DC-SIGN retained measurable recognition of L-configured glucose-containing ligands, and this interaction was competitively inhibited by soluble L-Glc. Thus, at least some host lectins may retain partial cross-chiral recognition of mirror-cell-surface glycans, potentially enabling weak adhesion, uptake or immune sensing of mirror organisms.

Conversely, by mirror symmetry, a mirror DC-SIGN-like lectin expressed by a mirror organism would be expected to recognize L-mannose- or L-fucose-containing glycans but would also potentially interact with D-Glc. Because circulating D-Glc is normally present at approximately 4–8 mM, its concentration overlaps with the L-Glc IC_50_ values measured here for natural DC-SIGN. Human blood glucose could therefore partially occupy or competitively inhibit analogous carbo-hydrate-binding sites on mirror microbial adhesins, reducing their ability to bind host glycans. We note this possibility cannot be inferred quantitatively from the present experiment, because soluble-monosaccharide competition might not reproduce the avidity, receptor density or glycan organization present at a host-microorganism interface. Nevertheless, this mechanism could constitute an unanticipated chemical barrier to mirror-organism adhesion in the bloodstream.

This report focused on proteins of mammals, plants, immune cells, and whole mammalian organisms. Biodistribution of M13 bacteriophage particles modified by L-glycans in other organisms is in principle feasible. We limit our investigation to monosaccharides, as the recognition of terminal monosaccharide epitopes by GBPs is a common mechanism for interactions. Synthesis of core di- and tri-saccharides present an avenue for further exploration. Preliminary exploration of disaccharide L-Man-α-(1→2)-L-Man only perpetuated an observed resilience of L-Man to recognition by all tested GBPs. We were unable to find any purified lectins that bound to disaccharide L-Man-α-(1→2)-L-Man to any appreciable extent. Lack of binding is an offers a challenge that might be resolved by a metagenomic, environmental or bioengineering search for microorganisms and proteins that might recognize this simple disaccharide. Resistance to binding is an attractive trait and might be attractive in prevention of biofouling. Lack of any measurable binding to any lectins was contrasted by the recognition of the same disaccharide L-Man-α-(1→2)-L-Man by IgM from 6 out of 12 human donors. The origin of these IgM antibodies is presently unclear, but it indicates the capacity of immune system to recognize glyco-epitopes that are mirror image of the terminal mannose epitopes in glycocalyx of many organisms.

Mirror-LiGA offered observations that provide an evidence-based, fundamental contribution to the discussion of the risks of mirror-life and the ability of conventional, extant life forms to interact with such life. Interaction between extant glycan binding proteins and a multivalent display of mirror image glycans that will be present in the glycocalyx of the mirror-life organisms is not silent. Recognition of the multivalent display of mirror-glycans by proteins, antibodies and immune cells *in vivo* can be studied decades before mirror-life is created. An alternative approach to studying this problem is synthesis of lectins made of D-amino acids and these investigations are presently ongoing. Not only such D-lectins can be tested in traditional glycan arrays, but also they can be explored in metagenomic studies such as Lectin-Seq^48^ and DNA-encoded formats such as LiLA^49^ and GOAT-seq^50^ to identify whether naturally expressed glycans on the surface of extant microorganisms are recognized by enantiomeric lectins. By symmetry, this investigation will teach how natural lectins made of L-amino acids interact with complex enantiomeric glycans that would be very difficult to synthesize.

Beyond serving as building blocks in nucleic acids and energy source, the role of glycans in the origin of life is rarely discussed; however, presence of glycans building blocks early in the origin of life suggests important of glycans and GBP-glycan recognition for evolution of living organisms. A hypothetical Last Universal Common Ancestor (LUCA) contains genes that use MurNAc to build a cell wall^51–53^ as well as putative glycan-binding proteins. Meteorites^54^ and the asteroid Bennu^55^ contain monosaccharides in amounts comparable to amino acids. Breslow and co-workers suggested that the prehistoric accumulation of L-amino acids might have catalyzed the preferential accumulation of D-glyceraldehyde, a possible precursor to D-monosaccharides^56^. It is not clear what role glycan chirality played before emergence of complex biogenic synthesis of glycans became possible. High-throughput tools such as mirror-LiGA that accumulate information about cross-chiral recognition between glycans and proteins can offer important information in evolutionary significant of glycans and their co-evolution with protein that recognize unique stereoisomers of glycans.

## Extended Data Figures

**Extended Data Figures 1:**
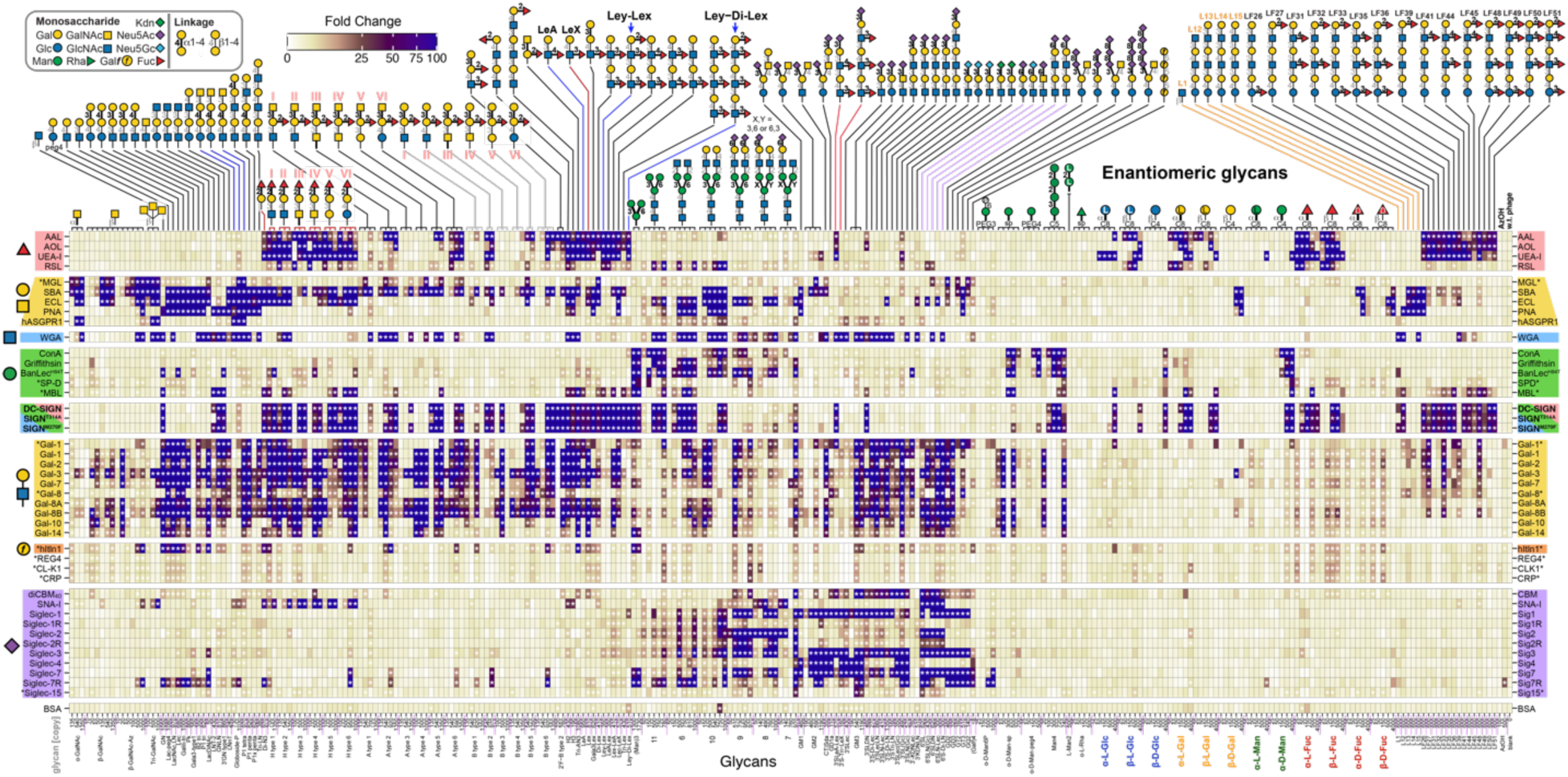
Recognition of enantiomeric glycans by diverse glycan binding proteins. The tested purified GBP include 37 unique lectins, five point mutants of Siglecs and DC-SIGN and different expression variants for Galectin-1 and DC-SIGN. Besides canonical glycan binding was observed for all tested lectins, we noticed various of extant plant-derived or human-derived lectins can recognize mirror-image monosaccharides in a density-dependent manner.

**Extended Data Figures 2:**
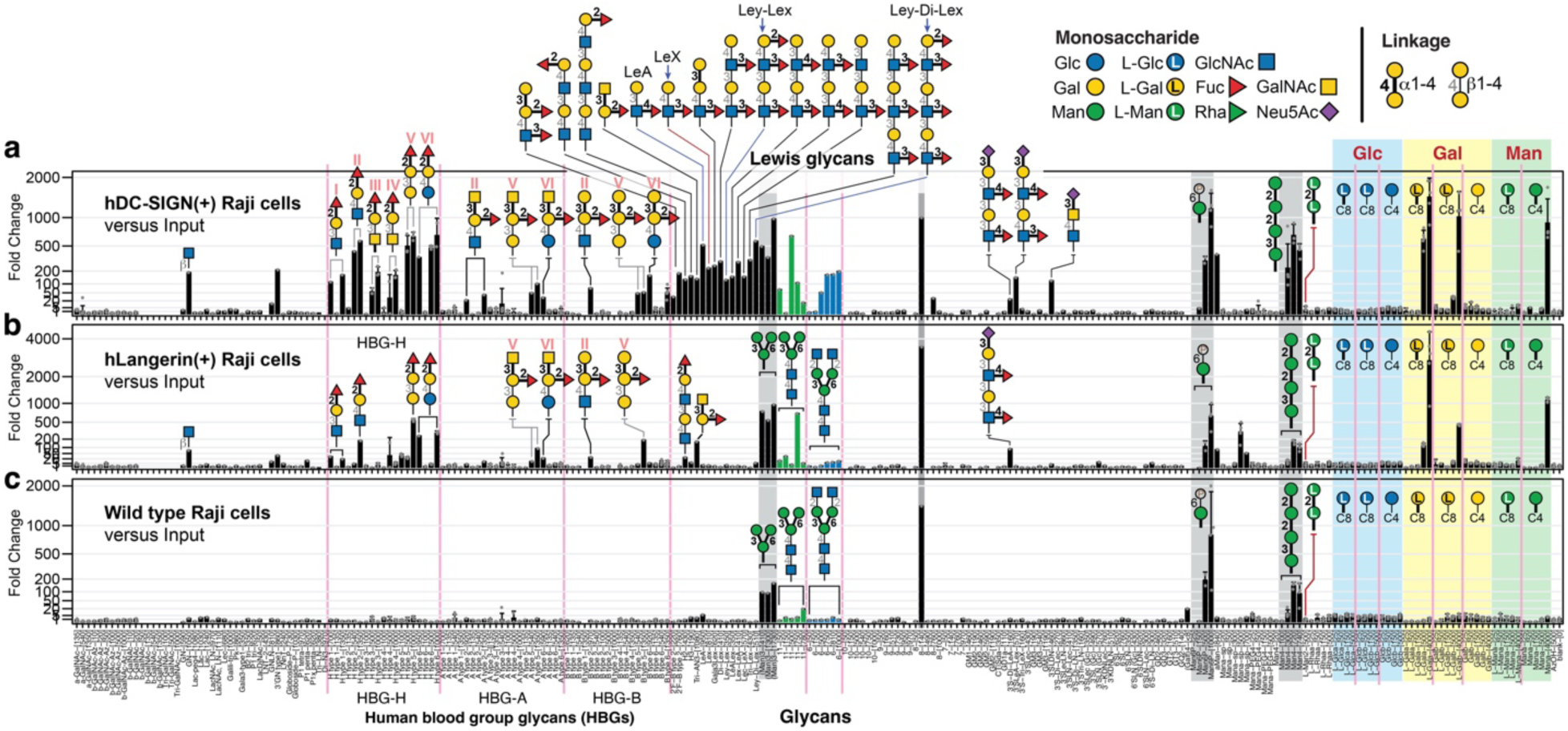
Glycan-binding profiles of human DC-SIGN^+^ and Langerin^+^ Raji cells. **(a)** Glycan-binding profile of DC-SIGN^+^ Raji cells. In addition to significant binding to H-type glycans, A type II, V and VI glycans, B type II, V and VI glycans, Lewis glycans and high-mannose glycans, DC-SIGN^+^ Raji cells showed strong binding to α-/β-_L_-Gal and α-_D_-Man, but not to β-_D_-Gal or α-_L_-Man. n = 5 for DC-SIGN^+^ Raji cells. **(b)** Glycan-binding profile of Langerin^+^ Raji cells. Langerin^+^ cells showed strong binding to α-/β-_L_-Gal and α-_D_-Man, but not to β-_D_-Gal or α-_L_-Man. n = 5 for Langerin^+^ Raji cells. **(c)** Glycan-binding profile of the wild type Raji cells. n = 5 for the tested Raji cells.

**Extended Data Figures 3:**
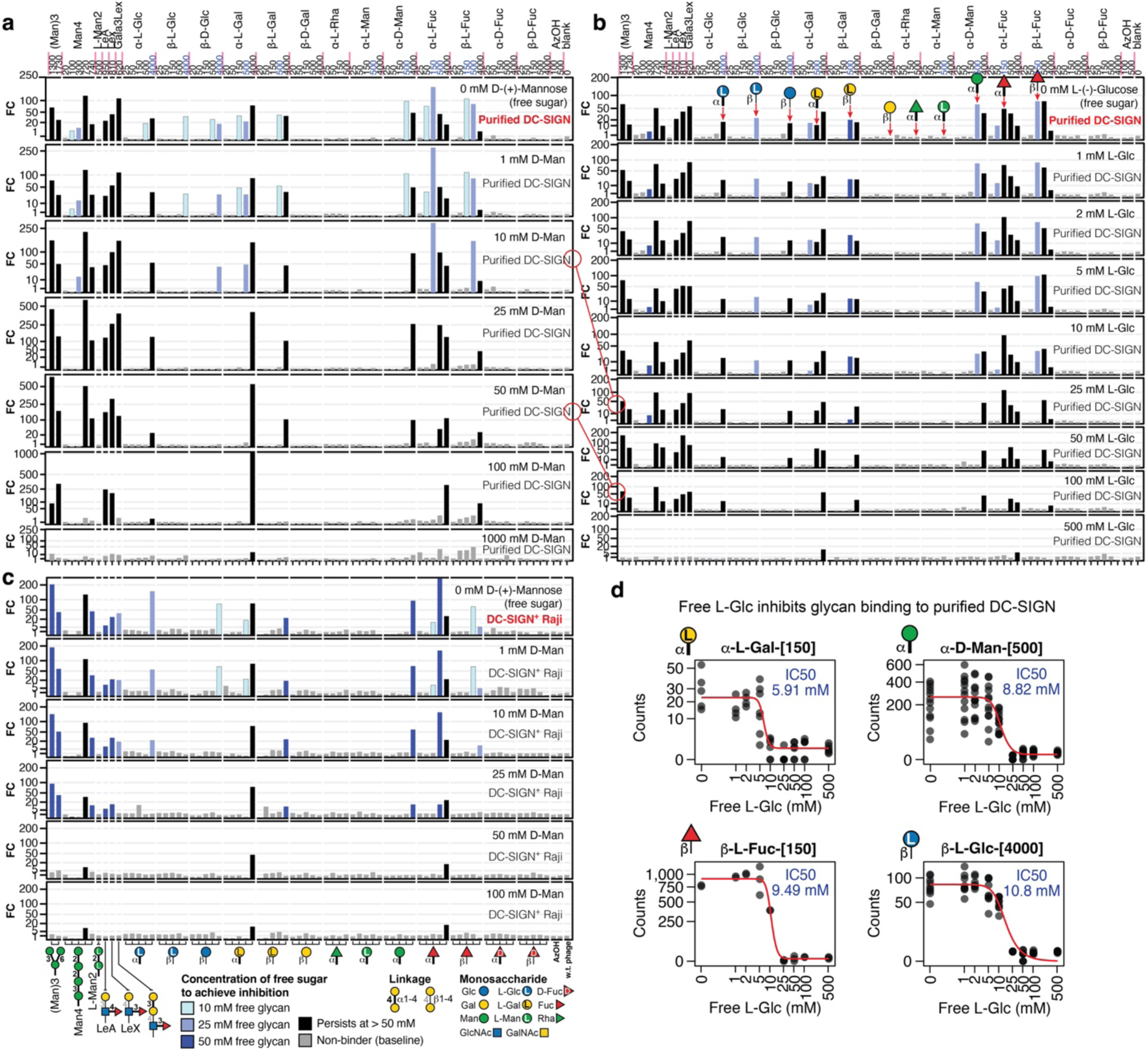
Inhibition of glycan binding to DC-SIGN by free _D_-mannose and _L_-glucose. **(a)** Inhibition of glycan binding to purified DC-SIGN extracellular domain (ECD) by free _D_-mannose. n = 3. **(b)** Inhibition of glycan binding to purified DC-SIGN ECD by free _L_-glucose. n = 3. **(c)** Inhibition of glycan binding to DC-SIGN+ Raji cells by free _D_-Man. **(d)** Inhibition curves and IC_50_ extrapolated from the normalized sequencing counts of purified DC-SIGN ECD by free _L_-Glc.

**Extended Data Figures 4:**
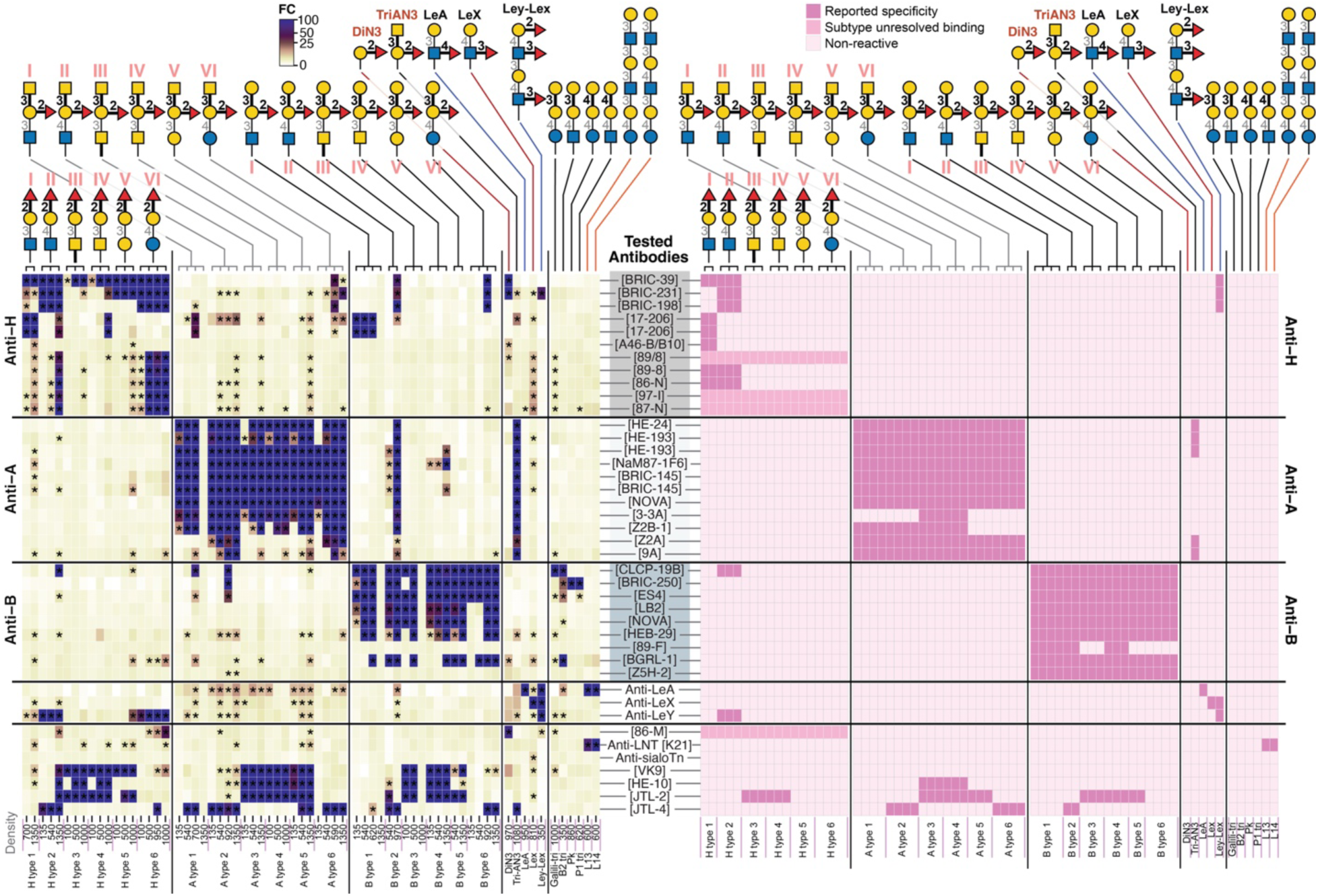
Comparison of glycan-binding specificities measured by LiGA (left) with reported specificities (right). Semi-qualitative assessment data adapted from previous publications or provided from were converted to a heat-map representation. Reported specificity measured by ELISA^57^: A46-B/B10 (anti-HBGH), Z2B-1 (anti-HBGA), 89-F (anti-HBGB), JTL-2, and JTL-4. Reported specificity measured by neoglycoprotein glycan microarray^58,59^: HE-10, HE-24 (anti-HBGA), HE-193 (anti-HBGA), Z2A (anti-HBGA), 9A (anti-HBGA), CLCP-19B (anti-HBGB), 7LE (anti-Lewis A), and F3 (anti-Lewis Y). Reported specificity measured by multiplex bead immunoassay (Luminex)^60^: Nova anti-A, Nova anti-B, JTL-2, and JTL-4. Specificity of all other antibodies were cited from suppliers. We did notice that reported specificity varies between assays. For example, HE-10 was previously reported to bind H type 3/4/5, A type 3/4/5, and B type 3/4/5^61^, whereas Gildersleeve and coworkers observed binding of HE-10 to A type 3/4, with no detectable binding to A type 5, B type 5, and H type 3/4/5^58^. Similarly, JTL-2 was reported to bind A/B type 3/4 and JTL-4 to bind A/B type 2 by ELISA^57^, whereas multiplex bead immunoassays revealed additional binding of JTL-2 to A/B type 5 and of JTL-4 to A type 6^60^.

**Extended Data Figures 5:**
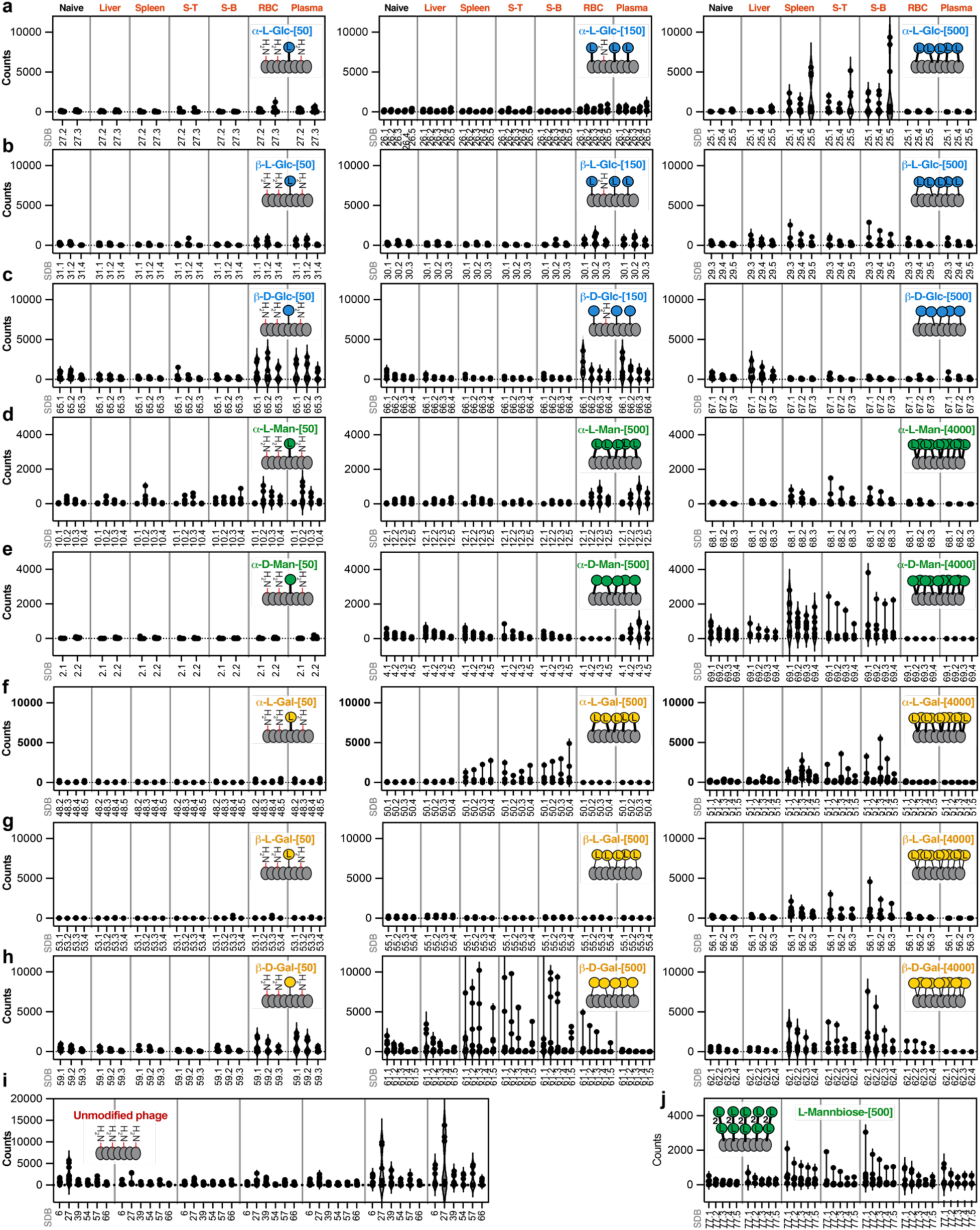
In vivo biodistribution of phages modified with enantiomeric glycans at low-medium-high densities. Density, or copies of glycans per viron, was represented as “[x]”. Naïve, injected LiGA; S-T, FACS-sorted splenic T cells; S-B, FACS-sorted splenic B cells; RBC, red blood cells; SDB, “silent” double-barcoded phage clones.

## Methods

### Glycan synthesis

All mirror-image monosaccharides were synthesized through GlycoNet Integrated Services (GIS) following previously reported procedures^12^. Detailed synthetic protocols are available upon request. Expanded blood group glycans (**Fig 2, 5** and **Extended Data Fig. 1,2,4**) were synthesized by following previous publications^62–64^. Synthesis of human milk oligosaccharides (**Extended Data Fig. 1**) were synthesized according to published procedures^65,66^. The source and synthesis of all other glycans in LiGA (**Extended Data Fig. 1**) have been described previously^67–69^. Detailed synthetic procedures and NMR spectra for α-1,2-L-mannobiose are provided in the **Supplementary Information**.

### Construction of mirror-LiGA

Construction and isolation of “silent” double-barcoded phages (SDB phage), conjugation of glycans to monoclonal SDB phages, and preparation of LiGA can be found in previous publications^67,70^. Phage glycosylated with mirror-image glycans were prepared by reacting clonal SDB phages (∼10^13^ PFU ml^-1^) with glycan-NHS ester in borate buffer (100 mM, pH 8.0) at room temperature for 1 h, followed by purification of glycophages using a Zeba column (Thermo Fisher Scientific, 89882). This conjugation process was repeated to allow glycan ultra-high density displaying. Resulting glycophages were analyzed using MALDI-TOF MS and stored in 1:1 (v/v) PBS:glycerol at –20 °C. MALDI-TOF MS spectra of mirror-image glycan modified SDB phages can be found in **Figure S1**. Composition of the tested mirror-LiGA is listed **Table S1**.

### Binding of mirror-LiGA to purified GBPs

A protein solution (50 µl, 50 μg ml^-1^) prepared in HEPES-buffered saline solution (HBS; 20 mM HEPES, 150 mM NaCl, 2 mM CaCl_2_, pH 7.4) was added to a high-binding 96-well plate (Corning, CLS3369). The plate was sealed with tape (Thermo Fisher Scientific, 15036) and incubated overnight at 4 °C. The next day, wells were washed with HBS containing 0.1% (v/v) Tween-20 (HBST; 3 × 200 µl), blocked with HBS containing 1% (w/v) BSA (HBSB; 200 µl), and then washed with HBST (3 × 200 µl), after which a LiGA solution (50 μl, 10^8^ PFU in HBS) was added and incubated at room temperature for 1 h. Wells were then washed with HBST (2 × 200 µl) and HBS solution alone (1 × 200 µl). Bound phages were eluted with acid (50 μl, 10 mM HCl, 9 min, room temperature) and neutralized with 5× Phusion HF buffer (25 μl; Thermo Fisher Scientific, F518L). Eluted phage samples were quantified by qPCR and submitted for deep sequencing.

### Binding of glycophages modified with mirror sugar to lectin microarray

Phages were labelled with fluorophore (0.2 mM TAMRA-NHS ester, Lumiprobe, 17120) in borate buffer (100 mM, pH 8.0) at room temperature and then purified using Zeba columns. Fluorescent phages were then glycosylated with glycan-NHS esters in borate buffer, purified using Zeba columns, and stored in 1:1 (v/v) PBS:glycerol at –20 °C. MALDI-TOF MS spectra of TAMRA-labeled glycophages can be found in **Figure S2**.

Glass slide printed with lectin microarray (manufactured in the Mahal laboratory at the University of Alberta) was thawed at room temperature and hydrated in ethanolamine/boric acid buffer (50 mM ethanolamine, 100 mM boric acid) for 1 h. The slide was sequentially washed with PBS containing 0.01% Tween-20 (PBST) and PBS, and dried on a slide spinner. Fluorescent glycophages (50 µl in PBS:glycerol solution) were mixed with binding buffer (50 µl; PBST supplemented with 0.1 mM CaCl_2_) and incubated with the lectin microarray at 30 °C for 1 h. Arrays were washed with PBST (2 × 150 µl) and PBS (2 × 150 µl), briefly dipped in deionized water, dried on a slide spinner, and scanned at 532 nm using a GenePix 4400A Microarray Scanner. Fluorescence values were extracted using GenePix Pro 7 software, and active binding was defined as background-subtracted fluorescence above 1,000 AU. Composition of the lectin microarray can be found in **Table S4–5**.

### Inhibition of mirror-LiGA binding to purified DC-SIGN by free sugars

A protein solution (50 µl, 50 μg ml^-1^ in HBS) was added to a high-binding 96-well plate, sealed with tape, and incubated overnight at 4 °C. The next day, wells were washed (HBST; 3 × 200 µl), blocked (HBSB; 200 µl), and washed (HBST; 3 × 200 µl), after which a LiGA solution (50 μl, 10^9^ PFU in HBS containing 0–500 mM free glycans) was added and incubated at room temperature for 1 h. Wells were washed with HBST containing corresponding concentrations of soluble glycan (2 × 200 μl) and HBS solution alone (1 × 200 μl). Bound phages were eluted with acid (50 μl, 10 mM HCl, 9 min, room temperature) and neutralized with 5× Phusion HF buffer (25 μl). Eluted phage samples were quantified by qPCR and submitted for deep sequencing.

### Inhibition of mirror-LiGA binding to DC-SIGN^+^ cells by free glycans

Stable transfected DC-SIGN^+^ Raji cells cultured in complete medium (RPMI-1640 basal medium supplemented with 10% Fetal Bovine Serum (FBS), 1% Penicillin/Streptomycin, and 1× GlutaMAX) were harvested at confluence and aliquoted (1 × 10^6^ cells) to FACS tubes (Corning, 352054). Cells were pelleted (300×g, 5 min), resuspended in LiGA solution (100 μl, 10^9^ PFU in HBSB containing 0–500 mM free glycans), and incubated on ice for 1 h. Cells were then washed with HBS containing 0.1% BSA and corresponding concentrations of soluble glycan (2 × 3 ml), then washed with HBS solution alone (1 × 800 μl). Cells were then pelleted (300×g, 5 min, 4 °C), resuspended in lysis buffer (1× Phusion HF buffer containing 0.1 mg ml^-1^ RNase A, 50 μl) and vortexed at maximum speed. The cell lysate was centrifuged (2,000×g, 3 min), and the supernatant was treated with proteinase K (0.1 mg ml^-1^, 55 °C, 10 min) and heat-inactivated (95 °C, 10 min). After pelleting cell debris (21,000×g, 3 min), the supernatant containing bound phage DNA was collected and submitted for deep sequencing.

### Binding of mirror-LiGA to human PBMCs and isolated T cells

Cryopreserved peripheral blood mononuclear cells (PBMCs) and isolated human peripheral blood CD4+ and CD8+ T cells from healthy adult donors (STEMCELL Technologies, 70025.2, 70026, 70027) were thawed in pre-warmed complete medium consisting of RPMI supplemented with 10% FBS, 100 U ml^-1^ penicillin, 100 μg ml^-1^ streptomycin, non-essential amino acids (Gibco, 11140-050) and 55 μM 2-mercaptoethanol, and allowed to recover overnight at 37 °C under 5% CO_2_.

The next day, cells were treated with active or heat-inactivated sialidase (from *Vibrio cholerae,* 550 mU) for 45 min at 37 °C. The heat-inactivated controls were prepared by incubating the enzyme at 60–65 °C for 30 min. Cells were then washed with FACS buffer (Ca^2+^and Mg^2+^-free HBSS supplemented with 1% BSA and 2 mM EDTA) and pelleted (300×g, 10 min).

For LiGA binding to pre-isolated CD4+ and CD8+ T cells, cells were resuspended in LiGA solution (100 µl, 10^9^ PFU in HBSB) and incubated on ice for 1 h. Cells were then washed with HBS supplemented with 0.1% BSA (2 × 3 ml) and HBS solution alone (1 × 800 µl). Cell-bound phage DNA was collected by following treatment as described in **Inhibition of mirror-LiGA binding to cells by free glycans**.

For LiGA binding and flow-cytometric sorting of bulk PBMCs, cells were first stained with Zombie Violet viability dye diluted 1:1,500 in PBS (BioLegend, 423113) for 10 mins at room temperature. The viability dye was quenched by washing cells with FACS buffer. Cells were then incubated with Fc-receptor blocking solution diluted 1:250 in FACS buffer (BioLegend, 422302) for 10 mins on ice, followed by incubation with phenotyping antibodies and LiGA for 1 h on ice. Cells were washed with FACS buffer and then sorted into monocyte, B-cell, NK-cell, CD4+ T-cell, and CD8+ T-cell populations using a five-way sort on a BD FACSymphony S6 (Centre for Immune Analytics, University of Toronto). Phenotyping antibodies used were as follows: anti-CD14 (BioLegend, 367123), anti-CD19 (BioLegend, 302237), anti-CD56 (BioLegend, 318355), anti-CD3 (BioLegend, 570470), anti-CD4 (BioLegend, 344685), and anti-CD8 (BioLegend, 344739). After sorting, cell-bound phage DNA was collected by following RNase A-proteinase K treatment as described in **Inhibition of mirror-LiGA binding to cells by free glycans**. All collected DNA samples were submitted for deep sequencing.

### Binding of mirror-LiGA to human serum IgM repertoire

Goat anti-human IgM (50 μl, 20 μg ml^-1^ in PBS; Jackson ImmunoResearch, 109-005-043) was added to a high-binding plate, sealed, and incubated at room temperature for 2 h. Wells were then washed with PBST (2 × 200 μl), blocked with 1% BSA in PBS for 1 h, and washed with PBST (3 × 200 μl). Human serum samples (50 µl, 1:100 diluted in PBS; Lori J. West lab, University of Alberta) were added to the well plate, sealed, and incubated overnight at 4 °C. On the next day, wells were washed with PBST (2 × 200 μl), then mirror-LiGA (50 μl, 10^9^ PFU ml^-1^ in PBS) was added and incubated at room temperature for 1.5 h. Wells were washed with PBST (3 × 200 μl), then bound phages were eluted from wells with acid, neutralized with 5× Phusion HF buffer, and submitted for deep sequencing.

### Mapping the biodistribution of mirror-LiGA in live mice

All animal procedures were approved by the University of Alberta Animal Care committee in accordance with the Canadian Council on Animal Care guidelines (AUP00002467). Mice were maintained under pathogen-free conditions at the University of Alberta animal facility.

Mirror-LiGA (0.2 ml, 1 × 10^11^ PFU ml^-1^ in PBS) was injected into the tail vein of 10–12-week-old male C57BL/6J mice. After 20 min of circulation, blood (0.1 ml) was collected from the tail vein into heparin-coated tubes. Upon anesthetization, mice were perfused with PBS, then liver and spleen were collected and kept in ice-cold PBS on ice. Blood samples were centrifuged (300×g, 5 min, 4 °C) to separate plasma and red blood cells (RBCs).

Organs were minced into ∼2 mm pieces using a single-edge razor blade and transferred into a 40 μm cell strainer (Fisher Scientific, 22-363-547) fitted onto a 50 ml Falcon tube. Tissues were homogenized by gently grinding with the plunger of a 3 ml syringe while rinsing with ice-cold PBS (5 ml). The resulting cell suspension was centrifuged (300×g, 5 min, 4 °C), resuspended in RBC Lysis Buffer (5 ml, 1:10; Invitrogen, 00-430-54), and incubated at room temperature for 5 min. Cells were washed with ice-cold PBS (20 ml), pelleted (300×g, 5 min, 4 °C), resuspended in FACS buffer (5 ml for liver and 1ml for spleen), and counted.

An aliquot of splenocytes (1 × 10^7^ cells in FACS buffer) was stained with an antibody cocktail on ice for 1 h and sorted into B-cell and T-cell populations on a Sony MA900 (Flow Core, University of Alberta). The antibody cocktail consisted of anti-mouse CD3 (1:100; Invitrogen, 11-0032-80) and anti-mouse CD19 (1:100; Invitrogen, 17-0193-80).

Cell-bound phage DNA was collected from homogenized tissues, sorted splenic T and B cells, plasma, and RBCs by following RNase A-proteinase K treatment as described in **Inhibition of mirror-LiGA binding to cells by free glycans** and then submitted for deep sequencing.

### Data analysis and statistics

Sequencing data was analyzed as previously described^70^. To account for variations in the amount of material sequences, the read counts were normalized based on the abundances of a set of unmodified (blank) phages. Then each group of experimental samples was compared to the reference (naïve) cases using EdgeR^71^. This tool models the distributions of read counts across replicate sequencings using negative binomial distributions and estimates the corresponding log fold-change and statistical significance (p-value) of each case. These p-values were then Benjamini-Hochberg adjusted to obtain False Discovery Rates (FDRs), before fold-changes and FDRs were averaged across replicate barcodes by taking median values. Cases with FDR < 0.05 are marked with asterisks, and those with 0.05 < FDR < 0.25 by dots.

## Data availability

All data required to assess the conclusions of this study are provided in the paper, the Supplementary Materials, and/or the associated auxiliary files. The LiGA datasets listed in **Table S2** are also publicly accessible at https://48hd.cloud. Supplementary information is available for this paper. Correspondence and requests for materials should be addressed to R.D.

## Acknowledgements

We would like to acknowledge that this work used services provided by CFI-MSI GlycoNet Integrated Services. We thank the staff at the University of Alberta mass spectrometry facility (Chemistry Department) for help with MALDI analysis and Sophie Dang at the molecular biology service unit for assistance with Illumina sequencing. We thank Dr. Bruce Motyka and Dr. Caishun Li for kindly providing human serum samples and purified anti-HBG antibodies. Cell sorting was performed at the University of Toronto Centre for Immune Analytics Core Facility (RRID: SCR_027612) and at the University of Alberta Faculty of Medicine & Dentistry Flow Cytometry Facility (RRID: SCR_019195) which receives financial support from the Faculty of Medicine & Dentistry and Canada Foundation for Innovation (CFI) awards to contributing investigators. Figures 4 and 6 were created by adapting templates from BioRender.com (https://Bio-Render.com/c9fhkbu).

## Author contributions

R.D. conceived the project. C.P. and M.T.L. constructed glycophages displaying mirror-image glycans and HMO glycans. S.G. and C.P. measured glycan binding by purified lectins and antibodies. N.T. conducted lectin microarray measurements. S.H. conducted inhibition of LiGA binding to DC-SIGN by soluble glycan inhibitors. M.E. measured glycan binding of human se-rum-derived IgM repertoires. H.C., V.A., and C.P. conducted glycan-binding profiling of human PBMCs and pre-isolated T cells. C.Y.Y.L. and C.P. measured biodistribution of mirror-LiGA in live mice. S.Y.W. synthesized the α-1,2-L-mannobiose. C.O.C., A.S.G., J.L., S.B., A.K., M.S.M., and L.J.W. expressed or provided the purified GBPs, DC-SIGN^+^ and langerin^+^ Raji cells, and human serum samples. C.P., E.J.C., S.G., S.H., and R.D. contributed to data analysis, statistical analysis, visualization, and figure preparation. R.D., L.J.E., L.L.K., T.L.L., M.S.M., C.R., L.K.M., C.C.Y., L.J.W., S.K.W., and M.T.L. contributed to manuscript writing and editing.

