## Supplementary figures and tables for "Biological recognition of mirror-image glycans"

**Table S6:** Fitted $IC_{50}$ for inhibition of glycophages binding to DC-SIGN$^{+}$ Raji by free D-Man. 26

**List of Abbrevations**

| | |
|---|---|
| **AAL** | *Aleuria aurantia* lectin |
| **Ac** | Acetyl- |
| **ACN** | Acetonitrile |
| **AOL** | *Aspergillus oryzae* lectin |
| **ASGPR1** | Asialoglycoprotein receptor 1 |
| **BanLec** | Banana lectin |
| **BSA** | Bovine serum albumin |
| **CL-K1** | Collectin Kidney 1 |
| **ConA** | Concanavalin A |
| **CRP** | C-reactive protein |
| **DC-SIGN** | Dendritic Cell-Specific Intercellular adhesion molecule-3-Grabbing Non-integrin |
| **ECL** | *Erythrina cristagalli* lectin |
| **EDC HCl** | 1-Ethyl-3-(3-dimethylaminopropyl)carbodiimide hydrochloride |
| **FACS** | Fluorescence-Activated Cell Sorting |
| **GBP** | Glycan-binding protein |
| **HBG** | Histo-blood group glycan(s)l |
| **HBSS** | Hank's Balanced Salt Solution |
| **HEPES** | (4-(2-Hydroxyethyl)-1-piperazineethanesulfonic acid) |
| **ITLN1** | Intelectin-1 |
| **LiGA** | Liquid Glycan Array |
| **LUCA** | Last Universal Common Ancestor |
| **MALDI-TOF** | Matrix-Assisted Laser Desorption/Ionization Time-of-Flight |
| **MBL** | Mannose-binding lectin |
| **MGL** | Macrophage galactose-type lectin |
| **NK cells** | Natural killer cells |
| **PBS** | Phosphate buffer saline |
| **PFU** | Plaque-forming unit |
| **PNA** | Peanut agglutinin |
| **REG-4** | Regenerating islet-derived protein 4 |
| **RFU** | Relative Fluorescence Units |
| **RSL** | *Ralstonia Solanacearum* Lectin |
| **rt** | Room temperature |
| **SBA** | Soybean agglutinin |
| **Siglecs** | Sialic acid-binding immunoglobulin-type lectins |
| **SNA-I** | *Sambucus nigra* agglutinin I |
| **SP-D** | Surfactant protein D |
| **TAMRA** | 5-Carboxytetramethylrhodamine |
| **UEA-I** | *Ulex europaeus* agglutinin I |
| **WGA** | Wheat germ agglutinin |

### 1.Biological methods

#### 1.1. Cloning

Bacterial expression constructs were generated in pET-24a. Recombinant human galectin-1 (UniProt P09382) and galectin-8 (UniProt O00214) were synthesized and cloned by Twist Bioscience. Galectin-1 was cloned with a C-terminal $His_6$ tag and AviTag, and galectin-8 was cloned with an N-terminal AviTag and a C-terminal $His_6$ tag.

Mammalian expression constructs were generated in pcDNA3.1. Human MGL (CLEC10A construct lacking residues 1–119; UniProt J3KR22), MBL (MBL2; UniProt P11226), DN-SPD (surfactant protein D construct lacking residues 24–45; UniProt P35247), intelectin-1 (ITLN1; UniProt Q8WWA0) and REG4 (UniProt Q9BYZ8) were synthesized as described previously [bioRxiv 2026.02.23.707224; PMID 40421735] and cloned by Twist Bioscience with an N-terminal AviTag and $His_6$ tag. CL-K1 (UniProt Q9BWP8) and CRP (UniProt P02741) were cloned identically, but with a C-terminal AviTag and $His_6$ tag. All construct sequences were verified by Sanger sequencing (Quintara Biosciences).

#### 1.2. Protein expression and purification

Galectin-1 and galectin-8 were expressed in *E. coli* D3 cells from pET-24a vectors. In brief, liter-scale cultures were grown to mid-log phase ($OD_{600}$ = 0.3–0.5), induced with 0.4 mM IPTG and incubated for 16 h at 18 °C. Cells were collected by centrifugation (3,000g, 30 min), and pellets were stored at −20 °C until purification. Pellets were resuspended in Ni-NTA loading buffer (20 mM sodium phosphate, 500 mM NaCl, 20 mM imidazole, pH 7.4), lysed using B-PER II reagent (Thermo Fisher Scientific, 78260), and clarified by centrifugation (24,000g, 60 min).

MGL, MBL, DN-SPD, CL-K1, CRP, intelectin-1 and REG4 were expressed transiently in Expi293F cells using the ExpiFectamine™ 293 Transfection Kit (Thermo Fisher Scientific, A14524) according to the manufacturer's instructions. Conditioned medium was harvested 3–5 d after transfection by centrifugation and passed through a 0.2 µm PES filter.

Filtered $His_6$-tagged proteins were loaded onto a Ni-NTA column (Cytiva), washed with loading buffer and eluted using a gradient to elution buffer (20 mM sodium phosphate, 500 mM NaCl, 500 mM imidazole, pH 7.4). Fractions were analyzed on stain-free tris-glycine gels. Purified galectins were dialyzed overnight at 4 °C into PBS containing 7 mM β-mercaptoethanol, and mammalian-expressed lectins were dialyzed overnight at 4 °C into 20 mM HEPES, 150 mM NaCl and 1 mM EDTA, pH 7.4, using 10 kDa MWCO Slide-A-Lyzer dialysis cassettes (Thermo Fisher Scientific). Protein concentrations were determined by $A_{280}$ using extinction coefficients and molecular weights calculated with ProtParam. Proper folding was confirmed by differential scanning fluorimetry or circular dichroism before use.

#### 1.3. Protein biotinylation

AviTag-bearing lectins were biotinylated using an enzymatic BirA biotinylation kit (Avidity) according to the manufacturer's recommendations for 2 h at 30 °C. Biotinylated proteins were desalted using Zeba 1.5 ml 7K MWCO spin columns (Thermo Fisher Scientific, 89882), and biotinylation was confirmed by streptavidin gel-shift. Biotinylated human Siglec-15 (SG5-H82E9) was obtained from ACROBiosystems.

**1.4. Analyzing glycosylated phages using MALDI-TOF mass spectrometry**

Chemical modifications on phages were analyzed by MALDI-TOF MS using an AB Sciex Voyager Elite MALDI mass spectrometer equipped with a MALDI-TOF pulsed nitrogen laser (337 nm; 3-ns pulse up to 300 µJ per pulse) operating in full-scan MS in positive ionization mode. For sample preparation, a matrix solution (0.7 µl, 10 mg $ml^{-1}$ sinapinic acid in 4:1, v/v, acetone:methanol) was spotted on a 100-spot MALDI target plate (stainless steel) and allowed to dry. Phage samples were prepared by mixing a phage solution (2 µl in water or PBS, avoiding high salt or glycerol) with a layer 2 matrix (4 µl, 10 mg $ml^{-1}$ sinapinic acid in 1:1, v/v, 0.1% TFA in water:acetonitrile). The phage-matrix mixture (2 µl) was then deposited onto the dried first matrix layer and allowed to dry. The dried phage sample spots were rinsed with 0.1% TFA in water (3 × 10 µl) prior to analysis. MALDI-TOF MS spectra of glycophages modified with mirror-image monosaccharides at various densities can be found in **Figure S1** and fluorescent labeled glycophage used in lectin microarray assay can be found in **Figure S2**.

### 2. Chemical methods used in glycan synthesis

#### 2.1. General NMR procedure

$^1$H and $^{13}$C NMR spectra were recorded on a Bruker AV-400 spectrometer. Chemical shifts are reported in parts per million (ppm, δ scale). For $^1$H NMR spectra, chemical shifts were referenced to the residual solvent signals of $CDCl_3$ (δ 7.24 ppm), $CD_3OD$ (δ 3.31 ppm), $D_2O$ (δ 4.80 ppm), or $(CD_3)_2SO$ (δ 2.50 ppm). For $^{13}$C NMR spectra, chemical shifts were referenced to the solvent signals of $CDCl_3$ (δ 77.0 ppm) or $CD_3OD$ (δ 49.15 ppm). NMR data are reported as follows: chemical shift, multiplicity (s = singlet, d = doublet, t = triplet, q = quartet, dd = doublet of doublets, m = multiplet, br = broad), coupling constant (*J*, Hz), and integration.

#### 2.2. Synthesis of α-1,2-L-mannobiose

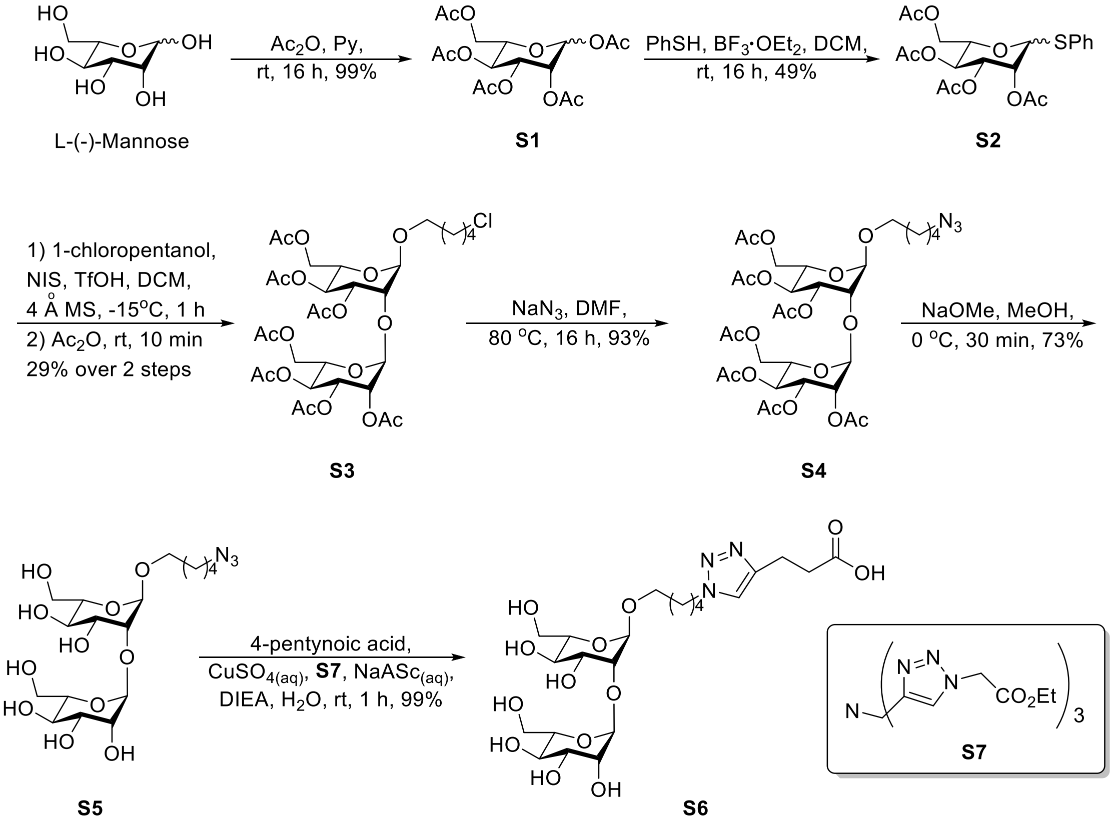

**1,2,3,4,6-Penta-*O*-acetyl-L-mannopyranose (S1)**

To a solution of L-(-)-mannose (0.20 g, 1.10 mmol) in pyridine (2.0 ml) was added acetic anhydride (1.6 ml, 16.50 mmol) in an ice bath. After 10 min, the ice bath was removed, and the reaction mixture was stirred under an argon atmosphere at room temperature for 16 h. The reaction was quenched by the addition of methanol in an ice bath, and the solvent was removed under reduced pressure. The residue was diluted with water and extracted with DCM. The combined organic layers were washed with 1N HCl (aq) and brine, dried over $Na_2SO_2$, filtered, and concentrated under reduced pressure to give compound **S1** (0.43 g, 99%) as a mixture of α/β pyranose anomers (3:1). The product was used for the next step without further purification. $^1$H NMR (400 MHz, $CDCl_3$): δ 6.05 (d, *J* = 1.9 Hz, 1H, H-1α), 5.35–5.31 (m, 2H, H-3α, H-4α),

5.26-5.22 (m, 1H, H-2α), 4.30-4.23 (m, 1H, H-6′α), 4.13–4.00 (m, 2H, H-6, H-5), 2.19 (s, 3H, $COCH_3$), 2.15 (s, 3H, $COCH_3$), 2.06 (s, 3H, $COCH_3$), 2.02 (s, 3H, $COCH_3$), 1.98 (s, 3H, $COCH_3$) ppm.

**Phenyl 2,3,4,6-tetra-*O*-acetyl-1-thio-L-mannopyranoside (S2)**

To a solution of compound **S1** (0.43 g, 1.10 mmol) in DCM (5.5 ml) was added a solution of thiophenol (PhSH, 0.17 ml, 1.66 mmol) in DCM (5.5 ml) via syringe. The reaction mixture was cooled in an ice bath, and $BF_3{\cdot}OEt_2$ (0.42 mL, 3.33 mmol) was added dropwise. The ice bath was then removed, and the reaction mixture was stirred at room temperature for 16 h. The reaction mixture was then diluted with water and extracted with DCM. The combined organic layers were washed with saturated aqueous $NaHCO_3$ solution and brine, dried over $Na_2SO_4$ and concentrated under reduced pressure. The residue was purified by column chromatography (EtOAc/hexanes = 1:2 to 1:1) to afford compound **S2** (0.32 g, 49%) as a mixture of α/β pyranose anomers (10:1). $^1$H NMR (400 MHz, $CDCl_3$) δ 7.48-7.45 (m, 2H, H-arom.), 7.32-7.27 (m, 3H, H-arom.), 5.48-5.47 (m, 2H, H-1, H-2), 5.32-5.28 (m, 2H, H-3, H-4), 4.54-4.50 (m, 1H, H-5), 4.28 (dd, $J$ = 12.2, 5.9 Hz, 1H, H-6), 4.08 (dd, J = 12.2, 2.4 Hz, 1H, H-6′), 2.13 (s, 3H, $COCH_3$), 2.05 (s, 3H, $COCH_3$), 2.03 (s, 3H, $COCH_3$), 2.00 (s, 3H, $COCH_3$).

**5-Chloropentyl 2,3,4,6-tetra-*O*-acetyl-α-L-mannopyranosyl-(1→2)-3,4,6-tri-*O*-acetyl-α-L-mannopyranoside (S3)**

To a solution of Compound **S2** (0.1 g, 0.23 mmol, 1.0 equiv.) and 1-chloropentanol (28.4 μl, 0.24 mmol, 1.05 equiv.) in DCM (1.2 ml) were stirred at room temperature in the presence of 4 Å MS (0.1 g) for 30 min. After cooling to -15 °C, N-iodosuccinimide (63.1 mg, 0.28 mmol, 1.2 equiv.) and trifluoromethanesulfonic acid (4.1 μl, 0.05 mmol, 0.2 equiv) were slowly added and stirring continued at –15 °C for 1 h. The reaction mixture was allowed to warm to room temperature and then acetic anhydride (55 μl, 2.5 equiv.) added. After 30 min, the molecular sieves were filtered through cotton and then washed with EtOAc. The filtrate was diluted with $H_2O$, extracted with EtOAc and washed with ice-cold 10% $Na_2S_2O_5$ (aq) and brine, the organic layer was dried over $Na_2SO_4$, filtered, and concentrated under reduced pressure. The crude product was purified by column chromatography (EtOAc/hexane = 1:3 to 1:1) to give compound **S3** (50.4 mg, 29%). $^1$H NMR (400 MHz, $CDCl_3$) δ 5.36 (dd, $J$ = 10.0, 3.4 Hz, 1H, H-3′), 5.30-5.20 (m, 4H, H-4, H-3, H-2′, H-4′), 4.88 (br s, 2H, H-1, H-1′), 4.21–4.05 (m, 5H, H-5, H-6a,b, H-6a′,b′), 3.98-3.97 (m, 1H, H-2), 3.88-3.85 (m, 1H, H-5′), 3.67 (td, $J$ = 8.0, 6.3 Hz, 1H, H-1b″), 3.51 (t, $J$ = 6.6 Hz, 2H, H-5″), 3.41 (td, $J$ = 8.0, 6.2 Hz, 1H, H-1a″), 2.10, 2.09, 2.04, 2.03, 2.00, 1.99, 1.96 (7 s, 21H, 7 × $COCH_3$), 1.79-1.72 (m, 2H, H-4″), 1.62-1.56 (m, 2H, H-2″), 1.50-1.45 (m, 2H, H-3″).

**5-Azidopentyl 2,3,4,6-tetra-*O*-acetyl-α-L-mannopyranosyl-(1→2)-3,4,6-tri-*O*-acetyl-α-L-mannopyranoside (S4)**

To a solution of compound **S3** (50.4 mg, 0.07 mmol) in anhydrous DMF (1.0 mL) was added sodium azide (6.8 mg, 0.11 mmol). The reaction mixture was stirred under an argon atmosphere at 80 °C for 16 h. After cooling to room temperature, the reaction mixture was diluted with water and extracted with Ethyl ether. The combined organic layers were washed with water and brine, dried over $Na_2SO_4$, filtered, and concentrated under reduced pressure to give compound **S4** (47.5 mg, 93%). The product was used for the next step without further purification. $^1$H NMR (400 MHz, $CDCl_3$) δ 5.36 (dd, $J$ = 10.0, 3.4 Hz, 1H, H-3′), 5.31-5.21 (m, 4H, H-4, H-3, H-2′, H-4′),

4.88 (br s, 2H, H-1, H-1′), 4.20–4.06 (m, 5H, H-5, H-6a,b, H-6a′,b′), 3.98-3.97 (m, 1H, H-2), 3.88-3.84 (m, 1H, H-5′), 3.67 (td, *J* = 8.1, 6.4 Hz, 1H, H-1b″), 3.40 (td, *J* = 8.0, 6.3 Hz, 1H, H-1a″), 3.25 (t, *J* = 6.7 Hz, 2H, H-5″), 2.11, 2.09, 2.04, 2.00, 1.99, 1.96 (6 s, 21H, 7 × $COCH_3$), 1.62-1.55 (m, 4H, H-4″, H-2″), 1.44-1.38 (m, 2H, H-3″).

**5-Azidopentyl α-L-mannopyranosyl-(1→2)-α-L-mannopyranoside (S5)**
To a solution of compound **S4** (47.5 mg, 0.06 mmol) in anhydrous MeOH (0.6 mL) at 0 °C was added NaOMe in MeOH (0.5 M, 25.4 µl, 0.01 mmol). The reaction mixture was stirred at 0 °C until completion. The reaction was neutralized with 1 N HCl (aq), and the solvent was removed under reduced pressure. The resulting crude was dissolved in water and acetonitrile, followed by centrifugation, and purified by semi-preparative HPLC using an Agilent 1200 or 1260 Infinity system equipped with a Zorbax C8 column (Zorbax Rx, 9.4 mm × 250 mm). Solvent A was $H_2O$, and solvent B was ACN. The purification was performed at a flow rate of 3 mL/min with a gradient of 5–90% B over 30 min. The product-containing fractions were collected and lyophilized to give compound **S5** (21.0 mg, 73%). $[\alpha]^{25}\_D$ = -50.1 (c 0.38, $CD_3OD$). IR (ATR) νmax 3406, 2930, 2866, 2099, 1457, 1132, 1061, 1030 $cm^{-1}$. $^1H$ NMR (400 MHz, $CD_3OD$) δ ppm 5.00 (s, 1H, H-1), 4.91 (d, *J* = 1.64 Hz, 1H, H-1′), 3.92 (dd, *J* = 3.2, 1.8 Hz, 1H, H-2′), 3.81-3.79 (m, 1H, H-2), 3.78 – 3.74 (m, 3H, H-6a′,b′, H-3), 3.70-3.59 (m, 5H, H-3′, H-5′, H-6a, H-1″), 3.55-3.37 (m, 4H, H-6b, H-4, H-4′, H-5), 3.26-3.23 (m, 2H, H-5″), 1.61-1.53 (m, 4H, H-2″, H-4″), 1.45-1.39 (m, 2H, H-3″); $^{13}C$ NMR (400 MHz, $CD_3OD$) δ ppm 104.17, 99.90, 80.66, 74.98, 74.64, 72.41, 72.19, 71.85, 69.04, 68.82, 68.37, 63.13, 63.05, 52.36, 30.10, 29.67, 24.56; HRMS (ESI) m/z: $[M + H]^+$ calcd for $C_{17}H_{31}N_3O_{11}$ 454.2031, found 454.2023.

**5-[4-(2-Carboxyethyl)-1H-1,2,3-triazol-1-yl]pentyl α-L-mannopyranosyl-(1→2)-α-L-mannopyranoside (S6)**
To a solution of compound **S5** (3.4 mg, 7.50 µmol) and 4-pentynoic acid (0.9 mg, 6.3 µmol) in $H_2O$ (1.4 ml) was added a premixed solution containing $CuSO_4$ (aq) (0.1 equiv., 40 mM), tris(triazolyl)amine ligand **S7** (0.1 equiv., 40 mM in DMSO), and sodium ascorbate (aq) (2.0 equiv., 800 mM). The reaction mixture was stirred at room temperature for 1 h. The resulting crude was dissolved in water and acetonitrile, followed by centrifugation, and purified by semi-preparative HPLC using an Agilent 1200 or 1260 Infinity system equipped with a Zorbax C8 column (Zorbax Rx, 9.4 mm × 250 mm). Solvent A was $H_2O$, and solvent B was ACN. The purification was performed at a flow rate of 3 mL/min with a gradient of 5–90% B over 50 min. The product-containing fractions were collected and lyophilized to give compound **S6** (4.1 mg, 99%). $[\alpha]^{25}\_D$ = -34.3 (c 0.19, $D_2O$). IR (ATR) νmax 3377, 3099, 2937, 1682, 1557, 1417, 1204, 1131, 1060, 1031 $cm^{-1}$. $^1H$ NMR (400 MHz, $D_2O$) δ ppm 7.84 (s, 1H, =CH), 5.08 (s, 1H, H-1), 5.03 (d, *J* = 1.04 Hz, 1H, H-1′), 4.42 (t, 2H, *J* = 6.74 Hz, H-5″), 4.09-4.08 (m, 1H, H-2′), 3.92-3.49 (m, 13H, H-2, H-6a′,b′, H-3, H-3′, H-5′, H-6a, H-1″, H-6b, H-4, H-4′, H-5), 3.03 (t, 2H, *J* = 7.10 Hz, H-1‴), 2.77 (t, 2H, *J* = 7.10 Hz, H-2‴), 1.96-1.89 (m, 2H, H-4″), 1.65-1.57 (m, 2H, H-2″), 1.37-1.27 (m, 2H, H-3″); $^{13}C$ NMR (400 MHz, $D_2O$) δ ppm 102.36, 98.12, 78.76, 73.29, 72.78, 70.34, 69.97, 67.58, 66.98, 66.94, 61.15, 60.93, 50.30, 28.99, 27.79, 22.31, 20.38; HRMS (ESI) m/z: $[M + H]^+$ calcd for $C_{22}H_{37}N_3O_{13}$ 552.2399, found 552.2365.

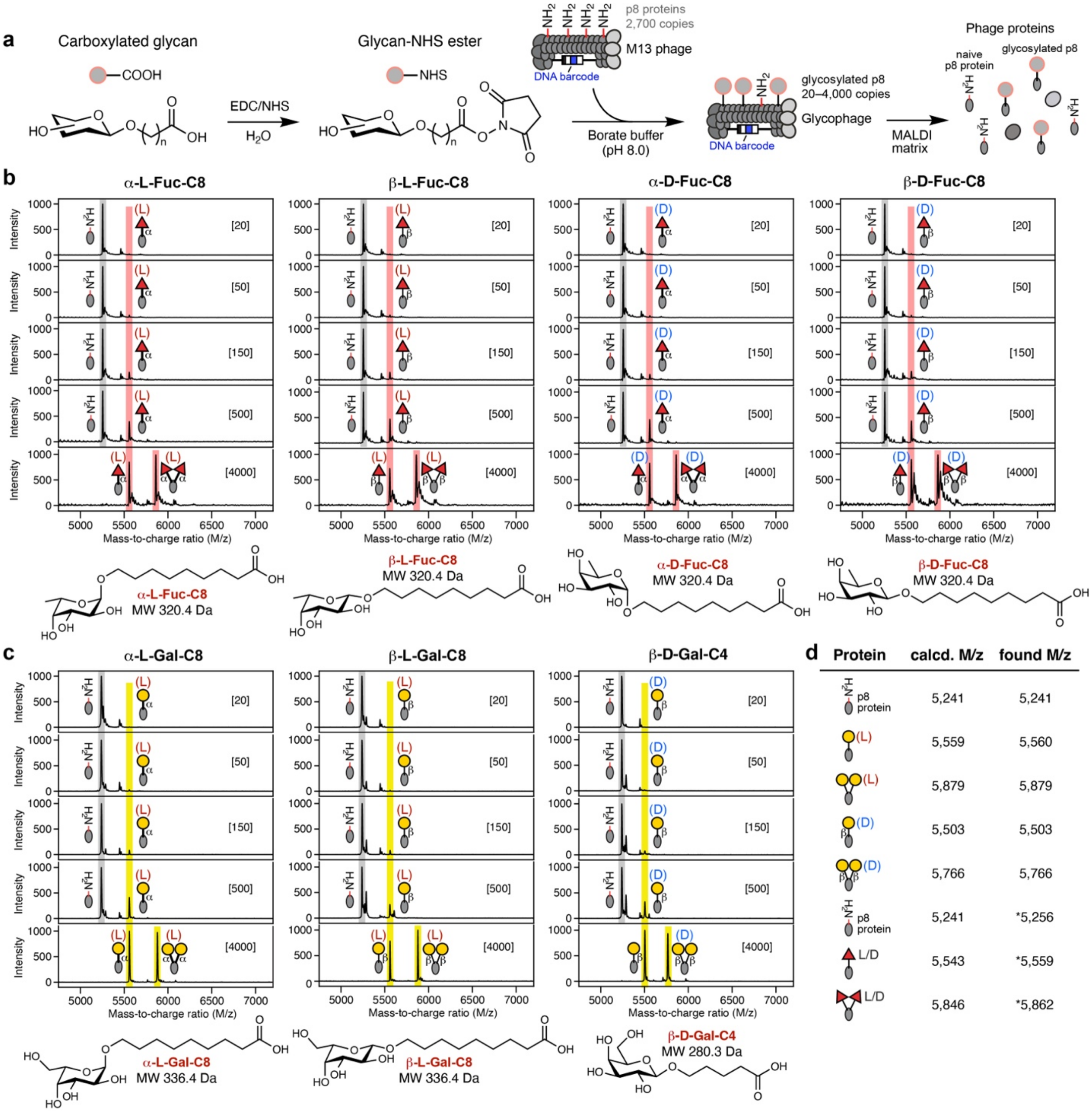

**Figure S1:** MALDI-TOF MS analysis of glycosylated phages used in the tested LiGA.
(a) Schematic representation of silent-barcoded glycosylated phage generation using EDC/NHS-mediated coupling. (b) MALDI-TOF MS characterization of enantiomeric fucose-phage conjugates at different glycan displaying densities. (c) MALDI-TOF MS characterization of enantiomeric galactose-phage conjugates. (d) Table of observed and calculated M/z.

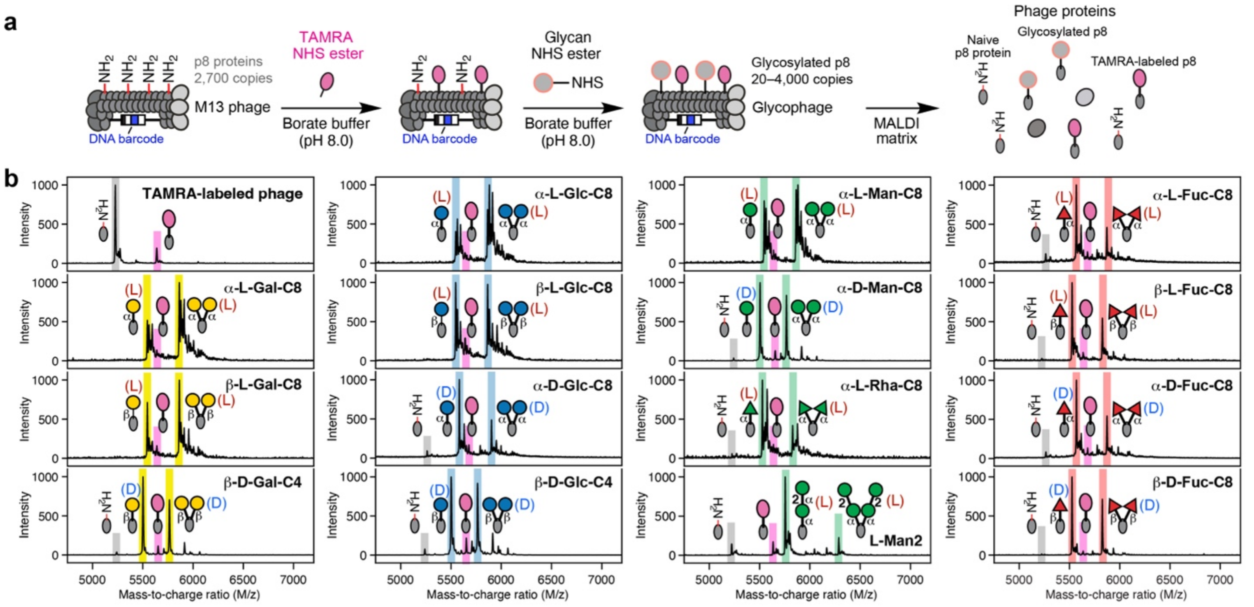

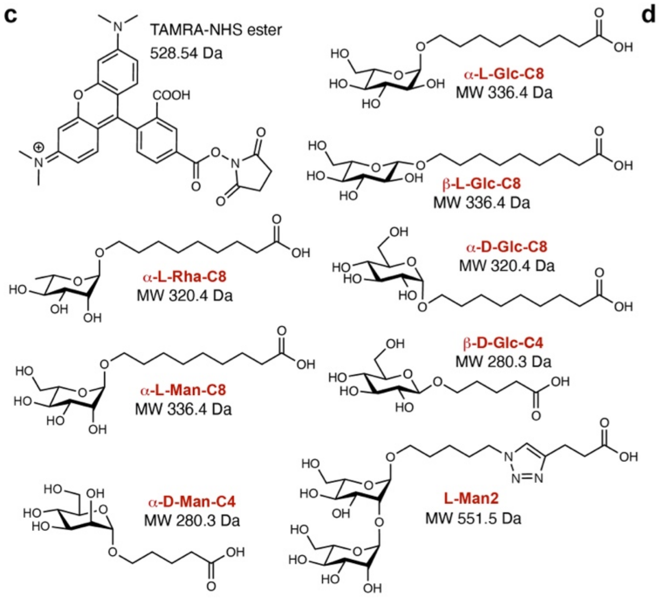

**d**

| Protein | calcd. M/z | found M/z | Protein | calcd. M/z | found M/z |
|---|---|---|---|---|---|
| p8 protein | 5,241 | 5,228 | (L) α | 5,559 | 5,544 |
| TAMRA-labeled p8 | 5,752 | 5,640 | (L) α α | 5,878 | 5,863 |
| (L) α/β | 5,559 | 5,546 | (D) α | 5,503 | 5,505 |
| (L) α/β | 5,878 | 5,863 | (D) α α | 5,766 | 5,767 |
| (D) β | 5,503 | 5,502 | (L) α | 5,543 | 5,527 |
| (D) β β | 5,766 | 5,764 | (L) α α | 5,846 | 5,828 |
| (L) α/β | 5,559 | 5,546 | 2 α (L) α | 5,775 | 5,755 |
| (L) α/β | 5,878 | 5,864 | 2 2 (L) α α | 6,308 | 6,288 |
| (D) α | 5,559 | *5,583 | L/D α/β | 5,543 | 5,525 |
| (D) α α | 5,878 | *5,903 | L/D α/β | 5,846 | 5,826 |
| (D) β | 5,503 | 5,503 | | | |
| (D) β β | 5,766 | 5,766 | | | |

**Figure S2:** MALDI-TOF MS analysis of fluorescent labeled glycosylated phages.
(a) Schematic representation of the production of fluorescent glycophages for use in Lectin Microarray assays. (b) MALDI-TOF MS characterization of enantiomeric Gal-, Glc-, Man- and Fuc-phage conjugates. (c) Structure detail of chemicals used in fluorescent labeling of glycophages. (d) Table of observed and calculated M/z.

| Common name | Displaying density (# of glycans per phage) | Chemical structure |
|---|---|---|
| a-GalNAc | [135], [540], [1350] | GalNAc(α-O-butyl-COOH |
| b-GalNAcAz | [10], [20], [50], [100], [500], [1000] | GalNAc(β-P3 |
| b-GalNAc | [10], [20], [50], [100], [500], [1000] | GalNAc(β-O-$PEG_3$-COOH |
| Tri-GalNAc | [100], [500], [1000] | $(β-D-GalNAc-sp)_3$-NHCO-$PEG_5$-COOH |
| GN | [1050] | GlcNAc(β-Sp |
| Lac-peg4 | [1080] | Galβ1-4Glc(β-P4 |
| LacNAc, LN | [970] | Galβ1-4GlcNAc(β-Sp |
| Lec | [680] | Galβ1-3GlcNAc(β-Sp |
| Galili-tri | [1000] | Galα1-3Galβ1-4Glc(β-Sp |
| Pk | [860] | Galα1-4Galβ1-4Glc(β-Sp |
| Gala3-type1 | [350] | Galα1-3Galβ1-3GlcNAc(β-Sp |
| B2 tri | [350] | Galα1-3Galβ1-4GlcNAc(β-Sp |
| P1 tri | [620] | Galα1-4Galβ1-4GlcNAc(β-Sp |
| LacDiNAc | [50] | GalNAcβ1-4GlcNAc(β-Sp |
| LNT-2 | [430] | GlcNAcβ1-3Galβ1-4Glc(β-Sp |
| GNLN | [810] | GlcNAcβ1-3Galβ1-4GlcNAc(β-Sp |
| 3'GN type1 | [860] | GlcNAcβ1-3Galβ1-3GlcNAc(β-Sp |
| LNnT | [240] | Galβ1-4GlcNAcβ1-3Galβ1-4Glc(β-Sp |
| Globoside-P | [730], [1030] | GalNAcβ1-3Galα1-4Galβ1-4Glc(β-Sp |
| P1 tetra | [970] | GalNAcβ1-3Galα1-4Galβ1-4GlcNAc(β-Sp |
| P1 penta | [620] | Galα1-4Galβ1-4GlcNAcβ1-3Galβ1-4Glc(β-Sp |
| P1x penta | [620] | Galβ1-3GalNAcβ1-3Galα1-4Galβ1-4GlcNAc(β-Sp |
| Tri-LN | [380] | Galβ1-4GlcNAcβ1-3Galβ1-4GlcNAcβ1-3Galβ1-4 GlcNAcβ1-3(β-Sp |
| Di-N3 | [970] | Fucα1-2Gal(β-Sp |
| H type 1 | [700] | Fucα1-2Galβ1-3GlcNAc(β-Sp |
| H type 1 | [135], [1350] | Fucα1-2Galβ1-3GlcNAc(β-X-COOH |
| H type 2 | [540] | Fucα1-2Galβ1-4GlcNAc(β-Sp |
| H type 2 | [135], [540], [1350] | Fucα1-2Galβ1-4GlcNAc(β-X-COOH |
| H type 3 | [100], [500], [1000] | Fucα1-2Galβ1-3GalNAc(α-X-COOH |
| H type 4 | [100], [500], [1000] | Fucα1-2Galβ1-3GalNAc(β-X-COOH |
| H type 5 | [100], [500], [1000] | Fucα1-2Galβ1-3Gal(β-X-COOH |
| 2'FL | [950] | Fucα1-2Galβ1-4Glc(β-Sp |
| H type 6 | [100], [500], [1000] | Fucα1-2Galβ1-4Glc(β-X-COOH |
| A tetra type 1 | [700] | GalNAcα1-3[Fucα1-2]Galβ1-3GlcNAc(β-Sp |
| A type 1 | [135], [540], [1350] | GalNAcα1-3[Fucα1-2]Galβ1-3GlcNAc(β-X-COOH |
| A tetra type 2 | [920] | GalNAcα1-3[Fucα1-2]Galβ1-4GlcNAc(β-Sp |
| A type 2 | [135], [540], [1350] | GalNAcα1-3[Fucα1-2]Galβ1-4GlcNAc(β-X-COOH |
| A type 3 | [135], [540], [1350] | GalNAcα1-3[Fucα1-2]Galβ1-3GalNAc(α-X-COOH |
| A type 4 | [100], [500], [1000] | GalNAcα1-3[Fucα1-2]Galβ1-3GalNAc(β-X-COOH |
| A type 5 | [135], [540], [1350] | GalNAcα1-3[Fucα1-2]Galβ1-4Gal(β-X-COOH |
| A tetra type 6 | [590] | GalNAcα1-3[Fucα1-2]Galβ1-4Glc(β-Sp |
| A type 6 | [135], [540], [1350] | GalNAcα1-3[Fucα1-2]Galβ1-4Glc(β-X-COOH |
| B tetra type 1 | [620] | Galα1-3[Fucα1-2]Galβ1-3GlcNAc(β-Sp |
| B type 1 | [135], [540], [1350] | Galα1-3[Fucα1-2]Galβ1-3GlcNAc(β-X-COOH |
| B tetra type 2 | [970] | Galα1-3[Fucα1-2]Galβ1-4GlcNAc(β-Sp |
| B type 2 | [540] | Galα1-3[Fucα1-2]Galβ1-4GlcNAc(β-X-COOH |
| B type 3 | [100], [500], [1000] | Galα1-3[Fucα1-2]Galβ1-3GalNAc(α-X-COOH |
| B type 4 | [135], [540], [1350] | Galα1-3[Fucα1-2]Galβ1-3GalNAc(β-X-COOH |
| B type 5 | [540], [1350] | Galα1-3[Fucα1-2]Galβ1-3Gal(β-X-COOH |

| | | |
|---|---|---|
| B tetra type 6 | [920] | Galα1-3[Fucα1-2]Galβ1-4Glc(β-Sp |
| B type 6 | [135], [540], [1350] | Galα1-3[Fucα1-2]Galβ1-4Glc(β-X-COOH |
| 2'F-B type 2 | [220], [760] | Galα1-3[Fucα1-2]Galβ1-4[Fucα1-3]GlcNAc(β-Sp |
| H2 | [430] | Fucα1-2Galβ1-4GlcNAcβ1-3Galβ1-4GlcNAc(β-Sp |
| H3 | [190] | Fucα1-2Galβ1-4GlcNAcβ1-3Galβ1-4GlcNAcβ1-3 Galβ1-4GlcNAc(β-Sp |
| Tri-AN3 | [1080] | GalNAcα1-3[Fucα1-2]Gal(β-Sp |
| LeA | [950] | Galβ1-3[Fucα1-4]GlcNAc(β-Sp |
| Lex | [810] | Galβ1-4[Fucα1-3]GlcNAc(β-Sp |
| Gala3Lex | [620] | Galα1-3Galβ1-4[Fucα1-3]GlcNAc(β-Sp |
| Di-Lex | [410] | Galβ1-4[Fucα1-3]GlcNAcβ1-3Galβ1-4[Fucα1-3] GlcNAcβ1-Sp |
| Ley-Lex | [350] | Fucα1-2Galβ1-4[Fucα1-3]GlcNAcβ1-3Galβ1-4[Fucα1-3] GlcNAc(β-Sp |
| LeALex | [350] | Galβ1-3[Fucα1-4]GlcNAcβ1-3Galβ1-4[Fucα1-3] GlcNAc(β-Sp |
| Lex-LeA | [410] | Galβ1-4[Fucα1-3]GlcNAcβ1-3Galβ1-3[Fucα1-4] GlcNAc(β-Sp |
| Lec-LeX | [570] | Galβ1-3GlcNAcβ1-3Galβ1-4[Fucα1-3]GlcNAc(β-Sp |
| Tri-Lex | [430] | Galβ1-4[Fucα1-3]GlcNAcβ1-3Galβ1-4[Fucα1-3] GlcNAcβ1-3Galβ1-4[Fucα1-3]GlcNAc(β-Sp |
| Ley-Di-Lex | [510] | Fucα1-2Galβ1-4[Fucα1-3]GlcNAcβ1-3Galβ1-4[Fucα1-3]GlcNAcβ1-3Galβ1-4[Fucα1-3]GlcNAc(β-Sp |
| (Man)3 | [1300], [1730] | Manα1-6[Manα1-3]Man(α-S6 |
| 11 | [50], [150], [500], [750], [780] | Manα1-6[Manα1-3]Manβ1-4GlcNAcβ1-4GlcNAc(β1-Sp |
| 6 | [50], [150], [500], [750], [810], [1000] | GlcNAcβ1-2Manα1-6[GlcNAcβ1-2Manα1-3]Manβ1-4 GlcNAcβ1-4GlcNAc(β1-Sp |
| 10 | [150], [150], [500], [540], [1000] | Galβ1-4GlcNAcβ1-2Manα1-6[Galβ1-4GlcNAcβ1-2Manα1-3]Manβ1-4GlcNAcβ1-4GlcNAc(β1-Sp |
| 9 | [50], [510], [730], [950], [970] | Neu5Acα2-6Galβ1-4GlcNAcβ1-2Manα1-6[Neu5Acα2-6Galβ1-4GlcNAcβ1-2Manα1-3]Manβ1-4GlcNAcβ1-4 GlcNAc(β1-Sp |
| GM1 | [460] | Neu5Acα2-3[Galβ1-3GalNAcβ1-4]Galβ1-4Glc(β-Sp |
| GM2 | [190], [460], [590], [1190] | Neu5Acα2-3[GalNAcβ1-4]Galβ1-4Glc(β-Sp |
| CTISda | [350] | Neu5Acα2-3[GalNAcβ1-4]Galβ1-4GlcNAc(β-Sp |
| GD1a | [110] | Neu5Acα2-3[Neu5Acα2-3Galβ1-3GalNAcβ1-4]Galβ1-4 Glc(β-Sp |
| 3'S-Di-LeA | [160] | Neu5Acα2-3Galβ1-3[Fucα1-4]GlcNAcβ1-32(β-Sp |
| 3'SLeA-Lex | [570] | Neu5Acα2-3Galβ1-3[Fucα1-4]GlcNAcβ1-3Galβ1-4[Fucα1-3]GlcNAc(β-Sp |
| 3'S-Tri-LeX | [160] | Neu5Acα2-3Galβ1-4[Fucα1-3]GlcNAcβ1-3Galβ1-4[Fucα1-3]GlcNAcβ1-3Galβ1-4[Fucα1-3]GlcNAc(β-Sp |
| 3'Slec | [430] | Neu5Acα2-3Galβ1-3GlcNAc(β-Sp |
| GM3 | [190], [540], [1190] | Neu5Acα2-3Galβ1-4Glc(β-Sp |
| 3'SLDN | [460] | Neu5Acα2-3GalNAcβ1-4GlcNAc(β-Sp |
| 3'S-Di-Lec | [270] | Neu5Acα2-3Galβ1-3GlcNAcβ1-3Galβ1-3GlcNAc(β-Sp |
| 3'SLecLN | [860] | Neu5Acα2-3Galβ1-3GlcNAcβ1-3Galβ1-4GlcNAc(β-Sp |
| 3'SLN-Lec | [410] | Neu5Acα2-3Galβ1-4GlcNAcβ1-3Galβ1-3GlcNAc(β-Sp |
| 3'S-Di-LN | [1350] | Neu5Acα2-3Galβ1-4GlcNAcβ1-3Galβ1-4GlcNAcβ1-Sp |
| 3'STri-LN | [160] | Neu5Acα2-3Galβ1-4GlcNAcβ1-3Galβ1-4GlcNAcβ1-3Galβ1-4GlcNAc(β-Sp |
| 3'SLec (Gc) | [350] | Neu5Gcα2-3Galβ1-3GlcNAc(β-Sp |
| 3'SL (Gc) | [490] | Neu5Gcα2-3Galβ1-4Glc(β-Sp |
| 3'SLN (Gc) | [570] | Neu5Gcα2-3Galβ1-4GlcNAc(β-Sp |
| 3'-KDNLec | [760] | Kdnα2-3Galβ1-3GlcNAc(β-Sp |

| | | |
|---|---|---|
| 3'KDNLN | [510] | Kdnα2-3Galβ1-4GlcNAc(β-Sp |
| 6'SL | [620] | Neu5Acα2-6Galβ1-4Glc(β-Sp |
| 6'SLN (Gc) | [430] | Neu5Gcα2-6Galβ1-4GlcNAc(β-Sp |
| 6'SLDN | [620] | Neu5Acα2-6GalNAcβ1-4GlcNAc(β-Sp |
| 6'SLN-Lec | [300] | Neu5Acα2-6Galβ1-4GlcNAcβ1-3Galβ1-3GlcNAc(β-Sp |
| 6'S-Di-LN | [300] | Neu5Acα2-6Galβ1-4GlcNAcβ1-3Galβ1-4GlcNAcβ1-Sp |
| GD2 | [460] | Neu5Acα2-8Neu5Acα2-3[GalNAcβ1-4]Galβ1-4Glc(β-Sp |
| GD3 | [320] | Neu5Acα2-8Neu5Acα2-3Galβ1-4Glc(β-Sp |
| GT2 | [160] | Neu5Acα2-8Neu5Acα2-8Neu5Acα2-3[GalNAcβ1-4] Galβ1-4Glc(β-Sp |
| GT3 | [510] | Neu5Acα2-8Neu5Acα2-8Neu5Acα2-3Galβ1-4Glc(β-Sp |
| GQ2 | [140] | Neu5Acα2-8Neu5Acα2-8Neu5Acα2-8Neu5Acα2-3 [GalNAcβ1-4]Galβ1-4Glc(β-Sp |
| (Galf)4 | [<8] | Gal*f* β1-5Gal*f* β1-5Gal*f* β1-5Gal*f* (β-S8 |
| (Man)4 | [20], [100], [300], [720], [740] | Manα1-2Manα1-2Manα1-3Man(α-S5 |
| (L-Man)2 | [500] | L-Manα1-2-L-Man(α-O-$(CH_2)_5$-triazole-$(CH_2)_2$-COOH |
| α-L-Rha | [20], [50], [150], [500] | L-Rha(α-Sp |
| α-L-Glc | [25], [50], [150], [500], [4000] | L-Glc(α-O-octyl-COOH |
| β-L-Glc | [25], [50], [150], [500], [4000] | L-Glc(β-O-octyl-COOH |
| β-D-Glc | [25], [50], [150], [500], [4000] | D-Glc(β-O-butyl-COOH |
| α-L-Gal | [25], [50], [150], [500], [4000] | L-Gal(α-O-octyl-COOH |
| β-L-Gal | [25], [50], [150], [500], [4000] | L-Gal(β-O-octyl-COOH |
| β-D-Gal | [25], [50], [150], [500], [4000] | D-Gal(β-O-butyl-COOH |
| α-L-Man | [25], [50], [150], [500], [4000] | L-Man(α-O-octyl-COOH |
| α-D-Man | [25], [50], [150], [500], [4000] | D-Man(α-O-butyl-COOH |
| α-L-Fuc | [25], [50], [150], [500], [4000] | L-Fuc(α-O-octyl-COOH |
| β-L-Fuc | [25], [50], [150], [500], [4000] | L-Fuc(β-O-octyl-COOH |
| α-D-Fuc | [25], [50], [150], [500], [4000] | D-Fuc(α-O-octyl-COOH |
| β-D-Fuc | [25], [50], [150], [500], [4000] | D-Fuc(β-O-octyl-COOH |
| AzOH | [100], [500] | 2-Azidoethanol |
| blank | [0] | Unmodified M13 phage |

**Table S1:** Chemical composition of the tested mirror-LiGA.
Linker abbreviations: Sp: -O-$(CH_2)_2$-$N_3$; S5: -O-$(CH_2)_5$-$N_3$; S6: -O-$(CH_2)_6$-$N_3$; S8: -O-$(CH_2)_8$-$N_3$; P3: -(O-$CH_2$-$CH_2$)$_3$-$N_3$; P4: -(O-$CH_2$-$CH_2$)$_4$-$N_3$; and -X-: -O-$(CH_2)_8$-S-$(CH_2)_2$-NHCO-$(CH_2)_4$-. Enantiomeric glycan-phage constructs were highlighted in pink. Conjugation of azido glycans to phage was performed as follows. Phages were first acylated with DBCO-NHS ester (Cayman Chemical, 40461) in 1:1 (v/v) PBS:glycerol, purified using Zeba spin desalting columns (7.0 kDa MWCO), and then conjugated to azido glycans by strain-promoted azide-alkyne cycloaddition (SPAAC). Carboxylate-containing glycans were conjugated to phage by activating the glycan carboxylate with EDC hydrochloride and NHS (1 equiv. each, relative to the glycan) in water, followed by amide coupling of the resulting glycan-NHS ester to phage in borate buffer (100 mM borate, 150 mM NaCl, pH 8.0).

| LiGA | Target | Dataset | n | Sample type | Comments |
|---|---|---|---|---|---|
| NOA | AAL | 20260205-87NOAaalPXA-EG | 4 | Purified lectin | **Figure 2** & **Extended Data Figure 1** |
| NOA | AOL* | 20260212-87NOAaolPXA-EG | 4 | Purified lectin | (measured on Corning Easy-wash |
| NOA | UEA-I | 20260212-87NOAueaPXA-EG | 4 | Purified lectin | High binding 96 well-plate) |
| NOA | RSL | 20260212-87NOArslPXA-EG | 4 | Purified lectin | |
| NOA | MGL* | 20260212-87NOAvaaaCYC-EG | 3 | Purified lectin | |
| NOA | SBA | 20260205-87NOAzfaPXA-EG | 4 | Purified lectin | |
| NOA | ECL | 20260212-87NOAeaaPXA-EG | 4 | Purified lectin | |
| NOA | PNA | 20260212-87NOAyfaPXA-EG | 4 | Purified lectin | |
| NOA | ASGPR1 | 20260402-87NOAasgPAN-IF | 4 | Purified lectin | |
| NOA | WGA | 20260212-87NOAwgalPXA-EG | 4 | Purified lectin | |
| NOA | ConA | 20260205-87NOAcaPXA-EG | 4 | Purified lectin | |
| NOA | Griffithsin | 20260430-87NOAowaPXA-EG | 4 | Purified lectin | |
| NOA | BanLec$^{H84T}$ | 20260430-87NOAnwaPXA-EG | 4 | Purified lectin | |
| NOA | BanLec$^{H84T}$* | 20260205-87NOAkqaCYC-EG | 3 | Purified lectin | |
| NOA | SP-D* | 20260212-87NOAvbaaCYC-EG | 3 | Purified lectin | |
| NOA | MBL* | 20260212-87NOAhaaaCYC-EG | 3 | Purified lectin | |
| NOA | DC-SIGN$^{WT}$ | 20260212-87NOAdcPXA-EG | 4 | Purified lectin | |
| NOA | DC-SIGN$^{T314A}$ | 20260212-87NOAdctPXA-EG | 4 | Purified lectin | |
| NOA | DC-SIGN$^{M270F}$ | 20260212-87NOAdcmPXA-EG | 4 | Purified lectin | |
| NOA | Galectin-1* | 20260212-87NOAoqaCYC-EG | 3 | Purified lectin | |
| NOA | Galectin-1 | 20260529-87NOAgoULC-EG | 4 | Purified lectin | |
| NOA | Galectin-2 | 20260529-87NOAgadULC-EG | 4 | Purified lectin | |
| NOA | Galectin-3 | 20260529-87NOAgaULC-EG | 4 | Purified lectin | |
| NOA | Galectin-7 | 20260529-87NOAgsULC-EG | 4 | Purified lectin | |
| NOA | Galectin-8* | 20260212-87NOAqaaaCYC-EG | 3 | Purified lectin | |
| NOA | Galectin-8A | 20260529-87NOAgeiULC-EG | 4 | Purified lectin | |
| NOA | Galectin-8B | 20260529-87NOAgebULC-EG | 4 | Purified lectin | |
| NOA | Galectin-10 | 20260529-87NOAgaeULC-EG | 4 | Purified lectin | |
| NOA | Galectin-14 | 20260529-87NOAghfULC-EG | 4 | Purified lectin | |
| NOA | hItln1* | 20260212-87NOAgbaaCYC-EG | 3 | Purified lectin | |
| NOA | REG4* | 20260212-87NOAzaaaCYC-EG | 3 | Purified lectin | |
| NOA | CL-K1* | 20260212-87NOAiaaaCYC-EG | 3 | Purified lectin | |
| NOA | CRP* | 20260212-87NOAfbaaCYC-EG | 3 | Purified lectin | |
| NOA | diCBM40 | 20260430-87NOAcjaPXA-EG | 3 | Purified lectin | |
| NOA | SNA-I | 20260212-87NOAsnaPXA-EG | 4 | Purified lectin | |
| NOA | Siglec-1$^{WT}$ | 20260212-87NOAbiaPXA-EG | 4 | Purified lectin | |
| NOA | Siglec-1R | 20260212-87NOAhqaPXA-EG | 4 | Purified lectin | |
| NOA | Siglec-2$^{WT}$ | 20260212-87NOAsigaPXA-EG | 3 | Purified lectin | |
| NOA | Siglec-2R | 20260212-87NOAhqaPXA-EG | 4 | Purified lectin | |
| NOA | Siglec-3$^{WT}$ | 20260212-87NOAqsaPXA-EG | 4 | Purified lectin | |
| NOA | Siglec-4$^{WT}$ | 20260212-87NOAksaPXA-EG | 4 | Purified lectin | |
| NOA | Siglec-7$^{WT}$ | 20260212-87NOAsigbPXA-EG | 3 | Purified lectin | |
| NOA | Siglec-7R | 20260212-87NOAsigcPXA-EG | 4 | Purified lectin | |
| NOA | Siglec-15$^{WT}$* | 20260212-87NOAjqaCYC-EG | 3 | Purified lectin | |
| NOA | BSA | 20260205-87NOAbsPXA-EG | 4 | Purified protein | |
| NOA | No target | 20260212-87NOAooCYC-EG | 3 | Naïve LiGA | |
| NOA | hDC-SIGN$^{+}$ Raji | 20260306-87NOArjdNZC-FH | 3 | 0 mM D-Man | **Figure 3 & Extended Data Figure 4** |
| NOA | hDC-SIGN$^{+}$ Raji | 20260306-87NOArjdBZC-FH | 3 | 1 mM D-Man | (measured against cultured DC-SIGN+ |
| NOA | hDC-SIGN$^{+}$ Raji | 20260306-87NOArjdJZC-FH | 3 | 10 mM D-Man | Raji cells with gradient |
| NOA | hDC-SIGN$^{+}$ Raji | 20260306-87NOArjdIZC-FH | 3 | 25 mM D-Man | concentrations of free D-(+)- |
| NOA | hDC-SIGN$^{+}$ Raji | 20260306-87NOArjdZZC-FH | 3 | 50 mM D-Man | mannose) |
| NOA | hDC-SIGN$^{+}$ Raji | 20260306-87NOArjdCZC-FH | 3 | 100 mM D-Man | |
| NOA | Purified hDC-SIGN | 20260611-87NOAdcADCM-FH | 3 | 0 mM D-Man | **Extended Data Figure 4** (measured |
| NOA | Purified hDC-SIGN | 20260611-87NOAdcBDCM-FH | 3 | 1 mM D-Man | against purified DC-SIGN ECD-coated |
| NOA | Purified hDC-SIGN | 20260611-87NOAdcCDC-FH | 3 | 10mM D-Man | Corning Easy-wash High binding 96- |
| NOA | Purified hDC-SIGN | 20260611-87NOAdcDDCM-FH | 3 | 25 mM D-Man | well plate with gradient |
| NOA | Purified hDC-SIGN | 20260611-87NOAdcEDCM-FH | 3 | 50mM D-Man | concentrations of free D-(+)- |
| NOA | Purified hDC-SIGN | 20260611-87NOAdcFDCM-FH | 3 | 0.1 M D-Man | mannose) |
| NOA | Purified hDC-SIGN | 20260611-87NOAdcGDCM-FH | 3 | 1 M D-Man | |
| NOA | Purified hDC-SIGN | 20260514-87NOAdcACAA-FH | 3 | 0 mM L-Glc | **Extended Data Figure 4** (measured |
| NOA | Purified hDC-SIGN | 20260514-87NOAdcBBAA-FH | 3 | 1 mM L-Glc | on purified DC-SIGN ECD-coated |
| NOA | Purified hDC-SIGN | 20260514-87NOAdcCBAA-FH | 3 | 2 mM L-Glc | Corning Easy-wash High binding 96- |
| NOA | Purified hDC-SIGN | 20260514-87NOAdcDAAA-FH | 3 | 5 mM L-Glc | well plate and inhibited by gradient |
| NOA | Purified hDC-SIGN | 20260514-87NOAdcEBAA-FH | 3 | 10 mM L-Glc | concentrations of free L-(-)-glucose) |

| | | | | | |
|---|---|---|---|---|---|
| NOA | Purified hDC-SIGN | 20260514-87NOAdcFBAA-FH | 3 | 25 mM L-Glc | |
| NOA | Purified hDC-SIGN | 20260514-87NOAdcGBAA-FH | 3 | 50 mM L-Glc | |
| NOA | Purified hDC-SIGN | 20260514-87NOAdcHAAA-FH | 3 | 0.1 M L-Glc | |
| NOA | Purified hDC-SIGN | 20260514-87NOAdcIAAA-FH | 3 | 1 M L-Glc | |
| NOA | Donor# 655 | 20260212-87NOAdbPXA-GG | 3 | Human serum | **Figure 4** & **Extended Data Figure 2** (measured on anti-IgM precoated Corning Easy-wash High binding 96 well-plate) |
| NOA | Donor# 663 | 20260212-87NOAnhPXA-GG | 3 | Human serum | |
| NOA | Donor# 1471 | 20260320-87NOAlcPXA-GG | 3 | Human serum | |
| NOA | Donor# 844 | 20260212-87NOAjgPXA-GG | 3 | Human serum | |
| NOA | Donor# 788 | 20260320-87NOAdlPXA-GG | 3 | Human serum | |
| NOA | Donor# 794 | 20260212-87NOAenPXA-GG | 3 | Human serum | |
| NOA | Donor# 796 | 20260320-87NOAmmoPXA-GG | 3 | Human serum | |
| NOA | Donor# 1029 | 20260320-87NOAlePXA-GG | 3 | Human serum | |
| NOA | Donor# 1109 | 20260212-87NOArhPXA-GG | 3 | Human serum | |
| NOA | Donor# 1417 | 20260320-87NOAphPXA-GG | 3 | Human serum | |
| NOA | Donor# 1221 | 20260212-87NOArgPXA-GG | 3 | Human serum | |
| NOA | Donor# 791 | 20260320-87NOAifPXA-GG | 3 | Human serum | |
| NOA | Donor# 688 | 20260212-87NOAohPXA-GG | 3 | Human serum | |
| NOA | No target | 20260212-87NOAooBO-GG | 3 | Naïve LiGA | |
| NOA | Anti-Lewis A [7LE] | 20260402-87NOAalaPAN-IF | 4 | Purified Ab | **Figure 4** & **Extended Data Figure 2** (measured on Corning Easy-wash High binding 96 well-plate) |
| NOA | Anti-Lewis X [HI198] | 20260402-87NOAalxPAN-IF | 4 | Purified Ab | |
| NOA | Anti-Lewis Y [F3] | 20260402-87NOAalyPAN-IF | 4 | Purified Ab | |
| NOA | AntiH [BRIC39] | 20260421-87NOAfvaPAN-IF | 4 | Purified Ab | |
| NOA | AntiH [BRIC231] | 20260421-87NOAtvaPAN-IF | 4 | Purified Ab | |
| NOA | AntiH [BRIC198] | 20260430-87NOApvaPAN-IF | 4 | Purified Ab | |
| NOA | AntiH [17-206] | 20260402-87NOAanhPAN-IF | 4 | Purified Ab | |
| NOA | AntiH [17-206]. | 20260421-87NOAwvaPAN-IF | 4 | Purified Ab | |
| NOA | AntiH [A46BB10] | 20260402-87NOAyvaPAN-IF | 4 | Purified Ab | |
| NOA | AntiHo [89/8] | 20260402-87NOAjuaPAN-IF | 4 | Purified Ab | |
| NOA | AntiH [89-8] | 20260430-87NOArtaPAN-IF | 4 | Purified Ab | |
| NOA | AntiH [86-N] | 20260430-87NOAuoaPAN-IF | 4 | Purified Ab | |
| NOA | AntiH [97-1] | 20260430-87NOAgkaPAN-IF | 4 | Purified Ab | |
| NOA | AntiH [87-N] | 20260430-87NOAkwaPAN-IF | 4 | Purified Ab | |
| NOA | AntiA [HE24] | 20260402-87NOAhedPAN-IF | 4 | Purified Ab | |
| NOA | AntiA [HE193] | 20260402-87NOAvgPAN-IF | 4 | Purified Ab | |
| NOA | AntiA [HE193]. | 20260421-87NOAataPAN-IF | 4 | Purified Ab | |
| NOA | AntiA [NaM871F6] | 20260430-87NOAlqaPAN-IF | 4 | Purified Ab | |
| NOA | AntiA [BRIC145] | 20260430-87NOAxwaPAN-IF | 4 | Purified Ab | |
| NOA | AntiA [BRIC145]. | 20260421-87NOAbriPAN-IF | 4 | Purified Ab | |
| NOA | AntiA [NOVA] | 20260430-87NOAfwaPAN-IF | 4 | Purified Ab | |
| NOA | AntiA [3-3A] | 20260430-87NOAeuaPAN-IF | 4 | Purified Ab | |
| NOA | AntiA [Z2B1] | 20260421-87NOAzbbPAN-IF | 4 | Purified Ab | |
| NOA | AntiA [Z2A] | 20260430-87NOAduaPAN-IF | 4 | Purified Ab | |
| NOA | AntiA [9A] | 20260402-87NOAavaPAN-IF | 4 | Purified Ab | |
| NOA | AntiB [CLCP19B] | 20260402-87NOAvvaPAN-IF | 4 | Purified Ab | |
| NOA | AntiB [BRIC250] | 20260430-87NOAvwaPAN-IF | 4 | Purified Ab | |
| NOA | AntiB [ES4] | 20260430-87NOAsuaPAN-IF | 4 | Purified Ab | |
| NOA | AntiB [LB2] | 20260430-87NOAtwaPAN-IF | 4 | Purified Ab | |
| NOA | AntiB [NOVA] | 20260430-87NOAglaPAN-IF | 4 | Purified Ab | |
| NOA | AntiB [HEB29] | 20260402-87NOAndPAN-IF | 4 | Purified Ab | |
| NOA | AntiB [89-F] | 20260402-87NOAlpaPAN-IF | 4 | Purified Ab | |
| NOA | AntiB [BGRL1] | 20260402-87NOAkvaPAN-IF | 4 | Purified Ab | |
| NOA | AntiB [Z2H-2] | 20260402-87NOAzahPAN-IF | 4 | Purified Ab | |
| NOA | [86-M] | 20260421-87NOAetaPAN-IF | 4 | Purified Ab | |
| NOA | Anti-LNT [K21] | 20260430-87NOAywaPAN-IF | 4 | Purified Ab | |
| NOA | Anti-SialoTn | 20260430-87NOAqwaPAN-IF | 4 | Purified Ab | |
| NOA | [VK9] | 20260430-87NOAblaPAN-IF | 4 | Purified Ab | |
| NOA | [HE10] | 20260430-87NOAwwaPAN-IF | 4 | Purified Ab | |
| NOA | [JTL-2] | 20260430-87NOAjvaPAN-IF | 4 | Purified Ab | |
| NOA | [JTL-4] | 20260430-87NOAiuaPAN-IF | 4 | Purified Ab | |
| NOA | No target | 20251218-87NOAooNA-IF | 5 | Naïve LiGA | |
| NOA | F-CD4-Δ | 20260212-87NOAztaHYC-EG | 2 | FACS-sorted cells | (Human PBMCs derived primary immune cells) |
| NOA | M-CD4-Δ | 20260212-87NOAjsaHYC-EG | 2 | FACS-sorted cells | |
| NOA | F CD4-S | 20260212-87NOAztaUYC-EG | 2 | FACS-sorted cells | |
| NOA | M-CD4-S | 20260212-87NOAjsaUYC-EG | 2 | FACS-sorted cells | |
| NOA | F-CD8-Δ | 20260212-87NOAttaHYC-EG | 2 | FACS-sorted cells | |
| NOA | M-CD8-Δ | 20260212-87NOAytaHYC-EG | 2 | FACS-sorted cells | |

| | | | | | |
|---|---|---|---|---|---|
| NOA | F-CD8-S | 20260212-87NOAttaUYC-EG | 2 | FACS-sorted cells | |
| NOA | M-CD8-S | 20260212-87NOAttaUYC-EG | 2 | FACS-sorted cells | |
| NOA | F-MO-Δ | 20260212-87NOArnaHYC-EG | 2 | FACS-sorted cells | **Figure 5** (Human PBMCs derived primary immune cells) |
| NOA | M-MO-Δ | 20260212-87NOAqtaHYC-EG | 2 | FACS-sorted cells | |
| NOA | F-MO-S | 20260212-87NOArnaUYC-EG | 2 | FACS-sorted cells | |
| NOA | M-MO-S | 20260212-87NOAqtaUYC-EG | 2 | FACS-sorted cells | |
| NOA | F-B-Δ | 20260212-87NOApraHYC-EG | 2 | FACS-sorted cells | |
| NOA | M-B-Δ | 20260212-87NOAdqaHYC-EG | 2 | FACS-sorted cells | |
| NOA | F-B-S | 20260212-87NOApraUYC-EG | 2 | FACS-sorted cells | |
| NOA | M-B-S | 20260212-87NOAdqaUYC-EG | 2 | FACS-sorted cells | |
| NOA | F-NK-Δ | 20260212-87NOAxtaHYC-EG | 2 | FACS-sorted cells | |
| NOA | M-NK-Δ | 20260212-87NOAfsaHYC-EG | 2 | FACS-sorted cells | |
| NOA | F-NK-S | 20260212-87NOAxtaUYC-EG | 2 | FACS-sorted cells | |
| NOA | M-NK-S | 20260212-87NOAfsaUYC-EG | 2 | FACS-sorted cells | |
| NOA | F-Iso.CD4-Δ | 20260212-87NOAvtaQYC-EG | 2 | Isolated cells | |
| NOA | M-Iso.CD4-Δ | 20260212-87NOAgiaQYC-EG | 2 | Isolated cells | |
| NOA | F-Iso.CD4-S | 20260212-87NOAvtaTBC-EG | 2 | Isolated cells | |
| NOA | M-Iso.CD4-S | 20260212-87NOAgiaQYC-EG | 2 | Isolated cells | |
| NOA | F-Iso.CD8-Δ | 20260212-87NOAhtaHYC-EG | 2 | Isolated cells | |
| NOA | M-Iso.CD8-Δ | 20260212-87NOAiqaHYC-EG | 2 | Isolated cells | |
| NOA | F-Iso.CD8-S | 20260212-87NOAhtaTBC-EG | 2 | Isolated cells | |
| NOA | M-Iso.CD8-S | 20260212-87NOAiqaHYC-EG | 2 | Isolated cells | |
| MEG | Liver (in vivo) | 20241030-87MEGbiEYC-GL1 | 1 | Mouse #1 | **Figure 6** (mapping in vivo biodistribution of enantiomeric glycans in live mice) |
| MEG | Liver (in vivo) | 20241030-87MEGbiEYC-GL2 | 1 | Mouse #1 | |
| MEG | Liver (in vivo) | 20241030-87MEGbiEYC-GL3 | 1 | Mouse #2 | |
| MEG | Liver (in vivo) | 20241030-87MEGbiEYC-GL4 | 1 | Mouse #2 | |
| MEG | Liver (in vivo) | 20241030-87MEGbiEYC-GL5 | 1 | Mouse #3 | |
| MEG | Liver (in vivo) | 20241030-87MEGbiEYC-GL6 | 1 | Mouse #3 | |
| MOX | Liver (in vivo) | 20250707-87MOXbiYXC-GL1 | 1 | Mouse #4 | |
| MOX | Liver (in vivo) | 20250707-87MOXbiYXC-GL2 | 1 | Mouse #5 | |
| MOX | Liver (in vivo) | 20250707-87MOXbiYXC-GL3 | 1 | Mouse #6 | |
| MOY | Liver (in vivo) | 20250716-87MOYbiYXC-DE1 | 1 | Mouse #7 | |
| MOY | Liver (in vivo) | 20250716-87MOYbiYXC-DE2 | 1 | Mouse #8 | |
| MEG | Spleen (in vivo) | 20241030-87MEGikEYC-GL1 | 1 | Mouse #1 | |
| MEG | Spleen (in vivo) | 20241030-87MEGikEYC-GL2 | 1 | Mouse #1 | |
| MEG | Spleen (in vivo) | 20241030-87MEGikEYC-GL3 | 1 | Mouse #2 | |
| MEG | Spleen (in vivo) | 20241030-87MEGikEYC-GL4 | 1 | Mouse #2 | |
| MEG | Spleen (in vivo) | 20241030-87MEGikEYC-GL5 | 1 | Mouse #3 | |
| MEG | Spleen (in vivo) | 20241030-87MEGikEYC-GL6 | 1 | Mouse #3 | |
| MOX | Spleen (in vivo) | 20250707-87MOXikYXC-GL1 | 1 | Mouse #4 | |
| MOX | Spleen (in vivo) | 20250707-87MOXikYXC-GL2 | 1 | Mouse #5 | |
| MOX | Spleen (in vivo) | 20250707-87MOXikYXC-GL3 | 1 | Mouse #6 | |
| MOY | Spleen (in vivo) | 20250716-87MOYikYXC-DE1 | 1 | Mouse #7 | |
| MOY | Spleen (in vivo) | 20250716-87MOYikYXC-DE2 | 1 | Mouse #8 | |
| PW | Spleen (in vivo) | 20250926-87PWikYX-DE1 | 1 | Mouse #9 | |
| PW | Spleen (in vivo) | 20250926-87PWikYX-DE2 | 1 | Mouse #10 | |
| PW | Spleen (in vivo) | 20250926-87PWikYX-DE3 | 1 | Mouse #11 | |
| MEG | S-T (in vivo) | 20241030-87MEGmaaFWA-GL1 | 1 | Mouse #1 | |
| MEG | S-T (in vivo) | 20241030-87MEGmaaFWA-GL2 | 1 | Mouse #1 | |
| MEG | S-T (in vivo) | 20241030-87MEGmaaFWA-GL3 | 1 | Mouse #2 | |
| MEG | S-T (in vivo) | 20241030-87MEGmaaFWA-GL4 | 1 | Mouse #2 | |
| MEG | S-T (in vivo) | 20241030-87MEGmaaFWA-GL5 | 1 | Mouse #3 | |
| MEG | S-T (in vivo) | 20241030-87MEGmaaFWA-GL6 | 1 | Mouse #3 | |
| MOX | S-T (in vivo) | 20250707-87MOXmaaFWA-GL1 | 1 | Mouse #4 | |
| MOX | S-T (in vivo) | 20250707-87MOXmaaFWA-GL2 | 1 | Mouse #5 | |
| MOX | S-T (in vivo) | 20250707-87MOXmaaFWA-GL3 | 1 | Mouse #6 | |
| MOY | S-T (in vivo) | 20250716-87MOYmaaFWB-DE1 | 1 | Mouse #7 | |
| MOY | S-T (in vivo) | 20250716-87MOYmaaFWB-DE2 | 1 | Mouse #8 | |
| PW | S-T (in vivo) | 20250926-87PWmaaFWB-DE1 | 1 | Mouse #9 | |
| PW | S-T (in vivo) | 20250926-87PWmaaFWB-DE2 | 1 | Mouse #9 | |
| PW | S-T (in vivo) | 20250926-87PWmaaFWB-DE3 | 1 | Mouse #10 | |
| PW | S-T (in vivo) | 20250926-87PWmaaFWB-DE4 | 1 | Mouse #10 | |
| PW | S-T (in vivo) | 20250926-87PWmaaFWB-DE5 | 1 | Mouse #11 | |
| PW | S-T (in vivo) | 20250926-87PWmaaFWB-DE6 | 1 | Mouse #11 | |
| MEG | S-B (in vivo) | 20241030-87MEGmbaFWA-GL1 | 1 | Mouse #1 | |
| MEG | S-B (in vivo) | 20241030-87MEGmbaFWA-GL2 | 1 | Mouse #1 | |

| | | | | | |
|---|---|---|---|---|---|
| MEG | S-B (in vivo) | 20241030-87MEGmbaFWA-GL3 | 1 | Mouse #2 | |
| MEG | S-B (in vivo) | 20241030-87MEGmbaFWA-GL4 | 1 | Mouse #2 | |
| MEG | S-B (in vivo) | 20241030-87MEGmbaFWA-GL5 | 1 | Mouse #3 | |
| MEG | S-B (in vivo) | 20241030-87MEGmbaFWA-GL6 | 1 | Mouse #3 | |
| MOX | S-B (in vivo) | 20250707-87MOXmbaFWA-GL1 | 1 | Mouse #4 | |
| MOX | S-B (in vivo) | 20250707-87MOXmbaFWA-GL2 | 1 | Mouse #5 | |
| MOX | S-B (in vivo) | 20250707-87MOXmbaFWA-GL3 | 1 | Mouse #6 | |
| MOY | S-B (in vivo) | 20250716-87MOYmbaFWB-DE1 | 1 | Mouse #7 | |
| MOY | S-B (in vivo) | 20250716-87MOYmbaFWB-DE2 | 1 | Mouse #8 | |
| PW | S-B (in vivo) | 20250926-87PWmbaFWB-DE1 | 1 | Mouse #9 | |
| PW | S-B (in vivo) | 20250926-87PWmbaFWB-DE2 | 1 | Mouse #9 | |
| PW | S-B (in vivo) | 20250926-87PWmbaFWB-DE3 | 1 | Mouse #10 | |
| PW | S-B (in vivo) | 20250926-87PWmbaFWB-DE4 | 1 | Mouse #10 | |
| PW | S-B (in vivo) | 20250926-87PWmbaFWB-DE5 | 1 | Mouse #11 | |
| PW | S-B (in vivo) | 20250926-87PWmbaFWB-DE6 | 1 | Mouse #11 | |
| MOX | Plasma (in vivo) | 20250707-87MOXcoBLD-GL1 | 1 | Mouse #4 | |
| MOX | Plasma (in vivo) | 20250707-87MOXcoBLD-GL2 | 1 | Mouse #5 | |
| MOX | Plasma (in vivo) | 20250707-87MOXcoBLD-GL3 | 1 | Mouse #6 | |
| MOY | Plasma (in vivo) | 20250716-87MOYcoBLD-DE1 | 1 | Mouse #7 | |
| MOY | Plasma (in vivo) | 20250716-87MOYcoBLD-DE2 | 1 | Mouse #7 | |
| MOY | Plasma (in vivo) | 20250716-87MOYcoBLD-DE3 | 1 | Mouse #8 | |
| MOY | Plasma (in vivo) | 20250716-87MOYcoBLD-DE4 | 1 | Mouse #8 | |
| MOX | RBC (in vivo) | 20250707-87MOXcwYXC-GL1 | 1 | Mouse #4 | |
| MOX | RBC (in vivo) | 20250707-87MOXcwYXC-GL2 | 1 | Mouse #5 | |
| MOX | RBC (in vivo) | 20250707-87MOXcwYXC-GL3 | 1 | Mouse #6 | |
| MOY | RBC (in vivo) | 20250716-87MOYcwYXC-DE1 | 1 | Mouse #7 | |
| MOY | RBC (in vivo) | 20250716-87MOYcwYXC-DE2 | 1 | Mouse #7 | |
| MOY | RBC (in vivo) | 20250716-87MOYcwYXC-DE3 | 1 | Mouse #8 | |
| MOY | RBC (in vivo) | 20250716-87MOYcwYXC-DE4 | 1 | Mouse #8 | |
| MEG | No target | 20241030-87MEGooBO-GL | 3 | Naïve LiGA | |
| MOX | No target | 20250707-87MOXooBO-GL | 3 | Naïve LiGA | |
| MOY | No target | 20250718-87MOYooBO-GL | 3 | Naïve LiGA | |
| PW | No target | 20250926-87PWooBO-DE | 3 | Naïve LiGA | |
| MOX | hDC-SIGN⁺ Raji | 20250610-87MOXrjdPANC-IF | 5 | Cultured cell line | **Extended Data Figure 3** |
| MOX | hLangerin⁺ Raji | 20250610-87MOXrjlPANC-IF | 5 | Cultured cell line | |
| MOX | WT Raji cells | 20250610-87MOXehPANC-IF | 5 | Cultured cell line | |
| MOX | No target | 20250610-87MOXooNA-EG | 5 | Naïve LiGA | |

**Table S2:** List of LiGA data used in this work.
All deep sequencing datasets mentioned in this study are publicly available at https://48hd.cloud/. * indicates the tested protein was biotinylated. F, female; M, male; Δ, cells were treated with heat-deactivated sialidase; S, cells were treated with sialidase (from *Vibrio Cholerae*); MO, monocytes; B, B lymphocytes; NK, natural killer cells; S-T, splenic T lymphocytes; S-B, splenic B lymphocytes; RBC, red blood cells.

| Glycan | Display density | MEG | MOX | MOY | PW |
|---|---|---|---|---|---|
| α-L-Man | [20] | Y | Y | Y | Y |
| α-L-Man | [50] | Y | Y | Y | Y |
| α-L-Man | [150] | Y | Y | Y | Y |
| α-L-Man | [500] | Y | Y | Y | Y |
| α-L-Man | [4000] | N | Y | Y | Y |
| α-D-Man | [20] | Y | Y | Y | Y |
| α-D-Man | [50] | Y | Y | Y | Y |
| α-D-Man | [150] | Y | Y | Y | Y |
| α-D-Man | [500] | Y | Y | Y | Y |
| α-D-Man | [4000] | N | Y | Y | Y |
| α-L-Glc | [20] | N | Y | Y | Y |
| α-L-Glc | [50] | N | Y | Y | Y |
| α-L-Glc | [150] | N | Y | Y | Y |
| α-L-Glc | [500] | N | Y | Y | Y |
| α-L-Glc | [4000] | N | N | N | N |
| β-L-Glc | [20] | N | Y | Y | Y |
| β-L-Glc | [50] | N | Y | Y | Y |
| β-L-Glc | [150] | N | Y | Y | Y |
| β-L-Glc | [500] | N | Y | Y | Y |
| β-L-Glc | [4000] | N | N | N | N |
| β-D-Glc | [20] | N | Y | Y | Y |
| β-D-Glc | [50] | N | Y | Y | Y |
| β-D-Glc | [150] | N | Y | Y | Y |
| β-D-Glc | [500] | N | Y | Y | Y |
| β-D-Glc | [4000] | N | N | Y | Y |
| α-L-Gal | [20] | N | Y | Y | Y |
| α-L-Gal | [50] | N | Y | Y | Y |
| α-L-Gal | [150] | N | Y | Y | Y |
| α-L-Gal | [500] | N | Y | Y | Y |
| α-L-Gal | [4000] | N | Y | Y | Y |
| β-L-Gal | [20] | N | Y | Y | Y |
| β-L-Gal | [50] | N | Y | Y | Y |
| β-L-Gal | [150] | N | Y | Y | Y |
| β-L-Gal | [500] | N | Y | Y | Y |
| β-L-Gal | [4000] | N | Y | Y | Y |
| β-D-Gal | [20] | N | Y | Y | Y |
| β-D-Gal | [50] | N | Y | Y | Y |
| β-D-Gal | [150] | N | Y | Y | Y |
| β-D-Gal | [500] | N | Y | Y | Y |
| β-D-Gal | [4000] | N | Y | Y | Y |

**Table S3:** LiGA used in in vivo glycan binding profiling.
LiGA used for **in vivo organ homing analysis** of glycosylated phages in four independent campaigns: MEG (20241030; n = 3 mice, with each mouse sequenced twice), MOX (20250707; n = 3 mice), MOY (20250716; n = 2 mice), and PW (20250926; n = 3 mice). MEG, MOX, MOY, and PW are different LiGAs. Y, the glycosylated phage exists in the tested LiGA; N, the glycosylated phage absents in the tested LiGA.

| | Description* |
|---|---|
| **1. Sample: Glycan-containing sample (e.g. glycan, glycoprotein, cell lysate, cell, glycopeptide, etc.)** | |
| Description of Sample | Glycophage constructs |
| Sample preparation protocol | Glycophage construct samples were labeled with TAMRA prior to analysis on the lectin microarrays, and thus required no sample preparation. |
| Labeling protocol for sample detection | Samples are labelled with TAMRA. |
| Two-color reference (if used) | N/A |
| Assay protocol | Lectin microarrays are blocked with blocking buffer (50 mM ethanolamine and 100 mM boric acid) for one hour at room temperature. Slides are rinsed once with PBST (0.01%) and once with 1X PBS for 5 minutes each, then dried using a slide spinner. Each slide is mounted on a 24-well format hybridization cassette (Arrayit), in which each well contains a subarray. To each well, 50 uL of sample was added, then diluted with 50 uL PBST (with 0.1 mM $Ca^{2+}$ to reach the final volume (100uL) and PBST concentration (0.005%). Slides are incubated in a hybridization oven on a rocker tray for one hour at 30°C in the dark. After hybridization, arrays are washed with PBST (0.01%) once for one minute, then once for five minutes. Arrays are lastly washed once with PBS for one minute, then once for ten minutes. Once finished, slides are removed from the cassette and briefly immersed in ultrapure water, then dried using a slide spinner. |
| **2. Lectin Library** | |
| General description of the lectin library used in the array | Lectin microarrays are generated in house. |
| List of lectins and/or glycan-binding proteins, their source, concentration, and buffer | Please see **Table S5**. |
| Modification of lectins | N/A |
| **3. Immobilization Surface; e.g., Microarray Slide** | |
| Immobilization surface | Nexterion Slide H Barcoded 3D Hydrogel Coated. |
| Manufacturer | Schott North America. |
| Custom preparation of the surface | N/A |
| **4. Array Production** | |
| Description of Arrayer | Nano-Plotter 2.1 piezoelectric printer (GeSim, Germany) with cooled microwell plate holder and cooled printing deck. |
| Lectin deposition | Triplicates of each lectin are printed onto each subarray. |

| | |
|---|---|
| Printing conditions | Dilute lectins to the pre-determined concentrations in the print buffer (final concentration of print buffer: 1 mM monosaccharide in PBS, 5 ng/mL Atto 532; Please see Supplemental Table 1 for the concentrations of lectins). Load the mixed solution to the microplate. Before printing, check the humidity of the print chamber. The humidity should be kept around 50% throughout the entire print. Ensure both microwell plate holder and printing deck are cooled. Adjust the cooling temperature based on ambient temperature and the temperature of the cooled slide deck surface, preventing moisture building up inside the print chamber. Once printing is complete, allow the slides to dry for at least one hour before vacuum sealing and storage at -20℃. |
| Array layout | Each microarray contains 21 subarrays (3 columns and 7 rows). In each subarray, triplicates of a lectin are printed, with six lectins in each row. The number of columns is 18, and the row number depends on how many lectin probes are printed on the arrays (i.e. 132 lectins require 22 rows). |
| Quality control | The well-characterized glycoproteins glycophorin, ovalbumin, asialofetuin, 1:1 A549:HEK293T cell lysate, human serum, fetuin, and RNase B are used for quality assurance of the printed microarrays. |
| **5. Detector and Data Processing** | |
| Instrument (scanner, flow cytometer) | Fluorescent Slide Scanner GenePix 4400A (Molecular Devices). |
| Instrument settings | A low resolution preview scan of the slide is performed to adjust photomultiplier tube (PMT) gain for the channel used (TAMRA: 532nm) so that the signals are not saturated and within the linear detection range. After adjusting PMT gain, slides are scanned in the given channel at 5 µm resolution. |
| Image analysis software | GenePix Pro 7 (Molecular Devices). |
| Data processing and statistical analysis | Extracted data is processed for quality checks using Grubbs outlier test with $\alpha = 0.05$. Log2 values of the average signals are median-normalized over the individual subarray in each channel. |
| **6. Lectin Microarray Data Presentation** | |
| Data presentation and interpretation | Raw values output by Genepix Pro 7 software were thresholded by a cut-off of >2000 AU. |
| **7. Data Location** | |
| Data Location | https://doi.org/10.7303/syn76179151 |

**Table S4:** Lectin Microarray Information.

| Lectin | Species/Origin | Print Conc. (μg/mL) | Rough Specificity /Inhibitory monosaccharide | Vendor/Source |
|---|---|---|---|---|
| AAA | *Anguilla anguilla agglutinin* | 2000 | Fucose | Vector |
| AAL | *Aleuria aurantia* | 2000 | Fucose | Medicago/Vector |
| ACA | *Amaranthus Caudatus* | 2000 | Gal-β1,3-GalNAc | EY |
| AIA | *Artocarpus integrifolia* | 2000 | β1,3-GalNAc | EY/Glycomatrix/Vector |
| AMA | *Allium moly* | 2000 | Oligo mannose | Vector |
| Anti-B.G. Lewis A | MAb mouse IgG1 [7LE] | undiluted | Lewis A | Abcam |
| Anti-B.G. Lewis B | MAb mouse IgM [2-25LE] | undiluted | Lewis B | Abcam/Sigma |
| Anti-B.G. Lewis X | MAb mouse IgM [P12] | undiluted | Lewis X | Sigma |
| Anti-B.G. Lewis Y | MAb mouse IgM [F3] | undiluted | Lewis Y | Abcam |
| Anti-Blood Group A | MAb mouse IgM [HE-193] | undiluted | Blood Group A | Thermo Fisher |
| Anti-Blood Group B | MAb mouse IgM [HEB-29] | undiluted | Blood Group B | Abcam/ Thermo Fisher |
| Anti-Blood Group H | MAb mouse IgG3 [17-206] | undiluted | Blood Group H | Thermo Fisher |
| Anti-Polysialic Acid | MAb mouse IgG2a [735] | undiluted | Polysialic Acid | Absolute Antibody |
| Anti-IgA | MAb IgG1 [KT41] | undiluted | IgA | Abcam |
| Anti-IgG | MAb IgG2a [KT131] | undiluted | IgG | Abcam |
| Anti-IgM | MAb IgG1[KT16] | undiluted | IgM | Abcam |
| AOL | *Aspergillus oryzae* | 2000 | Fucose | TCI America |
| ASA | *Allium sativum* | 2000 | Mannose | EY |
| BanLec H84T | *Musa acuminate* | 2000 | High Mannose | Generated in house |
| BC2L-A | *Burkholderia cenocepacia* | 2000 | Mannose | Elicityl |
| BPA/BPL | *Bauhinia purpurea* | 2000 | β-Gal / β-GalNAc | Vector |
| CA | *Colchicum autumnale* | 2000 | Bi-antennary N-linked glycans | EY |
| ConA | *Canavalia ensiformis* | 2000 | Tri-mannose core | Thermo Fisher/Vector |
| DBA | *Dolichos biflorus* | 2000 | GalNAc | Vector |
| diCBM40 | engineered NanI from *Clostridium perfringens* | 1500 | α Sialylation | Generated in house |
| DSA | *Datura stramonium* | 2000 | LacNAc | Vector |
| ECA | *Erythrina cristagalli* | 2000 | LacNAc | EY/Vector |
| GafD | *Escherichia coli* | 2000 | GlcNAc | Generated in house |
| GNA/GNL | *Galanthus nivalis* | 2000 | Oligo mannose | Sigma/Vector |
| GS/GSL-I | *Griffonia simplicifoia-I* | 2000 | α-Gal / Lac | Vector |
| GS/GSL-II | *Griffonia simplicifoia-II* | 2000 | GlcNAc | Vector |
| HHL | *Hippeastrum hybrid* | 2000 | Oligo/High mannose | BioWorld |
| HPA | *Helix pomatia* | 2000 | Blood Group A | Sigma |
| LcH | *Lens culinaris* | 2000 | Core Fucose | EY/Medicago |
| LEA/LEL | *Lycopersicon esculentum* | 2000 | GlcNAc | Vector |
| LTL Lotus | *Lotus tetragonolobus* | 2000 | Fucose | EY/Vector |
| MAA/MAL-I | *Maackia amurensis-I* | 2000 | Sialylation/Sulfation | EY/Vector |
| MAA/MAL-II | *Maackia amurensis-II* | 2000 | Sialylation/Sulfation | Vector |
| MNA-G | *Morus nigra Morniga G* | 2000 | GalNAc | EY |
| MNA-M | *Morus nigra Morniga M* | 2000 | Oligo mannose / Gal | EY |
| MPA/MPL | *Maclura pomifera* | 2000 | β1,3-GalNAc | Vector |

s

| | | | | |
|---|---|---|---|---|
| NPA/NPL | *Narcissus pseudonarcissus* | 2000 | Oligo mannose | EY |
| PA-IL | *Pseudomonas aeruginosa* | 2000 | Gal | Generated in house |
| PA-IIL | *Pseudomonas aeruginosa* | 2000 | Fucose | Elicityl |
| PapGII | *Escherichia coli* | 2000 | Gal | Generated in house |
| PHA-E | *Phaseolus vulgaris Erythroagglutinin* | 2000 | Bisecting GlcNAc | Sigma/Vector |
| PHA-L | *Phaseolus vulgaris Leukoagglutinin* | 2000 | β1,6 Branching N-Link glycans | Medicago/Vector |
| PhoSL | *Pholiota squarrosa* | 2000 | Core Fucose | Bio Basic |
| PNA | *Arachis hyogaea* | 2000 | Gal-β1,3-GalNAc | EY/Vector |
| Recombinant Protein A | *Staphylococcus aureus* | 2000 | Immunoglobulins | Thermo Fisher |
| Recombinant Protein G | *Streptococcus* Group G | 2000 | Immunoglobulins | Sigma |
| Recombinant Protein L | *Peptostreptococcus magnus* | 2000 | Immunoglobulins | Thermo Fisher |
| PSA | *Pisum sativum* | 2000 | Core Fucose | Vector |
| PSL | *Polyporus squamosus* | 2000 | α2,6 sialylation | TCI America |
| RCA120 | *Ricinus Communis Agglutinin I* | 2000 | Gal / Lac | Vector |
| rGRFT | *Griffithsia* | 1700 | High mannose | Generated in house |
| Ricin B Chain | *Ricinus communis* | 2000 | Gal | Vector |
| RCA120 | *Ricinus communis* | 2000 | Gal | Vector |
| RPA | *Robinia pseudoacacia* | 2000 | Multiantennary N-glycans with bisecting GlcNAc | EY |
| SBA | *Glycine max* | 2000 | LacdiNAc | Vector |
| SLBR-B | *Streptococcus gordonii M99* | 2300 | α2,6 sialylation | Generated in house |
| SLBR-H | *Streptococcus gordonii DL1* | 2000 | α2,3 sialylation | Generated in house |
| SLBR-N | *Streptococcus gordonii UB10712* | 2000 | α2,3 sialylation | Generated in house |
| SNA | *Sambucus nigra* | 2000 | α2,6 sialylation | EY/Vector |
| SNA-II | *Sambucus nigra-II* | 2000 | α2 Fucose /oligo mannose | EY |
| STL | *Solanum tuberosum Agglutinin* | 2000 | LacNAc | Vector |
| TJA-II | *Trichosanthes japonica-II* | 2000 | α2 Fucose | Aniara Diagnostica/Medicago |
| TL | *Tulipa sp.* | 2000 | GlcNAc | BioWorld |
| UDA | *Urtica dioica* | 2000 | GlcNAc / Oligo mannose | EY |
| UEA-I | *Ulex europaaeus-I* | 2000 | α2 Fucose | BioWorld/GeneTex/Vector |
| UEA-II | *Ulex europaaeus-II* | 2000 | GlcNAc | EY/BioWorld |
| VVA | *Vicia villosa* | 2000 | Terminal GalNAc | Vector |
| WFA | *Wisteria floribunda* | 2000 | GalNAc-β1,4 | Vector |
| WGA | *Triticum vulgare* | 2000 | GlcNAc | GeneTex |

**Table S5:** Lectins used in microarrays.

| Glycan | Top | Bottom | HillSlope | N_points | N_concentration | Fit_status | $R^2$ | IC50 (mM) |
|---|---|---|---|---|---|---|---|---|
| Man3-[1300] | 2615.147 | 0.000 | 1.688 | 14 | 6 | OK | 0.930 | IC50 = 20.1 mM |
| Man3-[1730] | 393.491 | 3.052 | 4.600 | 15 | 6 | OK | 0.887 | IC50 = 31.3 mM |
| Man4-[20] | 146.134 | 0.000 | 1.349 | 35 | 6 | OK | 0.461 | IC50 = 18.7 mM |
| Man4-[100] | 21.315 | 656.497 | 10.000 | 22 | 6 | OK | 0.414 | IC50 = 150 mM |
| Man4-[300] | 65.184 | 4.206 | 3.547 | 80 | 6 | OK | 0.801 | IC50 = 8.49 mM |
| Man4-[720] | 765.197 | 36.319 | 2.144 | 54 | 6 | OK | 0.520 | IC50 = 13.3 mM |
| Man4-[740] | 249.317 | 0.000 | 1.562 | 30 | 6 | OK | 0.951 | IC50 = 14.1 mM |
| L-Man2-[500] | 13.358 | 6.037 | 3.975 | 26 | 6 | OK | 0.305 | IC50 = 2.9 mM |
| LeA-[950] | 714.135 | 0.000 | 2.048 | 14 | 6 | OK | 0.870 | IC50 = 35.8 mM |
| Lex-[810] | 213.114 | 13.006 | 10.000 | 16 | 6 | OK | 0.960 | IC50 = 25.2 mM |
| Gala3Lex-[620] | 830.518 | 0.000 | 0.704 | 14 | 6 | OK | 0.928 | IC50 = 2.81 mM |
| L-Glca-[150] | 13.918 | 42.051 | 8.928 | 33 | 6 | OK | 0.255 | IC50 = 13.5 mM |
| L-Glca-[500] | 10.519 | 0.000 | 0.066 | 17 | 6 | OK | 0.478 | IC50 = 0.01 mM |
| L-Glca-[4000] | 447.668 | 0.000 | 0.433 | 68 | 6 | OK | 0.818 | IC50 = 0.119 mM |
| L-Glcb-[20] | NA | NA | NA | 68 | 6 | Fit failed | NA | IC50: NA |
| L-Glcb-[50] | 18.222 | 10.699 | 0.010 | 13 | 6 | OK | 0.119 | IC50 = 297 mM |
| L-Glcb-[150] | 19.694 | 22.534 | 10.000 | 35 | 6 | OK | 0.031 | IC50 = 42.6 mM |
| L-Glcb-[500] | NA | NA | NA | 37 | 6 | Fit failed | NA | IC50: NA |
| L-Glcb-[4000] | 10.918 | 7.031 | 0.010 | 12 | 6 | OK | 0.062 | IC50 = 0.0167 mM |
| D-Glcb-[20] | 23.955 | 40.680 | 0.010 | 72 | 6 | OK | 0.061 | IC50 = 80.1 mM |
| D-Glcb-[50] | 15.824 | 44.680 | 0.010 | 44 | 6 | OK | 0.169 | IC50 = 227 mM |
| D-Glcb-[150] | 25.750 | 44.194 | 0.010 | 56 | 6 | OK | 0.133 | IC50 = 83.2 mM |
| D-Glcb-[500] | 23.208 | 285.963 | 10.000 | 48 | 6 | OK | 0.277 | IC50 = 130 mM |
| D-Glcb-[4000] | 975.701 | 9.922 | 9.552 | 100 | 6 | OK | 0.465 | IC50 = 6.81 mM |
| L-Gala-[500] | 21.615 | 0.527 | 9.998 | 68 | 6 | OK | 0.922 | IC50 = 6.16 mM |
| L-Gala-[4000] | 827.225 | 0.000 | 1.605 | 89 | 6 | OK | 0.362 | IC50 = 43.9 mM |
| L-Galb-[150] | 13.743 | 10.447 | 10.000 | 6 | 6 | OK | 0.260 | IC50 = 39.2 mM |
| L-Galb-[500] | 32.089 | 26.113 | 2.759 | 61 | 6 | OK | 0.049 | IC50 = 3.65 mM |
| L-Galb-[4000] | 425.253 | 19.041 | 5.261 | 81 | 6 | OK | 0.504 | IC50 = 23.7 mM |
| D-Galb-[20] | 66.209 | 80.265 | 0.010 | 47 | 6 | OK | 0.009 | IC50 = 0.0478 mM |
| D-Galb-[50] | 42.471 | 31.009 | 10.000 | 52 | 6 | OK | 0.154 | IC50 = 14.4 mM |
| D-Galb-[150] | 25.808 | 343.174 | 10.000 | 71 | 6 | OK | 0.052 | IC50 = 148 mM |
| D-Galb-[500] | 52.584 | 31.886 | 10.000 | 54 | 6 | OK | 0.157 | IC50 = 9.75 mM |
| D-Galb-[4000] | 21.449 | 567.713 | 10.000 | 38 | 6 | OK | 0.210 | IC50 = 150 mM |
| L-Mana-[20] | 14.417 | 19.141 | 0.010 | 6 | 6 | OK | 0.121 | IC50 = 94.4 mM |
| L-Mana-[150] | 14.378 | 4.206 | 1.257 | 6 | 6 | OK | 0.983 | IC50 = 0.971 mM |
| L-Mana-[500] | NA | NA | NA | 23 | 6 | Fit failed | NA | IC50: NA |
| L-Mana-[4000] | 57.175 | 62.808 | 0.010 | 43 | 6 | OK | 0.002 | IC50 = 0.0116 mM |
| D-Mana-[4000] | 27.331 | 0.114 | 10.000 | 84 | 6 | OK | 0.774 | IC50 = 21.3 mM |
| L-Fuca-[20] | 32.860 | 39.690 | 10.000 | 14 | 6 | OK | 0.333 | IC50 = 11 mM |
| L-Fuca-[150] | 15.214 | 1.628 | 2.343 | 12 | 6 | OK | 0.855 | IC50 = 2.74 mM |
| L-Fuca-[500] | 1845.453 | 0.000 | 0.844 | 14 | 6 | OK | 0.869 | IC50 = 5.8 mM |
| L-Fuca-[4000] | 17.371 | 38.330 | 1.800 | 51 | 6 | OK | 0.156 | IC50 = 1.46 mM |
| L-Fucb-[20] | 21.257 | 16299.465 | 10.000 | 16 | 6 | OK | 0.497 | IC50 = 199 mM |
| L-Fucb-[50] | 35.034 | 27.901 | 2.921 | 12 | 6 | OK | 0.072 | IC50 = 3.85 mM |
| L-Fucb-[150] | 23.233 | 1323.536 | 10.000 | 12 | 6 | OK | 0.812 | IC50 = 148 mM |
| L-Fucb-[500] | 1696.232 | 46.143 | 10.000 | 14 | 6 | OK | 0.993 | IC50 = 3.07 mM |
| L-Fucb-[4000] | 105.472 | 3.584 | 6.666 | 43 | 6 | OK | 0.642 | IC50 = 35.5 mM |
| D-Fuca-[20] | 16.263 | 36.200 | 10.000 | 12 | 6 | OK | 0.705 | IC50 = 11.3 mM |
| D-Fuca-[50] | NA | NA | NA | 14 | 6 | Fit failed | NA | IC50: NA |
| D-Fuca-[150] | 48.703 | 25.575 | 3.218 | 6 | 6 | OK | 0.258 | IC50 = 3.61 mM |
| D-Fuca-[500] | 87.739 | 34.770 | 5.808 | 12 | 6 | OK | 0.415 | IC50 = 3.57 mM |
| D-Fuca-[4000] | 13.790 | 0.036 | 0.010 | 6 | 6 | OK | 0.278 | IC50 = 0.013 mM |
| D-Fucb-[20] | 62.438 | 21.547 | 8.316 | 12 | 6 | OK | 0.415 | IC50 = 3.72 mM |
| D-Fucb-[500] | 27.191 | 21.916 | 0.010 | 6 | 6 | OK | 0.168 | IC50 = 0.0278 mM |
| D-Fucb-[4000] | 22.935 | 19.580 | 0.010 | 14 | 6 | OK | 0.175 | IC50 = 118 mM |
| AzOH-[100] | 48.882 | 13.532 | 0.010 | 20 | 6 | OK | 0.180 | IC50 = 0.0106 mM |
| AzOH-[500] | 16.092 | 6.039 | 0.963 | 39 | 6 | OK | 0.092 | IC50 = 50.9 mM |

**Table S6:** Fitted $IC_{50}$ for inhibition of glycophages binding to DC-SIGN$^+$ Raji by free D-Man.
Mirror-LiGA was incubated with DC-SIGN$^+$ Raji cells in the presence of increasing concentrations of free D-(+)-mannose. Sequencing counts for each glycophage were normalized to those of the internal standard (unmodified blank phage) and fitted to inhibition curves. Fitted IC50 values with good fits ($R^2 > 0.75$) were highlighted in green, whereas those with moderate fits ($0.40 < R^2 \leq 0.75$) were highlighted in yellow. Glycans showing no enrichment in the absence of D-Man (0 mM) were excluded from the analysis. (**Fig. 3** and **Extended Data Fig. 3**).

### 3. NMR spectra

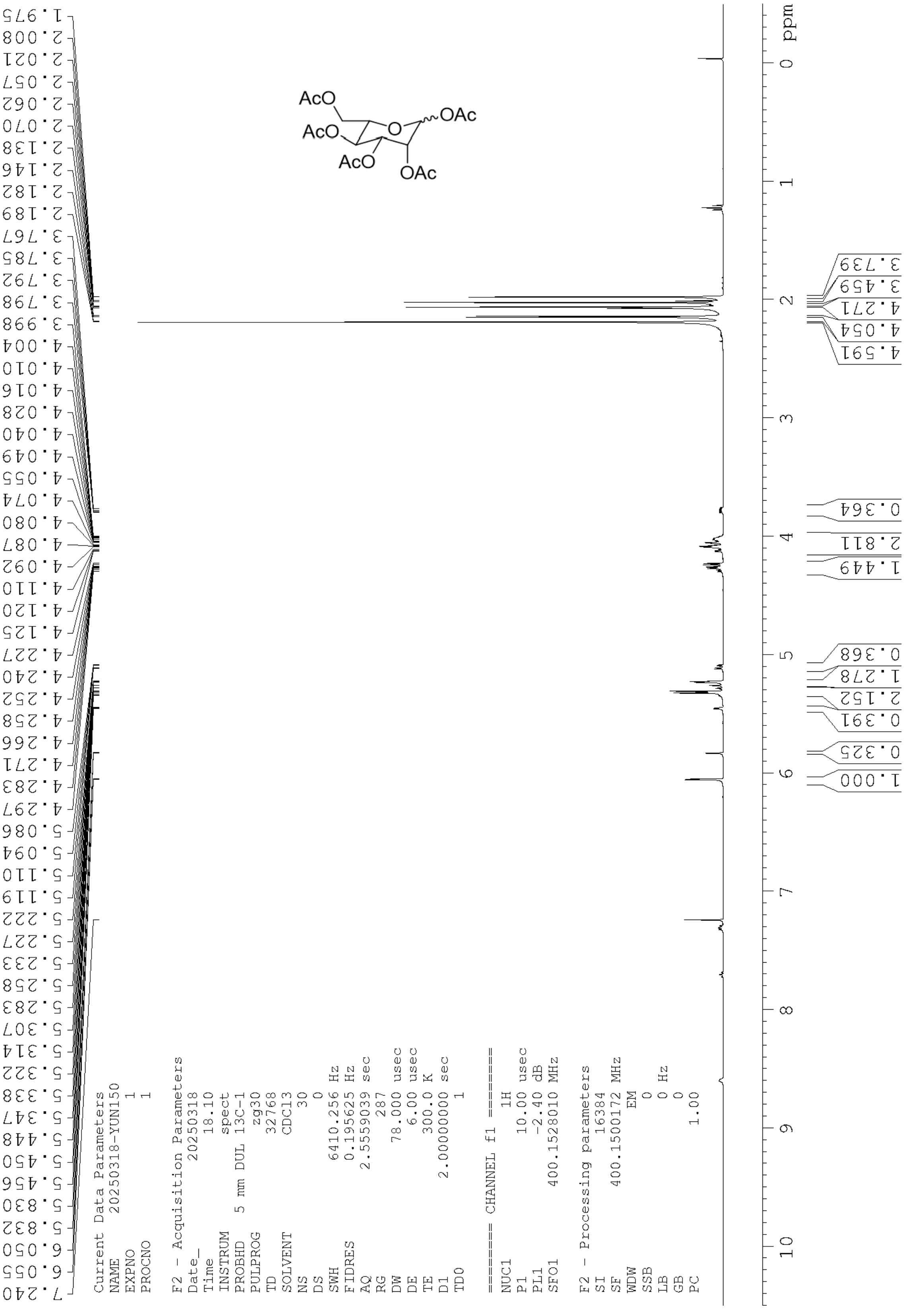

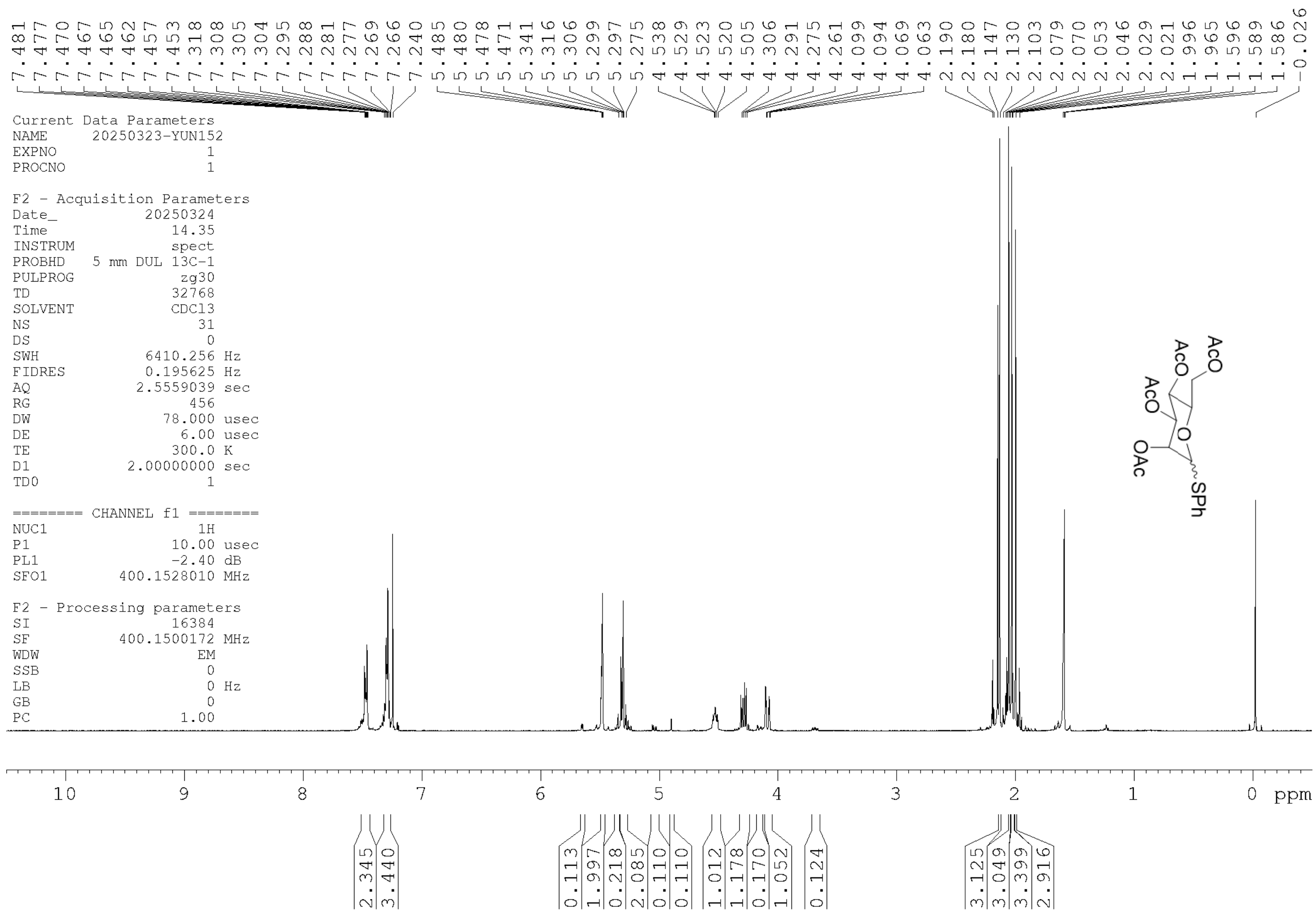
7.481
7.477
7.470
7.467
7.465
7.462
7.457
7.453
7.318
7.308
7.305
7.304
7.295
7.288
7.281
7.277
7.269
7.266
7.240
5.485
5.480
5.478
5.471
5.341
5.316
5.306
5.299
5.297
5.275
4.538
4.529
4.523
4.520
4.505
4.306
4.291
4.275
4.261
4.099
4.094
4.069
4.063
2.190
2.180
2.147
2.130
2.103
2.079
2.070
2.053
2.046
2.029
2.021
1.996
1.965
1.596
1.589
1.586
-0.026
Current Data Parameters
NAME 20250323-YUN152
EXPNO 1
PROCNO 1
F2 - Acquisition Parameters
Date_ 20250324
Time 14.35
INSTRUM spect
PROBHD 5 mm DUL 13C-1
PULPROG zg30
TD 32768
SOLVENT CDCl3
NS 31
DS 0
SWH 6410.256 Hz
FIDRES 0.195625 Hz
AQ 2.5559039 sec
RG 456
DW 78.000 usec
DE 6.00 usec
TE 300.0 K
D1 2.00000000 sec
TD0 1
======== CHANNEL f1 ========
NUC1 1H
P1 10.00 usec
PL1 -2.40 dB
SFO1 400.1528010 MHz
F2 - Processing parameters
SI 16384
SF 400.1500172 MHz
WDW EM
SSB 0
LB 0 Hz
GB 0
PC 1.00
AcO
AcO
AcO
O
OAc
SPh
10
9
8
7
6
5
4
3
2
1
0
ppm
2.345
3.440
0.113
1.997
0.218
2.085
0.110
0.110
1.012
1.178
0.170
1.052
0.124
3.125
3.049
3.399
2.916

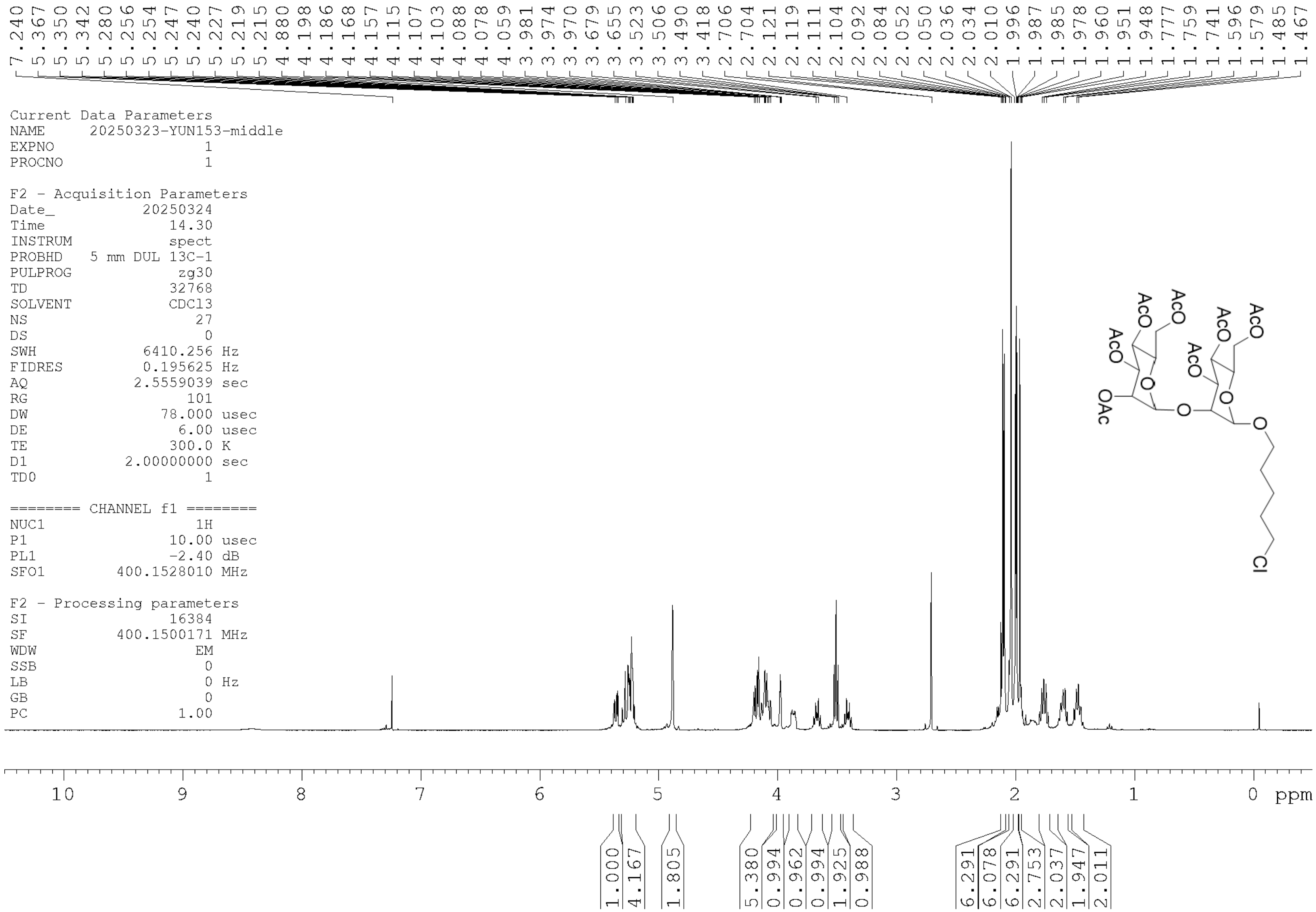
7.240 5.367 5.350 5.342 5.280 5.256 5.254 5.247 5.240 5.227 5.219 5.215 4.880 4.198 4.186 4.168 4.157 4.115 4.107 4.103 4.088 4.078 4.059 3.981 3.974 3.970 3.679 3.655 3.523 3.506 3.490 3.418 2.706 2.704 2.121 2.119 2.111 2.104 2.092 2.084 2.052 2.050 2.036 2.034 2.010 1.996 1.987 1.985 1.978 1.960 1.951 1.948 1.777 1.759 1.741 1.596 1.579 1.485 1.467
Current Data Parameters
NAME 20250323-YUN153-middle
EXPNO 1
PROCNO 1
F2 - Acquisition Parameters
Date_ 20250324
Time 14.30
INSTRUM spect
PROBHD 5 mm DUL 13C-1
PULPROG zg30
TD 32768
SOLVENT CDCl3
NS 27
DS 0
SWH 6410.256 Hz
FIDRES 0.195625 Hz
AQ 2.5559039 sec
RG 101
DW 78.000 usec
DE 6.00 usec
TE 300.0 K
D1 2.00000000 sec
TD0 1
======== CHANNEL f1 ========
NUC1 1H
P1 10.00 usec
PL1 -2.40 dB
SFO1 400.1528010 MHz
F2 - Processing parameters
SI 16384
SF 400.1500171 MHz
WDW EM
SSB 0
LB 0 Hz
GB 0
PC 1.00
AcO
AcO
AcO
AcO
AcO
AcO
OAc
O
O
O
O
Cl
10 9 8 7 6 5 4 3 2 1 0 ppm
1.000 4.167 1.805 5.380 0.994 0.962 0.994 1.925 0.988 6.291 6.078 6.291 2.753 2.037 1.947 2.011

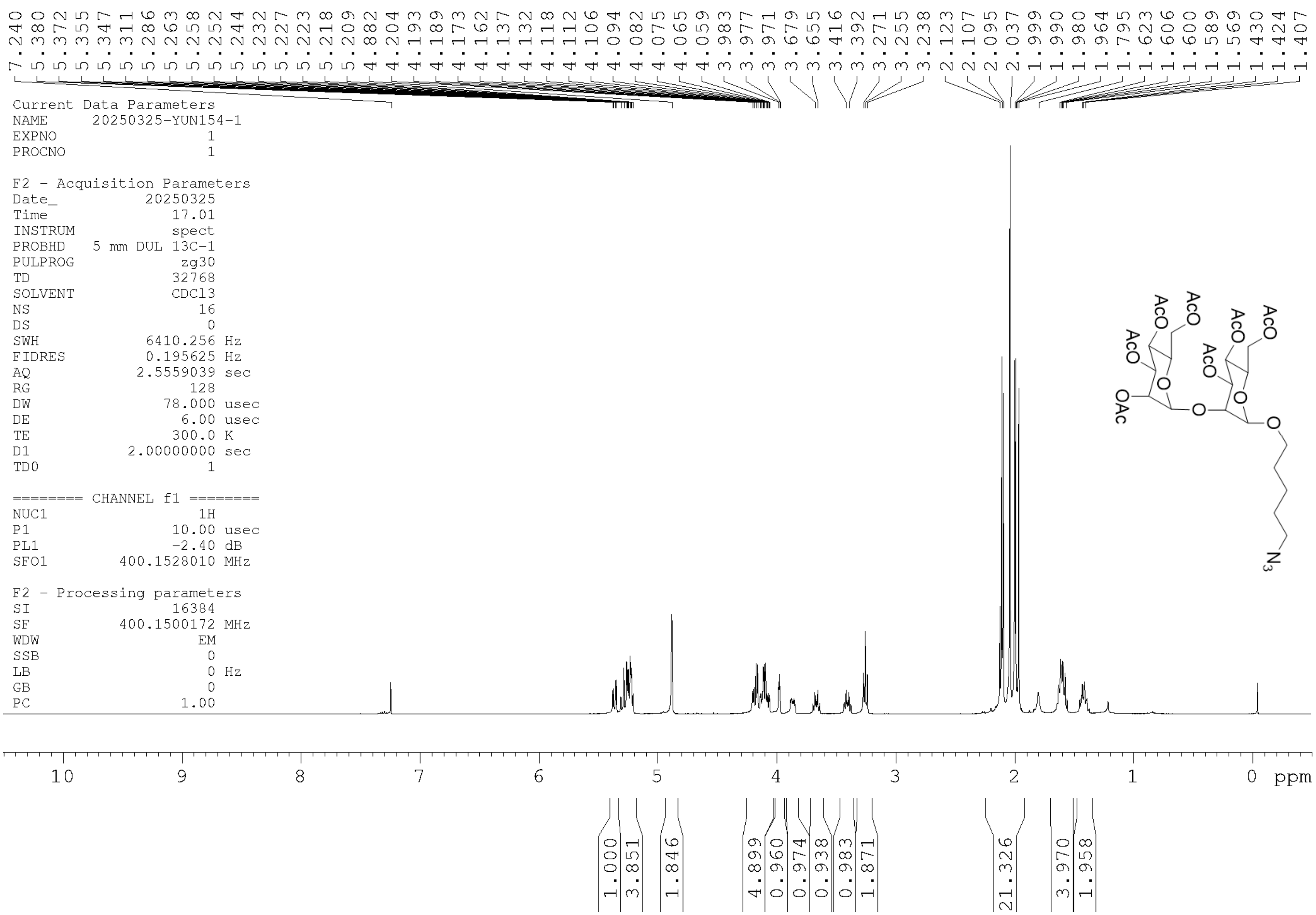

7.240
5.380
5.372
5.355
5.347
5.311
5.286
5.263
5.258
5.252
5.244
5.232
5.227
5.223
5.218
5.209
4.882
4.204
4.193
4.189
4.173
4.162
4.137
4.132
4.118
4.112
4.106
4.094
4.082
4.075
4.065
4.059
3.983
3.977
3.971
3.679
3.655
3.416
3.392
3.271
3.255
3.238
2.123
2.107
2.095
2.037
1.999
1.990
1.980
1.964
1.795
1.623
1.606
1.600
1.589
1.569
1.430
1.424
1.407
Current Data Parameters
NAME 20250325-YUN154-1
EXPNO 1
PROCNO 1
F2 - Acquisition Parameters
Date_ 20250325
Time 17.01
INSTRUM spect
PROBHD 5 mm DUL 13C-1
PULPROG zg30
TD 32768
SOLVENT CDCl3
NS 16
DS 0
SWH 6410.256 Hz
FIDRES 0.195625 Hz
AQ 2.5559039 sec
RG 128
DW 78.000 usec
DE 6.00 usec
TE 300.0 K
D1 2.00000000 sec
TD0 1
======== CHANNEL f1 ========
NUC1 1H
P1 10.00 usec
PL1 -2.40 dB
SFO1 400.1528010 MHz
F2 - Processing parameters
SI 16384
SF 400.1500172 MHz
WDW EM
SSB 0
LB 0 Hz
GB 0
PC 1.00
AcO
AcO
AcO
OAc
AcO
AcO
AcO
O
O
O
O
$N_3$
10
9
8
7
6
5
4
3
2
1
0
ppm
1.000
3.851
1.846
4.899
0.960
0.974
0.938
0.983
1.871
21.326
3.970
1.958

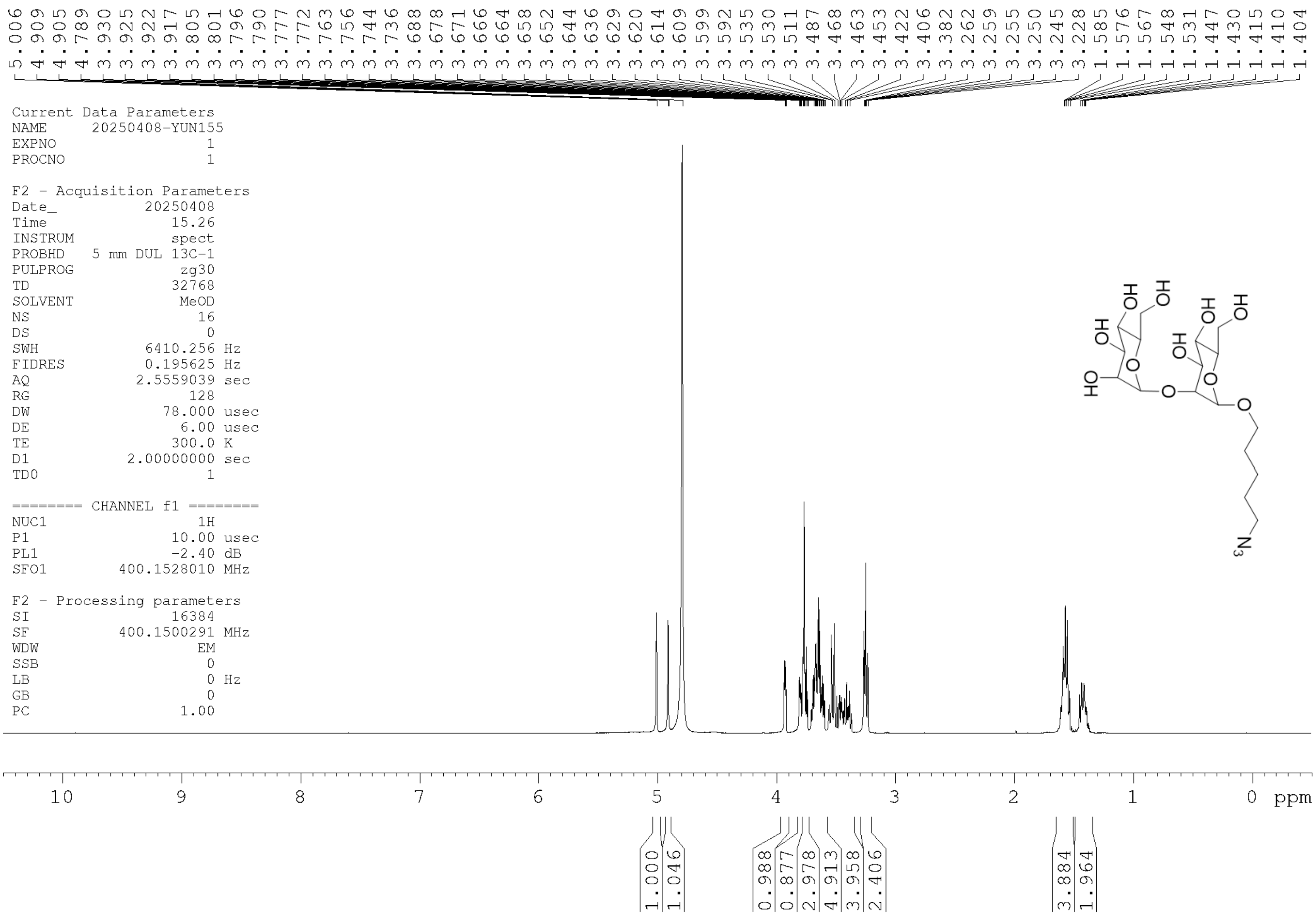

5.006
4.909
4.905
4.789
3.930
3.925
3.922
3.917
3.805
3.801
3.796
3.790
3.777
3.772
3.763
3.756
3.744
3.736
3.688
3.678
3.671
3.666
3.664
3.658
3.652
3.644
3.636
3.629
3.620
3.614
3.609
3.599
3.592
3.535
3.530
3.511
3.487
3.468
3.463
3.453
3.422
3.406
3.382
3.262
3.259
3.255
3.250
3.245
3.228
1.585
1.576
1.567
1.548
1.531
1.447
1.430
1.415
1.410
1.404
Current Data Parameters
NAME 20250408-YUN155
EXPNO 1
PROCNO 1
F2 - Acquisition Parameters
Date_ 20250408
Time 15.26
INSTRUM spect
PROBHD 5 mm DUL 13C-1
PULPROG zg30
TD 32768
SOLVENT MeOD
NS 16
DS 0
SWH 6410.256 Hz
FIDRES 0.195625 Hz
AQ 2.5559039 sec
RG 128
DW 78.000 usec
DE 6.00 usec
TE 300.0 K
D1 2.00000000 sec
TD0 1
======== CHANNEL f1 ========
NUC1 1H
P1 10.00 usec
PL1 -2.40 dB
SFO1 400.1528010 MHz
F2 - Processing parameters
SI 16384
SF 400.1500291 MHz
WDW EM
SSB 0
LB 0 Hz
GB 0
PC 1.00
HO
HO
HO
O
OH
HO
HO
HO
O
O
O
N3
10
9
8
7
6
5
4
3
2
1
0
ppm
1.000
1.046
0.988
0.877
2.978
4.913
3.958
2.406
3.884
1.964

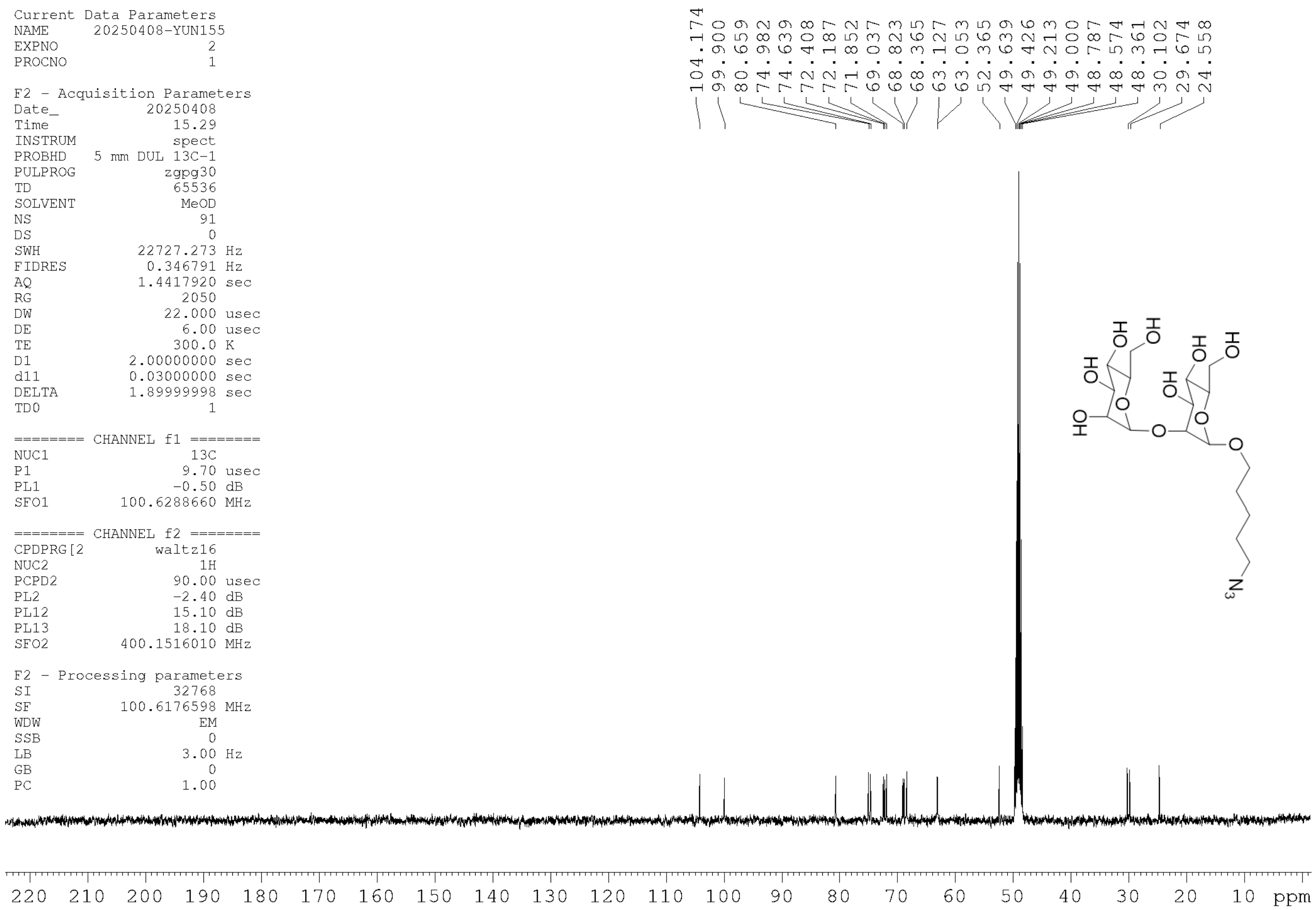

| | |
|---|---|
| Current Data Parameters | |
| NAME | 20250408-YUN155 |
| EXPNO | 2 |
| PROCNO | 1 |
| F2 - Acquisition Parameters | |
| Date_ | 20250408 |
| Time | 15.29 |
| INSTRUM | spect |
| PROBHD | 5 mm DUL 13C-1 |
| PULPROG | zgpg30 |
| TD | 65536 |
| SOLVENT | MeOD |
| NS | 91 |
| DS | 0 |
| SWH | 22727.273 Hz |
| FIDRES | 0.346791 Hz |
| AQ | 1.4417920 sec |
| RG | 2050 |
| DW | 22.000 usec |
| DE | 6.00 usec |
| TE | 300.0 K |
| D1 | 2.00000000 sec |
| d11 | 0.03000000 sec |
| DELTA | 1.89999998 sec |
| TD0 | 1 |
| ======== CHANNEL f1 ======== | |
| NUC1 | 13C |
| P1 | 9.70 usec |
| PL1 | -0.50 dB |
| SFO1 | 100.6288660 MHz |
| ======== CHANNEL f2 ======== | |
| CPDPRG[2 | waltz16 |
| NUC2 | 1H |
| PCPD2 | 90.00 usec |
| PL2 | -2.40 dB |
| PL12 | 15.10 dB |
| PL13 | 18.10 dB |
| SFO2 | 400.1516010 MHz |
| F2 - Processing parameters | |
| SI | 32768 |
| SF | 100.6176598 MHz |
| WDW | EM |
| SSB | 0 |
| LB | 3.00 Hz |
| GB | 0 |
| PC | 1.00 |

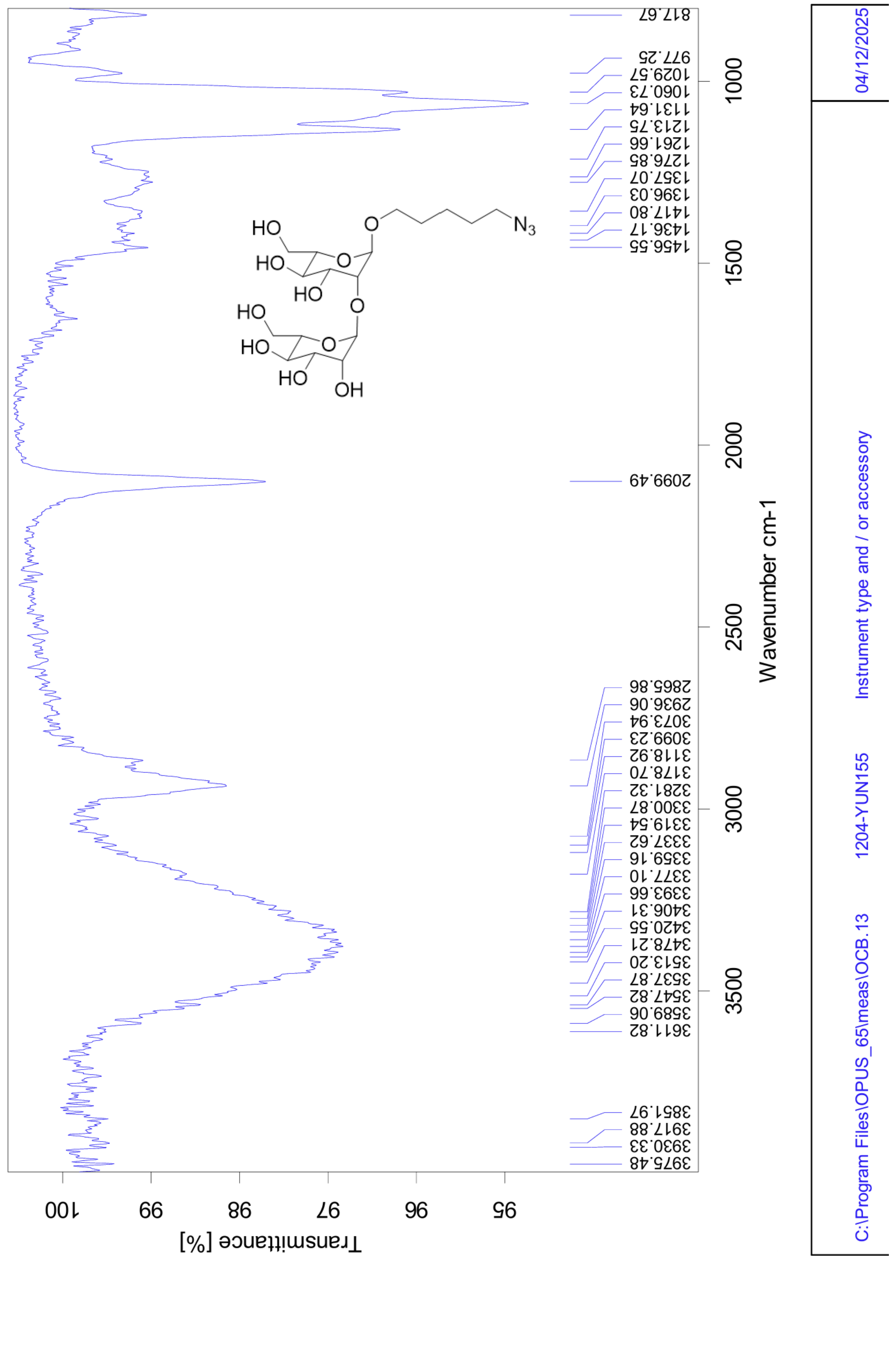
Transmittance [%]
100
99
98
97
96
95
Wavenumber cm-1
3500
3000
2500
2000
1500
1000
3975.48
3930.33
3917.88
3851.97
3611.82
3589.06
3547.82
3537.87
3513.20
3478.21
3420.55
3406.31
3393.66
3377.10
3359.16
3337.62
3319.54
3300.87
3281.32
3178.70
3118.92
3099.23
3073.94
2936.06
2865.86
2099.49
1456.55
1436.17
1417.80
1396.03
1357.07
1276.85
1261.66
1213.75
1131.64
1060.73
1029.57
977.25
817.67
HO
HO
HO
O
O
O
HO
HO
HO
O
OH
$N_3$
C:\Program Files\OPUS_65\meas\OCB.13
1204-YUN155
Instrument type and / or accessory
04/12/2025

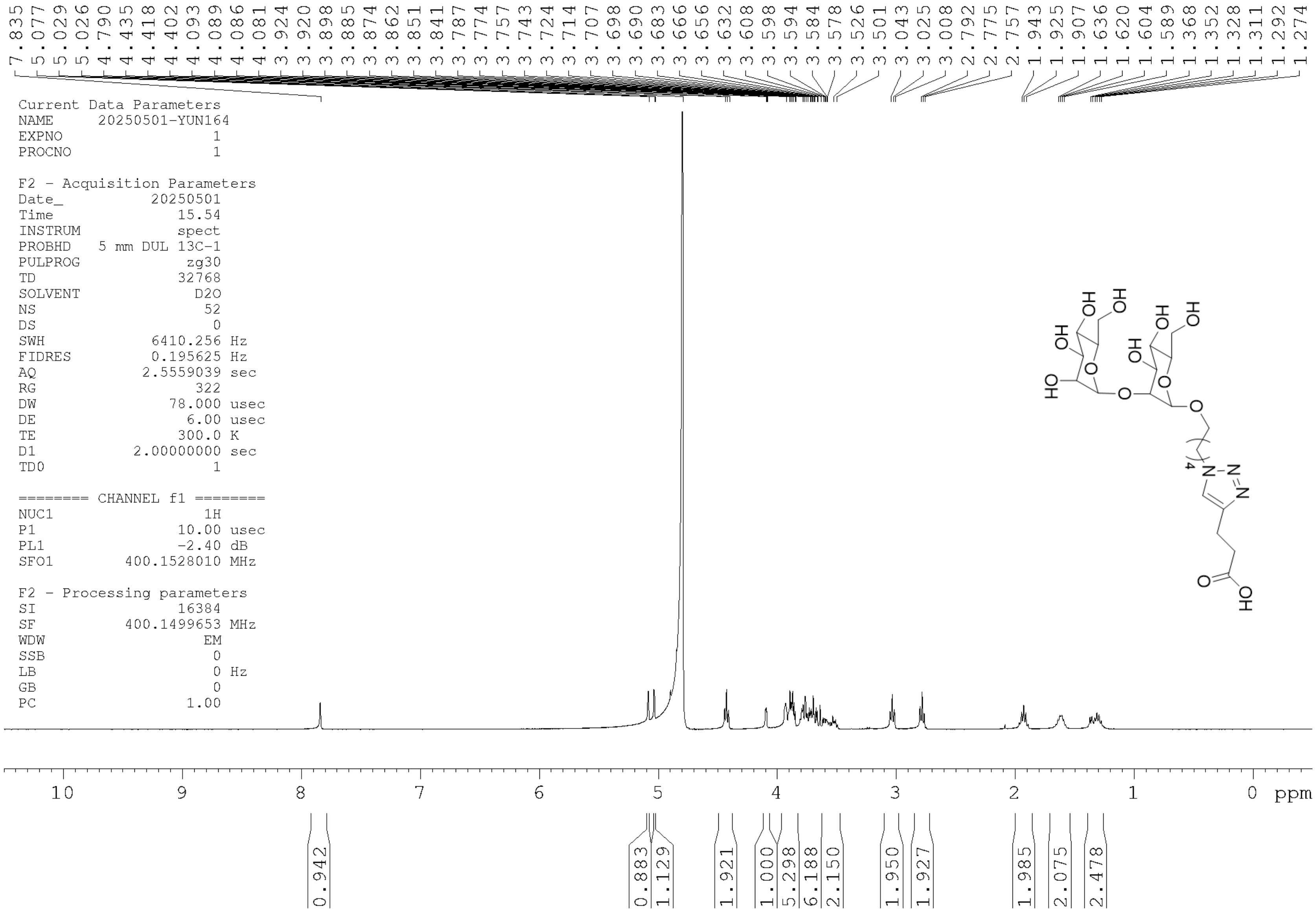
7.835
5.077
5.029
5.026
4.790
4.435
4.418
4.402
4.093
4.089
4.086
4.081
3.924
3.920
3.898
3.885
3.874
3.862
3.851
3.841
3.787
3.774
3.757
3.743
3.724
3.714
3.707
3.698
3.690
3.683
3.666
3.656
3.632
3.608
3.598
3.594
3.584
3.578
3.526
3.501
3.043
3.025
3.008
2.792
2.775
2.757
1.943
1.925
1.907
1.636
1.620
1.604
1.589
1.368
1.352
1.328
1.311
1.292
1.274
Current Data Parameters
NAME 20250501-YUN164
EXPNO 1
PROCNO 1
F2 - Acquisition Parameters
Date_ 20250501
Time 15.54
INSTRUM spect
PROBHD 5 mm DUL 13C-1
PULPROG zg30
TD 32768
SOLVENT D2O
NS 52
DS 0
SWH 6410.256 Hz
FIDRES 0.195625 Hz
AQ 2.5559039 sec
RG 322
DW 78.000 usec
DE 6.00 usec
TE 300.0 K
D1 2.00000000 sec
TD0 1
======== CHANNEL f1 ========
NUC1 1H
P1 10.00 usec
PL1 -2.40 dB
SFO1 400.1528010 MHz
F2 - Processing parameters
SI 16384
SF 400.1499653 MHz
WDW EM
SSB 0
LB 0 Hz
GB 0
PC 1.00
HO
HO
HO
OH
HO
HO
HO
HO
O
O
O
O
4
N
N
N
O
OH
10
9
8
7
6
5
4
3
2
1
0
ppm
0.942
0.883
1.129
1.921
1.000
5.298
6.188
2.150
1.950
1.927
1.985
2.075
2.478

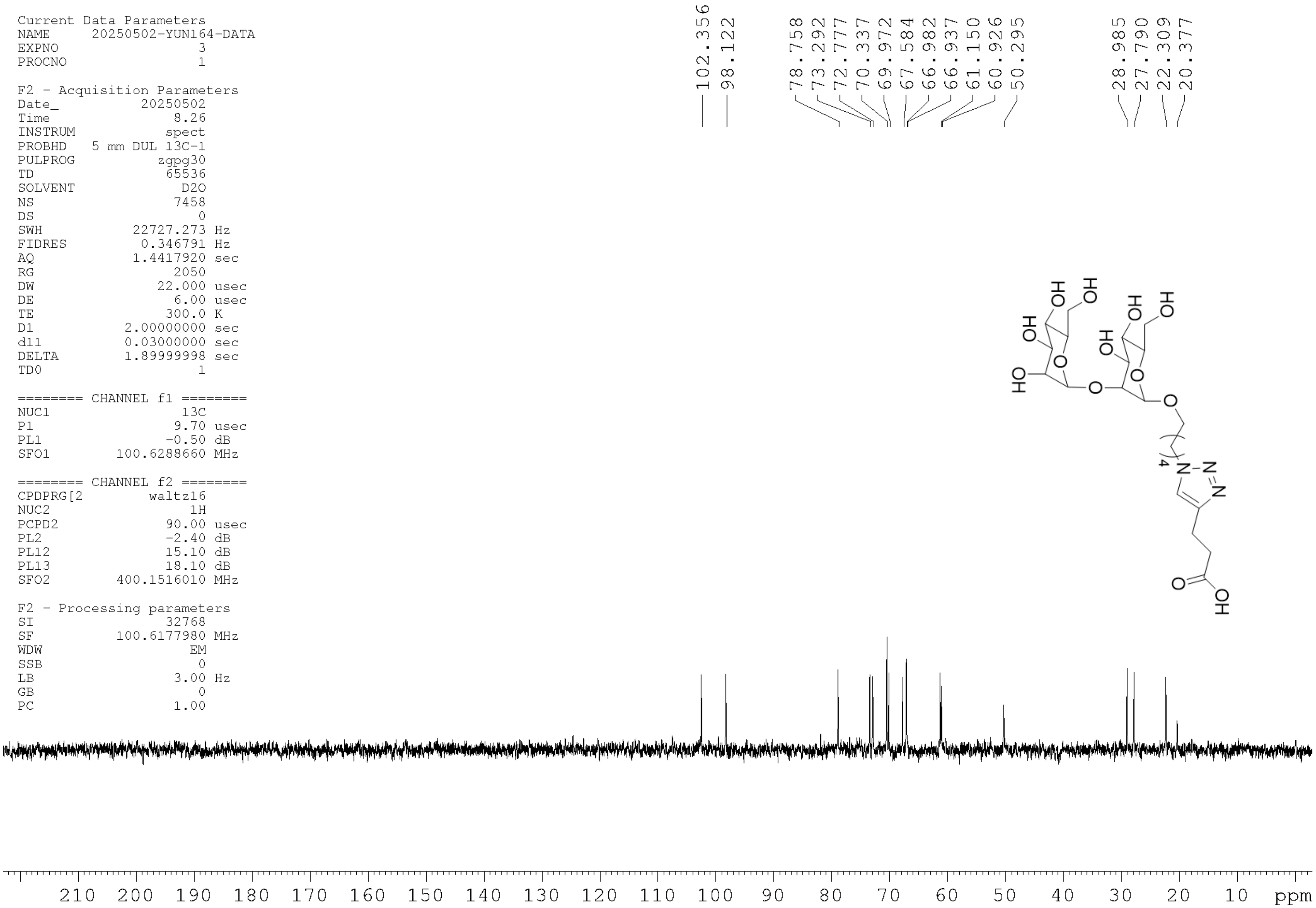
Current Data Parameters
NAME 20250502-YUN164-DATA
EXPNO 3
PROCNO 1

F2 - Acquisition Parameters
Date_ 20250502
Time 8.26
INSTRUM spect
PROBHD 5 mm DUL 13C-1
PULPROG zgpg30
TD 65536
SOLVENT D2O
NS 7458
DS 0
SWH 22727.273 Hz
FIDRES 0.346791 Hz
AQ 1.4417920 sec
RG 2050
DW 22.000 usec
DE 6.00 usec
TE 300.0 K
D1 2.00000000 sec
d11 0.03000000 sec
DELTA 1.89999998 sec
TD0 1

======== CHANNEL f1 ========
NUC1 13C
P1 9.70 usec
PL1 -0.50 dB
SFO1 100.6288660 MHz

======== CHANNEL f2 ========
CPDPRG[2 waltz16
NUC2 1H
PCPD2 90.00 usec
PL2 -2.40 dB
PL12 15.10 dB
PL13 18.10 dB
SFO2 400.1516010 MHz

F2 - Processing parameters
SI 32768
SF 100.6177980 MHz
WDW EM
SSB 0
LB 3.00 Hz
GB 0
PC 1.00

102.356
98.122
78.758
73.292
72.777
70.337
69.972
67.584
66.982
66.937
61.150
60.926
50.295
28.985
27.790
22.309
20.377

HO
OH
O
N
4
210 200 190 180 170 160 150 140 130 120 110 100 90 80 70 60 50 40 30 20 10 ppm

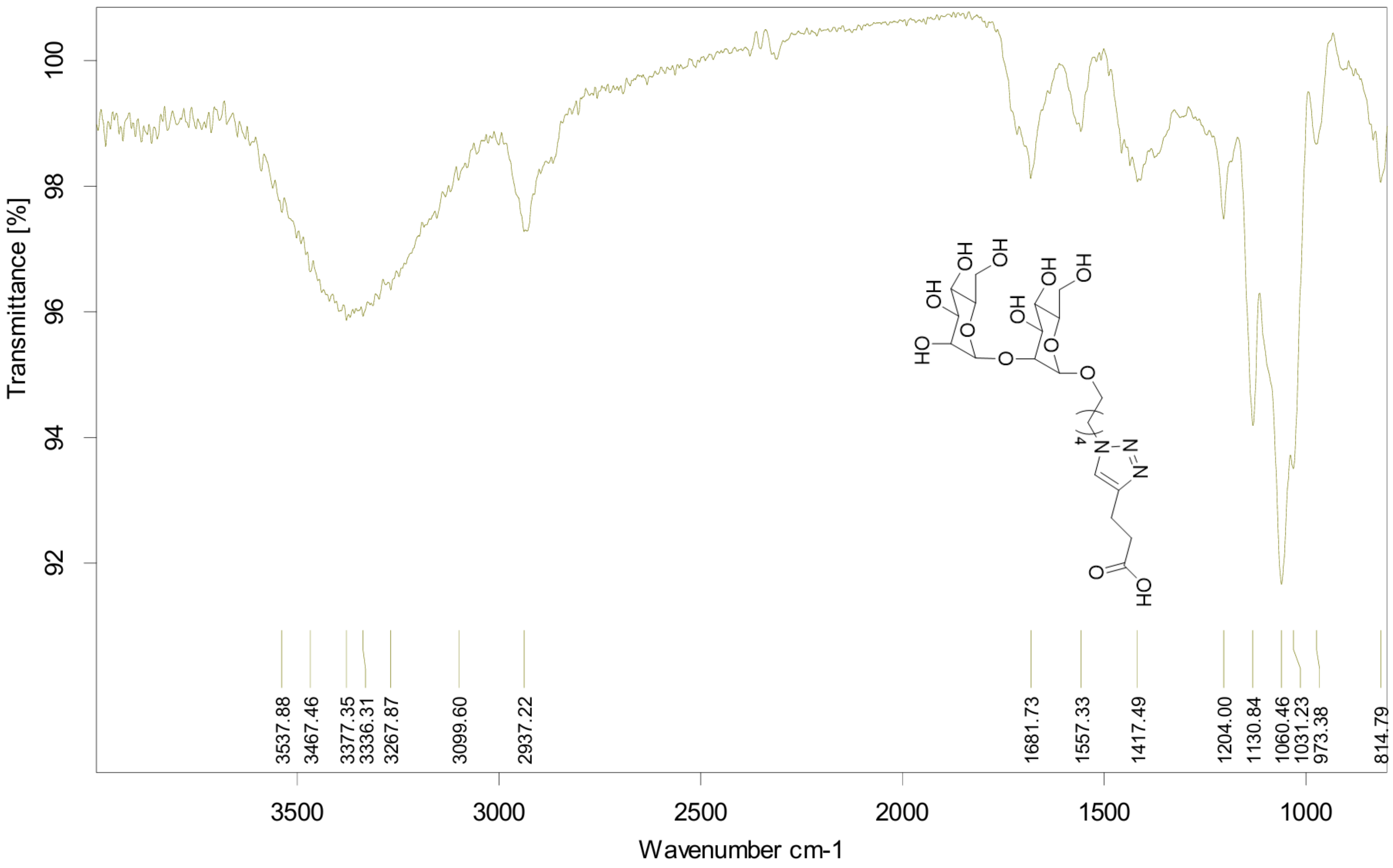

C:\Program Files\OPUS_65\meas\OCB.19 | 1204-YUN164 | Instrument type and / or accessory | 04/12/2025
